# BROOQS: Quartet-based Resolution of Level-1 Hybridization Cycles without Tests of Symmetry

**DOI:** 10.64898/2026.08.31.748319

**Authors:** Shayesteh Arasti, Siavash Mirarab

## Abstract

Modern phylogenomic analyses often seek to reconstruct both vertical and reticulate evolutionary histories. While the prevalence of non-vertical evolution is increasingly appreciated, inferring networks remains conceptually challenging and computationally demanding. Following the success of quartet-based methods for species tree inference despite gene tree discordance, several quartet-based network inference methods have been developed. Most of these quartet-based methods boil down to detecting asymmetry in minor quartet frequencies, often using statistical tests to control for noise. However, these approaches have not yet been amenable to dynamic programming algorithms used in species tree inference for search, nor have they allowed efficient calculation of statistics across all quartets without listing all quartets. Instead, existing methods either enumerate all quartets, losing some scalability, or subsample them, losing information. For search, an effective recently developed strategy for level-1 networks is to first build a multifurcating tree called a tree-of-blobs and then resolve each polytomy into a cycle. This two-step approach makes the problem easier both conceptually and computationally. However, resolving blobs still requires either subsampling quartets or sacrificing scalability. We introduce BROOQS, a quartet-based method for resolving trees of blobs into a level-1 phylogenetic network. BROOQS efficiently aggregates information from all quartets around a blob in quadratic time and builds a pairwise similarity matrix that we prove encodes the cyclic ordering around the blob without relying on individual quartet symmetry tests. To recover the order from the matrix, we use two approaches. BROOQS-PQtree uses an existing combinatorial method, which allows us to prove statistical consistency under the network multi-species coalescent (NMSC) model for blobs of size 5 to 15. BROOQS-Spectral uses a spectral algorithm, which we conjecture (but have not proved) is consistent in similar conditions based on our simulations, and shows better robustness to noise than BROOQS-PQtree. Across simulated and empirical datasets, BROOQS-spectral consistently improves accuracy and scalability compared to existing methods and extends to thousands of taxa.

## 1 Introduction

The recognition that phylogenetic networks are often better representations of evolutionary history than trees has increased in recent years, and the need for better methods has grown as access to higher-quality genomic data has accelerated (Bjornson et al., 2024; Kong et al., 2025). Inference of phylogenetic networks, however, has been computationally challenging for two reasons. First, the space of phylogenetic networks is substantially larger than the space of trees on the same set of taxa, making the search for the optimal network difficult (Wen et al., 2018). Second, evaluating a fixed network can itself be computationally expensive. For network inference to be credible, it needs to account for incomplete lineage sorting (ILS), which is possible under the network multi-species coalescent (NMSC) model. However, under NMSC, likelihood calculations must account for possible gene tree histories within the network, making full-likelihood calculations even more expensive than for trees (Zhu and Nakhleh, 2018). Maximum-likelihood and Bayesian methods have been developed to infer networks under the NMSC (Yu and Nakhleh, 2015; Wen et al., 2018), but their computational requirements have historically limited their application to relatively small datasets (Hejase and Liu, 2016; Zhu and Nakhleh, 2018). These limitations have motivated the development of pseudolikelihood and other approximations that moderately improve scalability (Yu and Nakhleh, 2015; Solís-Lemus and Ané, 2016).

Even for species trees, calculating likelihood under the multi-species coalescent (MSC) model of ILS is computationally expensive (Wu, 2012). This realization has pushed the field towards a scalable approach that first estimates gene trees and then summarizes them to infer a species tree (Mirarab et al., 2021). Quartet-based methods such as ASTRAL seek the species tree that maximizes the number of gene-tree quartets with which it agrees (Mirarab et al., 2014). The use of quartets is attractive because quartet-based scores can be statistically consistent under the MSC. Moreover, although there are 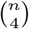 quartets on *n* taxa, ASTRAL computes and optimizes the aggregate quartet scores in roughly quadratic time (i.e., without explicitly enumerating all quartets), easily scaling to thousands of taxa and genes (Zhang et al., 2025). However, this approach does not directly extend to phylogenetic networks. The main theorem underlying these methods (that the species tree quartet has higher quartet frequency than the two alternatives (Allman et al., 2011)) does not hold for networks (Solís-Lemus et al., 2016). Under the NMSC, quartet frequencies are affected by both ILS and reticulation, with many scenarios of non-identifiable network topologies documented in the literature (Baños, 2019; Allman et al., 2024a). Nevertheless, the scalability of quartet-based methods has motivated many researchers to examine if quartets can be used for scalable network inference (Solís-Lemus and Ané, 2016; Rhodes et al., 2021).

Quartet-based network inference methods started with likelihood-based inference. For example, SNaQ uses quartet concordance factors (CF) to define a pseudolikelihood and searches for the network that maximizes this pseudolikelihood (Solís-Lemus and Ané, 2016). More recent methods are instead built on a key observation about CF asymmetry: Under the NMSC, a quartet of taxa without relevant reticulation events (i.e., a tree quartet) is expected to have one dominant quartet topology and two equally frequent alternative topologies, whereas cycle-inducing quartets can break the symmetry of alternative CFs (Fig. S1). Thus, many methods use statistical tests of asymmetry on quartet CFs to distinguish tree-like and network quartets (Mitchell et al., 2019; Allman et al., 2019).

To further reduce conceptual complexity and computational demand, a two-step approach has recently emerged (Allman et al., 2022, 2024b, 2025). Rather than constructing the entire network from scratch, these methods first infer a species tree and then contract edges that show evidence of reticulation. The resulting “tree of blobs” preserves the tree-like part of the network while representing each non-tree-like region as a multifurcation, which can subsequently be resolved into a cycle (Allman et al., 2022, 2024b). Recent theoretical results show that the tree of blobs is identifiable under the NMSC and can be consistently inferred from quartet frequencies (Allman et al., 2022, 2024b). This separates network inference into two smaller problems: inferring the tree of blobs, as in TINNIK (Allman et al., 2024b) and TOB-QMC (Dai et al., 2026), and resolving the resulting tree of blobs into a phylogenetic network, as in NANUQ and NANUQ+ (Allman et al., 2019, 2025) and NetCS (Dai and Molloy, 2026).

Although promising, the current two-step methods have two shortcomings. First, they either consider all 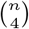 quartets, which limits their scalability (e.g. NANUQ and NANUQ+), or sample a subset of quartets and therefore do not use all the available information (e.g. NetCS). In contrast, quartet-based methods for species tree inference, such as ASTRAL, aggregate all quartets in roughly quadratic time (i.e., do not explicitly enumerate quartets), allowing them to scale to thousands of taxa. Such methods have not yet been developed for network inference because their underlying dynamic programming algorithm does not trivially extend to statistical tests used in network inference. Second, the statistical tests of asymmetry used by these methods can be misled. Various evolutionary and systematic processes, other than reticulations, can violate the expected symmetry in individual quartets. Strong selection, long-branch attraction, rate heterogeneity, and compositional bias are among these processes that can produce asymmetric and conflicting signals (He et al., 2020; Adams et al., 2018; Felsenstein, 1978; He et al., 2026; Kück et al., 2022; Jermiin et al., 2004; Frankel and Ané, 2023). Moreover, these tests are sensitive to the statistical thresholds used to determine whether a quartet is classified as tree-like or network-like. Because the power of these statistical tests depends on the number of sampled quartets and gene trees, there is no globally optimal choice of these thresholds, making it challenging to tune them.

In this paper, we develop a new set of algorithms for resolving the tree of blobs into a level-1 network, seeking to address shortcomings of existing methods. Instead of relying on individual quartet tests, we introduce a similarity measure between every pair of components around a blob that aggregates the signal from all quartets involving the two components and encodes the cyclic ordering around it. We develop an efficient quadratic time dynamic programming algorithm to compute this measure without explicitly examining all quartets or subsampling. We show the matrix is a strict Robinson matrix (Hubert et al., 1998) for blobs of size 5 to 15, which enables us to use existing *seriation* algorithms to recover the cyclic ordering of the components and resolve the blob into a level-1 cycle. We explore two seriation methods: a combinatorial method that provides guarantees of correctness, and a spectral algorithm, which we also conjecture is correct under the same conditions. We show that our method, Blob Resolution by Optimal Ordering of Quartet-based Similarities (BROOQS), drastically outperforms alternative methods when the true tree of blobs is given, and is consistently among the most accurate methods when the tree of blobs is estimated. We further show that BROOQS is highly scalable and memory-efficient, extending to datasets with thousands of taxa. Finally, we use BROOQS to study debated relationships in avian and plant datasets with approximately 60,000 gene trees and 1,000 taxa, respectively, demonstrating its ability to recover reticulation patterns in large empirical datasets.

## 2 Materials and Methods

### 2.1 Preliminaries

#### 2.1.1 Notation and Background

##### Networks

A binary *directed* phylogenetic network *N* ^+^ = (*V, E*) is a directed acyclic graph defined with a set of vertices *V* and a set of edges *E* with a unique *root r* ∈ *V* with in-degree zero and out-degree two. The leafset of *N* ^+^ is the set of all vertices with out-degree zero and in-degree one and is denoted by *L*. The rest of the nodes in *V* are divided into two sets: *tree* nodes *V*_*T*_ with in-degree one and out-degree two, and *hybrid* (*a*.*k*.*a. reticulation*) nodes *V*_*H*_ with in-degree two and out-degree one. Edges incoming to hybrid nodes are called *hybrid* (or *reticulation*) edges and the rest are called *tree edges*. Although *N* ^+^ is acyclic, throughout the paper, we define an undirected *cycle* as a closed path from a node *v ∈ V* to itself, ignoring the direction of the edges in *E*. A phylogenetic network *N* ^+^ is *level-1* if no two cycles in *N* ^+^ share an edge (Huson et al., 2010); in a level-1 network, every cycle contains exactly one hybrid node. The size of a cycle is the number of edges it contains. We refer to a cycle of size *d* as a *d*-cycle. A semi-directed network *N* can be obtained from *N* ^+^ by removing the direction of all tree edges, and suppressing all resulting degree-two nodes (Fig. 1a).

**Figure 1.**
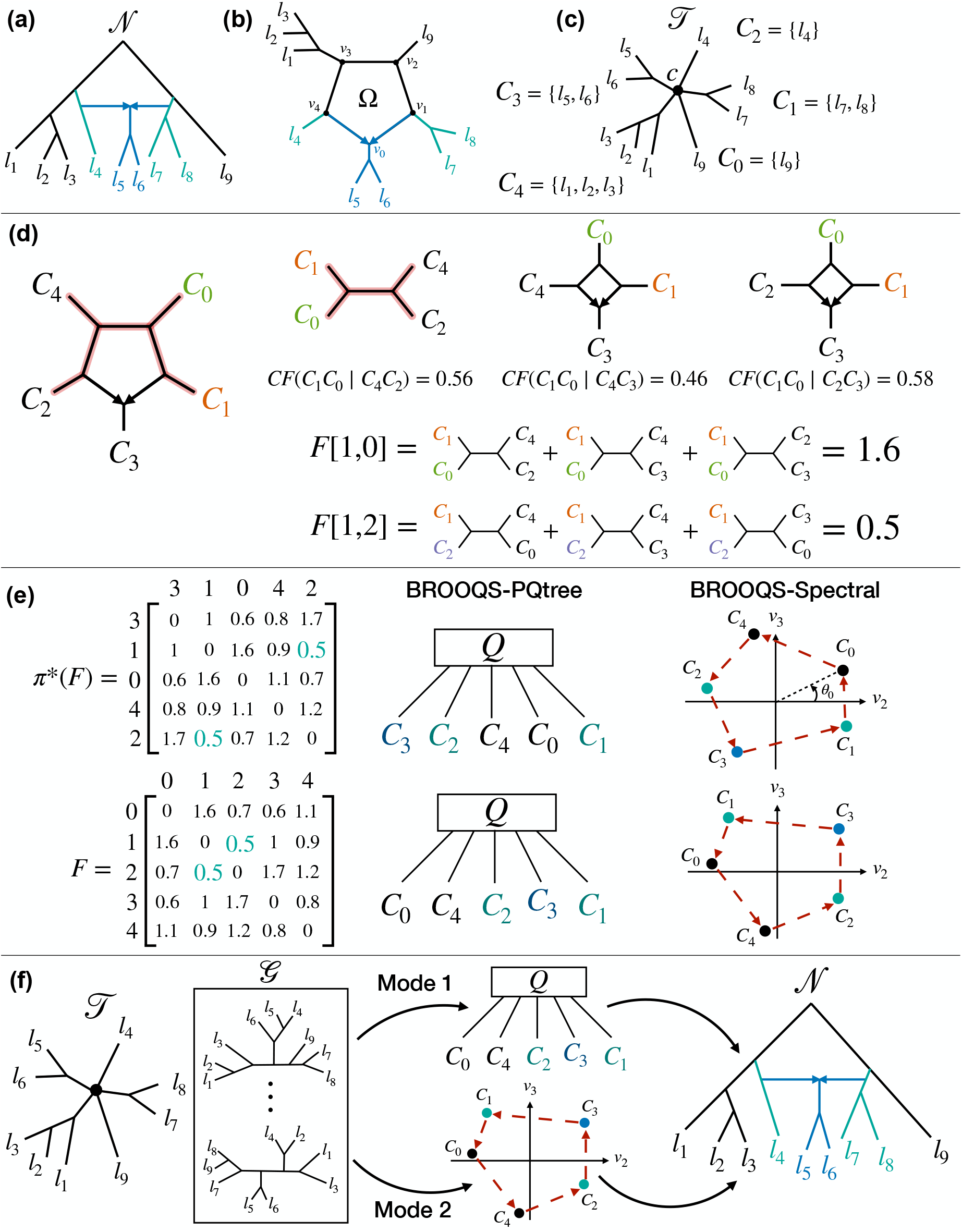
An overview of BROOQS. **(a)** A semi-directed network *N* on a set of taxa *L* = *{l*_1_, *l*_2_,…, *l*_9_*}*, with one reticulation event. **(b)** The 5-cycle Ω in *N* defined by the closed path [*v*_0_, *v*_1_,…, *v*_4_]. The hybrid component and its two neighboring components are highlighted. **(c)** Contracting the cycle Ω into a 5-blob results in the tree of blobs *T* and a polytomy *c* with degree 5. Removing *c* from *T* results in the partition *C* = *{C*_0_, *C*_1_,…, *C*_4_*}* on *L*. Finding the hybrid component (*C*_3_) and the right order *ω&*\*(*C*) = [*C*_3_, *C*_1_, *C*_0_, *C*_4_, *C*_2_] recovers Ω. **(d)** The F-matrix *F* (Eq. (1)) is a symmetric similarity matrix that sums the probability of a pair of components being grouped together in a quartet, over all pairs of the remaining components. All the summands of *F* [1, 0] and *F* [1, 2] are shown. See Figure S1 for calculation of concordance factors. Note that *F* [1, 0] *> F* [1, 2] because *C*_1_, *C*_2_ are further from each other than *C*_1_, *C*_0_. **(e)** The cyclic order is recovered from *F* in two ways: BROOQS-PQtree, using the PQ-tree algorithm of Armstrong et al. (2021), and BROOQS-Spectral, which sorts components by their angles in the spectral embedding of Coifman et al. (2008). Both are invariant to the permutation of *F* ; shown are an arbitrary permutation and the true one, *π*\*(*F* ), which is circular Robinson (Definition 1). The minimum non-diagonal entry of *F* marks the two neighbors of the hybrid component. **(f)** The BROOQS approach. The input is a tree of blobs *T* as well as a set of gene trees *G* evolved on *N* . BROOQS constructs the level-1 networks by: 1) Computing the empirical F-matrix 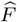, 2) recovering the correct cyclic ordering of the components using either BROOQS-PQtree (Mode 1) or BROOQS-Spectral (Mode 2), and 3) identifying the hybrid component.

A *metric* phylogenetic network with *H* hybrid nodes is a tuple (*N, ω, γ*), where *ω* ∈ ℝ^+|*E*|^ represents edge lengths and each *γ* ∈ (0, 1)^*H*^ represents *inheritance probabilities*, such that for a hybrid node *h*, the right incoming edge has probability γ_*h*_ and other 1 − γ_*h*_ (Baños, 2019). We omit the subscript when clear. In this paper, we focus on level-1 metric networks.

##### Quartets/quarnets

A quartet is an unrooted binary phylogenetic tree on four taxa. For any four taxa *a, b, c, d*, there are exactly three distinct quartet topologies, denoted by *ab*|*cd, ac*|*bd*, and *ad*|*bc*, where the quartet *ab*|*cd* is the topology in which the internal edge separates taxa *a, b* from *c, d*. A phylogenetic tree *T* displays a quartet on *a, b, c, d* if the tree induced on these four taxa (hereby referred to as *T* |_{*a,b,c,d*}_) has the same topology as the quartet. For a subset of taxa *L*^’^ ⊆ *L*, the induced phylogenetic network *N* on *L*^’^, also referred to as 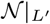, is defined as in Baños (2019). In particular, it is obtained by restricting the network to *L*^’^ and simplifying the resulting subnetwork by suppressing degree-two nodes and removing 2-cycles (Fig. S1). An induced network on four taxa is called a *quarnet* (Huber et al., 2018). If the original network is metric, the induced network is called a *metric quarnet*.

##### Tree of Blobs

An edge is a *cut edge* if its removal creates two disconnected components in *N*, and is called a *non-cut edge* otherwise. Every hybrid edge in *N* is also a non-cut edge. A *blob* is a maximal connected subnetwork of *N* with no cut edges (Allman et al., 2022). For a level-1 network, each blob corresponds to a single cycle (Fig. 1b), and we define the degree of the blob by its number of incident cut edges, referring to a blob with degree *d* as a *d*-blob. In this definition, every leaf *l* ∈ *L* and every tree node *v* ∈ *V*_*T*_ form trivial blobs of degree 1 and 3. Henceforth, we use the term blob to denote non-trivial blobs. A *tree-of-blob T* = (*V*_*T*_, *E*_*T*_ ) of a network *N* is a tree obtained by contracting all non-cut edges in *N*, and removing the direction of the cut edges to obtain an unrooted tree (Gusfield et al., 2007). Each blob in *N* with degree *d* is represented with a single degree-*d* node in the tree of blobs (Fig. 1c).

##### Network Multi-Species Coalescent (NMSC) Model

We assume gene trees are sampled from NMSC, an extension of the multi-species coalescent model (MSC) to phylogenetic networks (Nakhleh, 2011; Yu et al., 2012). Under the NMSC model, individual copies of genes evolve and coalesce along a metric phylogenetic network. At each hybrid node, a lineage chooses one of the two incoming hybrid edges according to their inheritance probabilities. These inheritance choices, together with stochastic coalescent events along the branches of the network, result in a rooted binary phylogenetic tree called a *gene tree*. Under the NMSC model, the distribution of gene trees on four taxa depends only on the induced metric quarnet (Baños, 2019). The probabilities of the three quartet topologies are called the quartet concordance factors (CFs), denoted by (*CF*(*ab*|*cd*),*CF*(*ac*|*bd*),*CF*(*ad*|*bc*)), and are shown in Figure S1.

#### 2.1.2 Problem statement: resolving tree of blobs

Resolving a level-1 tree of blobs *T* is the process of replacing every node *c* ∈ *V*_*T*_ of degree *d >* 3 with a *d*-cycle. The resolution replaces *c* and its incident edges with one hybrid node and *d −* 1 tree nodes, adding two hybrid edges from two of the new tree nodes to the hybrid node, adding *d* − 2 non-cut edges connecting the remaining tree nodes, and reconnecting the resulting *d* nodes to the *d* connected components created by removing *c*. The result will be a connected level-1 network by construction.

Let the node *c* of *T* correspond to a degree-*d* cycle Ω in *N*, represented by the closed path [*v*_*h*_, *v*_1_, *v*_2_,…, *v*_*d*−1_, *v*_*h*_], where *v*_*h*_ ∈ *V*_*H*_ is the hybrid node and every pair of consecutive nodes is adjacent in Ω. Removing *c* and its incident edges from *T* creates *d* connected components (Fig. 1c). We identify each connected component with its leaf set and denote the resulting partition of *L* by *C* = *{C*_0_, *C*_1_,…, *C*_*d*−1_*}*. Each component *C*_*i*_ ∈ *C* corresponds uniquely to a node *v* on Ω (Fig. 1b).

A cyclic ordering of *C* is defined by a sequence *θ*(*C*) = [*C*_*θ*(0)_, *C*_*θ*(1)_,…, *C*_*θ*(*d*−1)_], where (*θ*(0),. .., *θ*(*d* − 1)) is a permutation of (0,…, *d* − 1). The traversal of Ω induces the *true ordering* 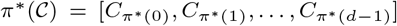, where 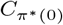 corresponds to the hybrid node *v*_*h*_, and for 1 ≤ *i* ≤ *d* − 1, 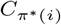 corresponds to *v*_*i*_. Thus, We call 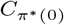 the *hybrid component*, and 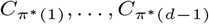 the *non-hybrid components*. For an ordering *θ*(*C*) and an index 0 ≤ *r* ≤ *d* − 1, we define its rotation by Rot(*θ*(*C*), *r*) = [*C*_*θ*(*r*)_, *C*_*θ*((*r*+1) mod *d*)_,…, *C*_*θ*((*r*+*d*−1) mod *d*)_], and its reversal by Rev(*θ*(*C*)) = [*C*_*θ*(0)_, *C*_*θ*(*d*−1)_, *C*_*θ*(*d*−2)_,…, *C*_*θ*(1)_]. Since cyclic orderings have no distinguished starting point or orientation, we consider all rotations of *π*\*(*C*) and all rotations of Rev(*π*\*(*C*)) to be equivalent to *π*\*(*C*). Resolving a blob in *T* in effect involves two questions: finding one of the equivalent true cyclic orderings of the components *C*, and finding the hybrid component 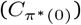.

##### Tree of blobs resolution problem

Let (*N* ^+^, *ω, γ*) be a metric level-1 rooted phylogenetic network, and let *G* = *G*_1_, *G*_2_,…, *G*_*m*_ be a set of *m* gene trees evolved on (*N* ^+^, *ω, γ*) under the NMSC model. Given the gene trees *G* and the tree of blobs *T*, our task is to reconstruct the correct semi-directed level-1 network *N* .

### 2.2 BROOQS Algorithm

#### 2.2.1 Theoretical foundations

We present a method, called BROOQS, for the tree of blobs resolution problem. Our method relies on the theory of circular seriation from a similarity matrix. Before stating our main results, we define a class of matrices first introduced by Hubert et al. (1998) and redefined by Armstrong et al. (2021) that enables theoretical guarantees for our method:

##### Definition 1

((*strict*) Circular Robinson similarity matrix) Let *S* ∈ R^*d* × *d*^ be a symmetric similarity matrix. The matrix *S* is called a *(strict) circular Robinson similarity matrix* if every row is *(strict) circularly unimodal*; that is, for every 1≤ *i* ≤ d, the cyclic sequence [*S*_*i, i* +1_,…, *S*_*i, d*_, *S*_*i*, 1_,…, *S*_*i, i* − 1_}] is *(strictly) decreasing* up to a minimum and *(strictly) increasing* thereafter, where the minimum may be attained at at most two consecutive positions (Armstrong et al., 2021).

Our main contribution is introducing a similarity matrix between the set of components *C* of the cycle Ω, which we refer to as the *F-matrix F* ∈ ℝ^*d* × *d*^. We prove that the true permutation of the F-matrix π* (*F*) is a strict circular Robinson similarity matrix on *C* based on the cyclic ordering of Ω, which can be efficiently computed from gene trees. We show two approaches to recover the true cyclic ordering π* (*C*) from the matrix *F*, one based on the PQ-tree algorithm by Armstrong et al. (2021), and a spectral method based on that of Coifman et al. (2008). We prove statistical consistency for the former for 4 < *d* < 16, and conjecture that the latter also recovers the true ordering for 4 ≤ *d* < 20 based on extensive simulations (no proofs).

If *C*_*i*_, *C*_*j*_, *C*_*k*_, *C*_*l*_ ∈ *C* are four distinct components, then any choice of taxa *t*_*i*_ ∈ *C*_*i*_, *t*_*j*_ ∈ *C*_*j*_, *t*_*k*_ ∈ *C*_*k*_, and *t*_*l*_ ∈ *C*_*l*_ induces the same metric quarnet (Baños, 2019). Consequently, the quartet CFs are independent of the particular choice of taxa.

##### Definition 2

(Component Concordance Factors) For any four distinct components *C*_*i*_, *C*_*j*_, *C*_*k*_, *C*_*l*_ ∈ *C*, we define the *component concordance factors (CF)* (*CF*(*C*_*i*_*C*_*j*_ | *C*_*k*_*C*_*l*_),*CF*(*C*_*i*_*C*_*k*_ | *C*_*j*_ *C*_*l*_),*CF*(*C*_*i*_*C*_*l*_ | *C*_*j*_ *C*_*k*_)) to be the quartet CFs of the metric quarnet induced by any choice of taxa *t*_*i*_ ∈ *C*_*i*_, *t*_*j*_ ∈ *C*_*j*_, *t*_*k*_ ∈ *C*_*k*_, *t*_*l*_ ∈ *C*_*l*_, which is independent of the particular choice of taxa *t*_*i*_, *t*_*j*_, *t*_*k*_, *t*_*l*_. A metric quarnet induced on four components is a *tree quarnet* if *C*_*i*_, *C*_*j*_, *C*_*k*_, *C*_*l*_ are all non-hybrid components, and a *network quarnet* otherwise (Baños, 2019, also see Fig. S1).

As one may expect (and we prove), two components that are adjacent on the cycle tend to appear on the same side of a quartet more frequently than those that are further apart. This observation motivates the following definition of the F-matrix.

##### Definition 3

(F-Matrix) Let Ω be a *d*-cycle with components *C* = {*C*_0_, *C*_1_,…, *C*_*d*−1_}. The *F-matrix* on *C* is a symmetric matrix *F* ∈ ℝ^*d*×*d*^ whose rows and columns are indexed by [0, 1,…,*d* − 1] and whose entries are given by

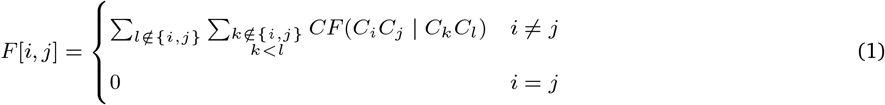

Intuitively, the F-score measures how strongly two components are associated by summing the probabilities that taxa from those two components are grouped together in a quartet over all pairs of the remaining components (Fig. 1d).

The following theorem (proved in Appendix Section B.1) is the main result that is the basis of our method:

##### Theorem 1

Let Ω be a *d*-cycle where 4 ≤ *d* ≤ 15 with components *C* and true cyclic ordering π* (*C*). Then, assuming NMSC, the permuted F-matrix π* (*F*) is a strict circular Robinson similarity matrix.

Let 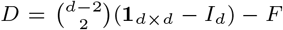 denote the pairwise *dissimilarity* matrix with zero diagonal. By definition, 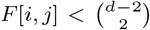 for any pair of components *C*_*i*_, *C*_*j*_ ; therefore, *D*[*i, j*] > 0 for *i* ≠ *j* and *D*[*i, j*] = 0, otherwise. It follows trivially that if *F* is a strict Robinson similarity matrix, *D* is a strict dissimilarity matrix, where each row and each column strictly increase to one or two consecutive maximum(s) and strictly decrease afterward. The following lemma (proved in Appendix Section B.4) is a consequence of the theory of PQ-tree algorithm for strict dissimilarity matrices by Armstrong et al. (2021).

##### Lemma 1

Let Ω be a *d*-cycle where 4 ≤ *d* ≤ 15 with components *C* and true cyclic ordering π* (*C*) with permuted F-matrix π* (*F*). Then, the PQ-tree algorithm of Armstrong et al. (2021) on the dissimilarity matrix 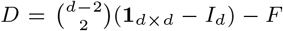 returns at most two inequivalent orderings П = {π_1_, π_2_}, such that 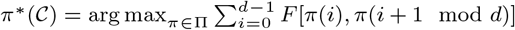.

While Lemma 1 provides the theoretical foundation that the true ordering π* (*C*) can be *uniquely* recovered from the PQ-tree algorithm of Armstrong et al. (2021), we introduce a second approach based on the spectral algorithm for circular seriation of Coifman et al. (2008). Given a symmetric similarity matrix *F*, let *S* = diag(*F* **1**_*d*_) be its degree matrix, and let *L* = *S* − *F* denote the corresponding graph Laplacian (von Luxburg, 2007). We state the following result as a conjecture.

##### Conjecture 1

(Circular spectral ordering for F-matrix) Let Ω be a *d*-cycle where 4 ≤ *d* ≤ 19 with the true cyclic ordering π* (*C*) and the F-matrix *F*. Then the two-dimensional spectral embedding of *F*, formed by the eigenvectors corresponding to the second- and third-smallest eigenvalues of its graph Laplacian *L*, embeds the rows of *F* along a closed curve that follows the cyclic ordering induced by π* (*C*).

The correctness of spectral embeddings has only been proved for a particular class of (strict) circular Robinson similarity matrices (Recanati et al., 2018), referred to as the *circulant* matrices. While our F-matrix is not circulant, we conjecture that the particular properties of the F-matrix imposed by the structure of the cycle Ω ensure that the spectral embeddings form a closed curve consistent with the true ordering π* (*C*). This conjecture is supported by our extensive simulations, where spectral methods recovered the true ordering for all cycles of size 5 ≤ *d* ≤ 19 in millions of replicates (see Section C.2).

##### Remark 1

Theorem 1 holds for any *d* ≥ 4 for the non-hybrid components. The hybrid component can break the Robinson property and thus Theorem 1 when *d* > 15 especially under extreme parameter settings (Fig. S2 shows a counterexample). Nevertheless, our empirical evaluation indicates that the PQ-tree algorithm recovers the correct cyclic ordering for such large cycles, including cases where the Robinson property is violated. Additionally, the spectral embedding recovers the true ordering for all cycles of size *d* ≤ 19, including cases where the Robinson property is violated. However, it fails on some larger cycles, especially for a particular selection of cycle parameters. See Section C.2 and Table S3) for our empirical analysis.

Once the cyclic ordering is determined, we use a second theorem (Appendix Section B.2) to identify the hybrid component.

##### Theorem 2

Let Ω be a *d*-cycle in *N*, where *d* > 4, with true component ordering π * (*C*) = [*C*_*π\**_ (0), *C*_*π\**_ (1),…, *C*_*π\**_(*d*−1)], where *C*_*π\**_ (0) is the hybrid component and *C*_*π\**_ (1) and *C*_*π\**_(*d*−1) are the neighboring components to *C*_*π\**_ (0). Then *F*[π* (1), π* (*d* − 1)] < *F*[π* (*i*), π* (*j*)] for every unordered pair (*i, j*) ≠ (1,*d* − 1). Equivalently, *F*[π* (1), π* (*d* − 1)] = min_*i*≠*j*_ *F*[*i, j*], and this minimum is unique.

While these theorems and results are based on the true CFs, we can estimate the F-matrix from the data. Let G ∈ *G* be a gene tree and let *t*_*i*_ ∈ *C*_*i*_, *t*_*j*_ ∈ *C*_*j*_, *t*_*k*_ ∈ *C*_*k*_, and *t*_*l*_ ∈ *C*_*l*_. By definition of the component CF, 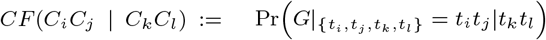. Therefore, we can estimate the component CF by

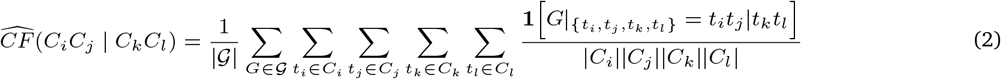

That is, 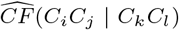 is the proportion of quartets that display the topology *t*_*i*_*t*_*j*_|*t*_*k*_*t*_*l*_ among all gene trees and all choices of taxa from each of *C*_*i*_, *C*_*j*_, *C*_*k*_, *C*_*l*_. Since the gene trees are sampled independently under the NMSC model, by the law of large numbers, 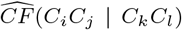 converges almost surely to *CF*(*C*_*i*_*C*_*j*_ | *C*_*k*_*C*_*l*_) for every choice of four distinct components. We can plug 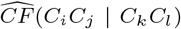 values into Equation (1) to obtain an estimated 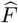 (we will show a trick to dramatically speed up calculation of 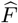). Since each F-score is a finite sum of component CFs, 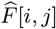 converges almost surely to *F*[*i, j*]. Therefore, every entry of 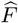 converges almost surely to the corresponding entry of *F*.

##### Proposition 1

As the number of gene trees *m* → ∞, the empirical F-matrix converges to the true F-matrix. That is, 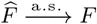.

##### Remark 2

Proposition 1 together with Theorem 2 and Lemma 1 prove the statistical consistency of the PQ-tree algorithm under the NMSC model given true gene trees and a true tree of blobs. Note that all the proofs used in this paper are based on inequalities between the elements of the F-matrix *F*. Let *ϵ* be the smallest of such differences. Then, as *m* → ∞, with arbitrarily high probability 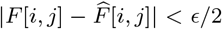 for *any* pair 0 ≤ *i* < *j* ≤ *d*, ensuring correct recovery of both the cyclic ordering and the hybrid component.

#### 2.2.2 BROOQS Algorithmic steps

Given a tree of blobs *T* and a set of gene trees *G*, BROOQS resolves each blob of *T* independently (Fig. 1f). Our algorithm (Algorithm 1) has three steps for each degree-*d* node c ∈ V_*T*_ corresponding to the components *C* = {*C*_0_, *C*_1_,…, *C*_*d*−1_}.

##### Step 1: Efficient calculation of 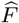.

Computing the empirical F-matrix naively from Equation (2) is computationally expensive.

Let *n* be the number of taxa in *L, m* the number of gene trees in *G*, and *d* the size of the cycle Ω. Assuming that the induced quartet G|_{*t,t,t,t* }_ can be obtained in *O*(*n*) time, computing a single empirical F-score 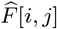 requires *O*(*md*^2^*n*^5^) time in the worst case. Consequently, done naively, computing all pairwise F-scores and constructing the empirical F-matrix 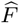 requires *O*(*md*^4^*n*^5^) time. We can do far better.

We use a dynamic programming approach that computes the empirical F-matrix in Θ(*md*^2^*n*) time. The algorithm essentially relies on a long-known technique (Bryant et al., 2000) that the ASTRAL family of tools have also adopted to compute quartet scores summed over all quartets without listing all quartets. During the computation of 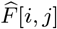 for the pair (*C*_*i*_, *C*_*j*_ ), rather than computing individual 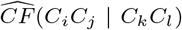 values for all possible choices of *C*_*k*_, *C*_*l*_, our dynamic programming approach directly computes 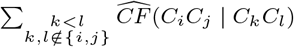 in *O*(*n*), and therefore, avoids unnecessary computation of intermediate values such as 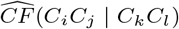. Algorithm S3 performs a one-time *O*(*nd*) preprocessing for each gene tree G ∈ *G*, during which it computes Θ(*d*) variables for each node. Then, it computes an intermediate matrix 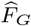 in *O*(*d*^2^*n*) time by a bottom-up traversal of G that performs a constant number of operations at each node (Algorithm S4). The empirical F-matrix is then obtained by averaging these matrices over all gene trees: 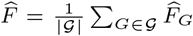. This reduces the total running time from *O*(*md*^4^*n*^5^) to *O*(*md*^2^*n*). See Appendix Section B.5 for all the details and implementation of this algorithm.

Our algorithm can handle missing data naturally as long as at least one taxon is observed from each component of the cycle Ω. The natural way to handle cases where gene trees miss entire components (i.e., not counting those in individual terms of 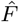) would break our dynamic programming. Instead, to avoid sacrificing running time, we use a heuristic approximation based on the assumption that the probability of a component being missing from a gene tree G is independent of the presence or absence of other components. Appendix Section B.5.1 provides a full description of this heuristic and Fig. S4 provides empirical evaluation, showing that it is highly effective.

##### Step 2: Recover the cyclic ordering

BROOQS has two modes of recovering the cyclic ordering of Ω: BROOQS-PQtree and BROOQS-Spectral (Fig. 1e). BROOQS-PQtree uses the PQ-tree algorithm by Armstrong et al. (2021) to iteratively combine the nearest components and order them based on their proximity on the F-matrix *F* (see Section B.3 for full description of the PQ-tree algorithm and Algorithm S2 for its implementation). This algorithm returns at most two inequivalent orderings that are consistent with *F*. We choose the ordering that maximizes the *tour score*; i.e. π* (*C*) = arg 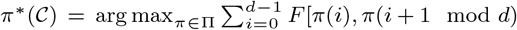. By Lemma 1, this approach correctly recovers the true cyclic ordering π* (*C*) for any *d*-cycle where 4 ≤ *d* ≤ 15. BROOQS-Spectral uses the spectral ordering algorithm of Coifman et al. (2008) to recover the cyclic ordering. Let *L* = *S* −*F* be the graph Laplacian of *F*, where *S* is the corresponding degree matrix. Let v_2_ and v_3_ denote the eigenvectors corresponding to the second- and third-smallest eigenvalues of *L*, respectively. Since the smallest eigenvalue of *L* is always 0, these are the first two non-trivial eigenvectors. By Conjecture 1, the vectors v_2_ and v_3_ define a two-dimensional spectral embedding whose cyclic ordering is identical to the ordering of the rows and columns of the underlying circular Robinson similarity matrix π* (*F*). For each component *C*_*i*_, we compute its polar angle *θ*_*i*_ = atan2(v_3_[*i*], v_2_[*i*]). We output the ordering π(*C*) obtained by sorting the components according to their projection angles (see Fig. 1e and Algorithm 1).

###### Algorithm 1

BROOQS

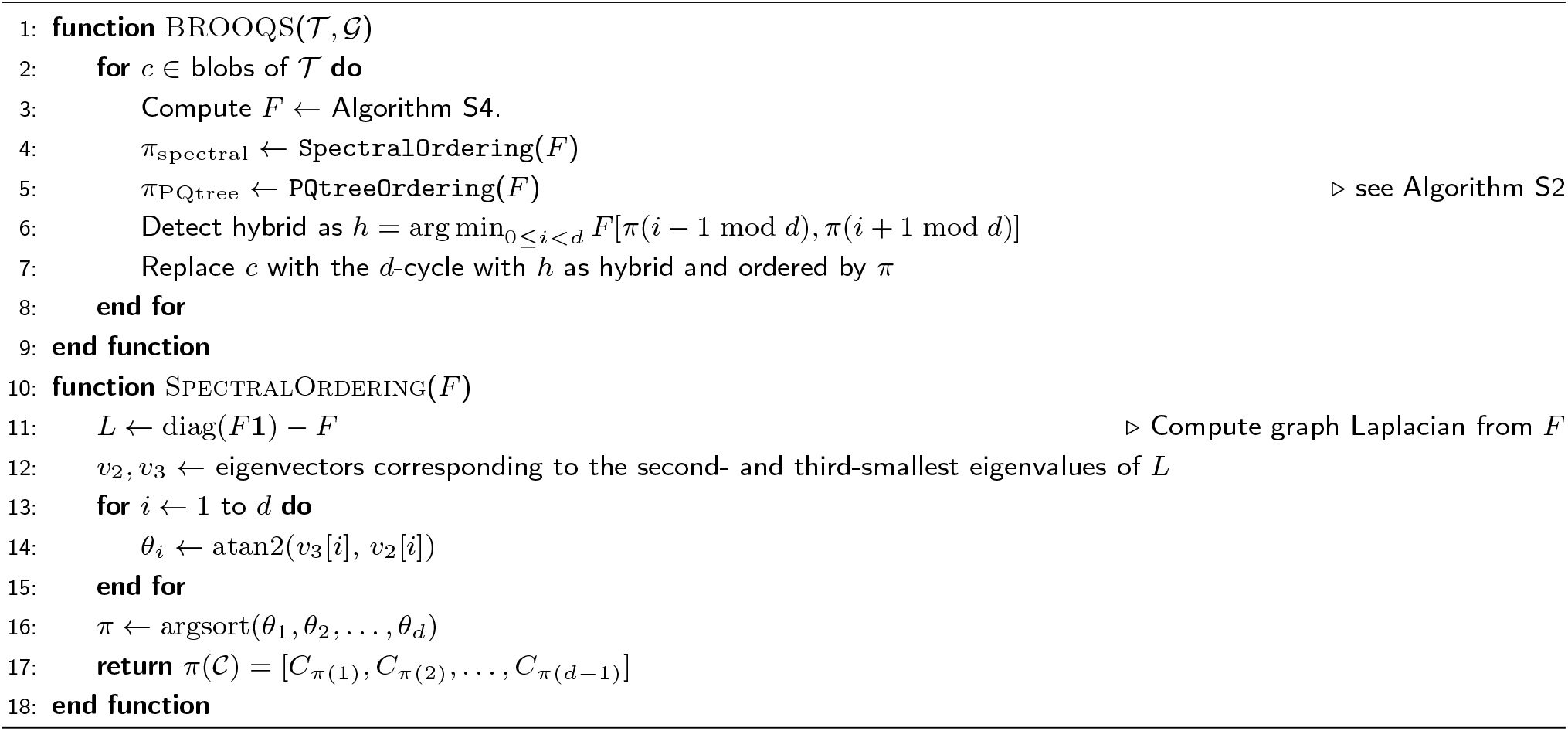

In BROOQS-Spectral, one additional step is taken to further improve robustness. Two consecutive components may have nearly identical embedded angles, and their relative order can be sensitive to the noise in *F* . Let *C*_*i*_ and *C*_*j*_ be two consecutive components in the ordering π(*C*) recovered by BROOQS-Spectral, with embedded angles *θ*_*i*_ and *θ*_*j*_ . If min 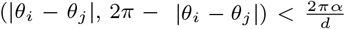 (we set *α* = 0.2), we resolve their relative order as follows. Let *C*_*l*_ and *C*_*r*_ be the components immediately before *C*_*i*_ and after *C*_*j*_ in π(*C*). If (*l, i, j, r*) is consistent with the true cyclic order, then by the strict circular Robinson property of *F* (Theorem 1), *F*[*l, i*] > *F*[*l, j*], *F*[*i, j*] > *F*[*l, j*], *F*[*i, j*] > *F*[*i, r*], and *F*[*j, r*] > *F*[*i, r*]. We count the violations of these inequalities in 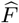 for the orders (*l, i, j, r*) and (*l, j, i, r*), and swap *C*_*i*_ and *C*_*j*_ in π(*C*) if the latter has fewer violations.

Finally, we ensure rootability. For a semi-directed network *N*, we say that rooting *N* on a node v ∈ V_*N*_ is valid if all hybrid edges point away from the root in the resulting rooted network. For a cycle, we define the hybrid score for the hybrid component *C*_*h*_ as Score_*h*_ = *F*[π(h − 1 mod *d*), π(h + 1 mod *d*)]. After resolving all cycles in *T*, if a valid rooting exists, we reroot *N* on a leaf that does not belong to any hybrid component. Otherwise, for each cycle we identify an alternative hybrid *C*_*h*_∈, the component with the lowest hybrid score among the remaining components. We then take the cycle with the smallest relative gap (Score_*h*_∈ − Score_*h*_)/Score_*h*_ between the scores of the alternative hybrid and the hybrid, and assign *C*_*h*_∈ as its hybrid without changing the cyclic order. We repeat this process, changing the hybrid of each cycle at most once, until a valid rooting of *N* is found. While this greedy process does not guarantee a valid rooted network in theory, in our analysis it never failed to return a valid rooted network.

##### Step 3: Identify the hybrid component

Assuming the recovered cyclic ordering π(*C*) is one of the equivalent true orderings, the hybrid component *C*_*π\**_ (0) is surrounded by its two true neighboring components *C*_*π\**_(1) and *C*_*π\**_ (*d*−1) in π(*C*). By Theorem 2, if *h* = arg min_0≤*i<d*_ *F*[π(*i* − 1 mod *d*), π(*i* + 1 mod *d*)], then *C*_*h*_ is the hybrid component of Ω. Thus, we can also identify the correct hybrid component from the F-matrix *F* by first finding the minimum value in *F* among components that are neighbors-but-one in π, and taking the component between them.

##### Optional heuristic resolution of 4-cycles

For 4-cycles, the hybrid component is not theoretically identifiable from quartet CFs (Baños, 2019; Allman et al., 2024a), i.e., the minimum F-score is not unique. Consequently, Theorem 2 does not extend to *d* = 4. Nevertheless, steps 1 and 2 of BROOQS are still correct, and thus the cyclic ordering of the four components is still identifiable. In our evaluations, we resolve 4-cycles by selecting one of the minimum F-score candidates at random as the hybrid component. While gene tree topology alone cannot recover the hybrid from CF values, other information could. In addition to the random selection, when gene trees have branch lengths and *T* can be rooted, we provide an optional heuristic to identify the hybrid. Let *d*^*G*^(*x, y*) be the total path length between *x* and *y* on a gene tree. We compute all pairwise distances between randomly selected representative taxa from each component other than root (say, *t*_1_, *t*_2_, *t*_3_) in each gene tree, find 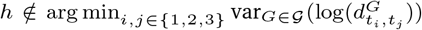, and designate the corresponding component as the hybrid. The intuition behind this heuristic is that, in a rooted 4-cycle, the pair of components with no hybrid descendant has only one minimum coalescence time, whereas the other two pairs, which contain the hybrid component, have two minimum coalescence times (Fig. S6a). Actual coalescent times are exponentially distributed above these minimums, but nonetheless, the non-hybrid node is expected to have less variation once we account for the mean. We use a log transformation of pairwise distances to focus on broad patterns and limit the impact of the mean and exponential coalescent times. Based on our experiments on a simulated dataset (Varying-ILS dataset in Section 2.3.1), using this heuristic instead of random hybrid selection could increase the accuracy of the resulting network, even in the presence of ILS and gene tree estimation error (Fig. S6b).

##### Complexity

To resolve *c* into a *d*-cycle, BROOQS first computes the empirical *d* × *d* matrix 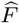 on *C* from the gene trees in *O*(*md*^2^*n*) time. To recover the cyclic ordering, BROOQS-PQtree recovers the cyclic ordering from 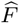 in *O*(*d*^2^) (Armstrong et al., 2021) using Algorithm S2, whereas BROOQS-Spectral computes the eigendecomposition of the graph Laplacian of 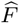 in *O*(*d*^3^) time and recovers the cyclic ordering of the components using Algorithm 1 in *O*(*d* log *d*) time. Finally, the hybrid component is identified in *O*(*d*) time. Since the computation of the empirical F-matrix dominates the remaining steps, resolving a single blob requires *O*(*md*^2^*n*) time. Therefore, resolving all blobs in *T* requires *O*(*mnD*), where *D* is the sum of squared blob degrees; i.e. 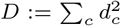. In the worst case, a blob can have degree *O*(*n*), resulting in a worst-case running time of *O*(*mn*^3^). However, our empirical analysis shows that *D* ≈ *O*(*n*^0.9^) for the typical datasets (Fig. S3a). Therefore, the observed running time of BROOQS is approximately *O*(*mn*^1.8^) (Fig. S3b).

### 2.3 Experiment Setup

We evaluated the performance of BROOQS on three simulated datasets and two biological datasets. On the simulated datasets, we asked whether BROOQS is more accurate and scalable than existing methods for tree of blobs resolution when the true tree of blobs is given (Experiment 1), and whether it can outperform end-to-end level-1 network inference methods given an inferred tree of blobs (Experiment 2). Finally, we ran BROOQS on two biological datasets (Experiment 3) and evaluated our findings based on the existing literature on these datasets.

#### 2.3.1 Simulated Datasets

##### Fixed-ILS

We used a dataset generated by Willson and Warnow (2026), with varying numbers of ingroup taxa (50 replicates each) *n* ∈ *{*15, 25, 50, 100, 150, 200*}* plus one outgroup. For each replicate, a true level-1 network was simulated using SiPhyNetwork (Justison et al., 2023), and 1000 true gene trees were simulated from each network using PhyloCoalSimulations (Fogg et al., 2023), resulting in 58% quartet similarity to the true tree of blobs. The average blob size in this dataset ranges between 7 and 14, depending on the model condition (Table S1a). Sequences were evolved using INDELible (Fletcher and Yang, 2009), and estimated gene trees were inferred using IQ-TREE 3 (Wong et al., 2026) for *n* ∈ *{*15, 25*}* and FastTree 2 (Price et al., 2010) for *n* ≥ 50, with an average gene tree estimation error of ≈ 20%.

##### Varying-ILS

To evaluate the effect of ILS on BROOQS, we also studied a dataset published by Dai et al. (2026). This dataset contains 50 level-1 network replicates for each *n* ∈ *{*50, 100, 200*}*, generated using a simulation process similar to that of the Fixed-ILS Dataset with similar statistics (Table S1b). However, to vary the effect of ILS, branch lengths of the true networks were multiplied by 2, 1, 0.5, and 0.25, resulting in 68%, 61%, 52%, and 43% quartet similarity to true tree of blobs, respectively. For each replicate, 1000 true gene trees were simulated using PhyloCoalSimulations, and estimated gene trees were inferred using IQ-TREE 3. Gene tree estimation error for this dataset varies from 40% to 70%.

##### Varying-ILS+GTEE

We also studied a dataset by Kolbow et al. (2026), where both ILS and gene tree estimation error (GTEE) are varied across model conditions. For each *n* ∈ {25, 50, 100, 200}, true level-1 networks were simulated using SiPhyNetwork, with two levels of ILS corresponding to average branch lengths of 1.5 (low ILS) and 0.15 (high ILS), resulting in 69% and 46% quartet similarity between true gene trees and tree of blobs, respectively and varying blob sizes (Table S1c). For each replicate, 1000 true gene trees were simulated using PhyloCoalSimulations, and estimated gene trees were inferred from sequences of length 100 bp and 1000 bp, resulting in average GTEE of ∼ 33% and ∼ 11%, respectively. On this dataset, we removed the analysis on estimated gene trees for replicates with fewer than 100 estimated gene trees (13 replicates out of 160 across all model conditions).

#### 2.3.2 Simulation Experiments

##### Experiment 1

We compared the performance of BROOQS to two state-of-the-art tree of blobs resolution methods, NANUQ+ (Allman et al., 2025) and NetCS (Dai and Molloy, 2026). Both methods use a two-step approach that first estimates a tree of blobs and then resolves it into a level-1 network. Thus, their blob resolution steps can be directly compared to BROOQS by providing all methods with the true tree of blobs. Since both methods use the same statistical tests of quartet topology, with parameters *α* and *β* for testing tree and star quartets, respectively, we set *α* = 1*e* − 7 and *β* = 0.95 for both methods in all simulation experiments, as this choice of parameters resulted in consistent positive results for both methods in previous studies (Dai et al., 2026). We ran BROOQS in both modes BROOQS-PQtree and BROOQS-Spectral, and used both true and estimated gene trees as input to all three methods. Each method was given 24 hours and 50GB of memory per replicate. With these constraints, both BROOQS and NetCS ran on all replicates, while NANUQ+ failed on all replicates where *n* ≥ 150.

We evaluated the methods based on blob resolution accuracy, running time, and memory usage. Since both BROOQS-PQtree and BROOQS-Spectral have the same time and memory complexity, we only report those metrics for BROOQS-Spectral. For accuracy, we report the hard-wired cluster distance (HWCD; Cardona et al. (2009)) between the true and estimated networks, which is an extension of Robinson-Foulds (RF) distance (Robinson and Foulds, 1981) to networks. We also report a new metric we define set to the minimum normalized RF distance (min nRF) between the trees displayed by the true and estimated networks, weighted by the inheritance probabilities of the former (see Section C.1 for a formal description). This metric measures how closely each evolutionary history in the true network can be explained by a displayed tree in the estimated network, while emphasizing evolutionary histories with higher total inheritance probability. Since resolving a blob requires 1) recovering the cyclic ordering and 2) identifying the hybrid component, we also report two metrics that separately evaluate these processes. For cyclic ordering accuracy, we use a modified Kendall-Tau error metric (Kendall, 1938) designed for cyclic orders, similar to the metric used by Dai and Molloy (2026) (see Section C.1 for a detailed description). For hybrid component identification, we report the proportion of hybrid components in the true network that are not correctly identified in the estimated network (FNR). Since hybrid components are not identifiable for 4-cycles, and consequently NetCS does not resolve 4-blobs, we report both metrics for cycles of size 5 or higher. In our method, to focus on the impact of our theoretically valid method for *d >* 4 rather than the *d* = 4 heuristic, we choose the hybrid component of 4-cycles uniformly at random in all simulation results.

##### Experiment 2

In the second experiment, we evaluated a full level-1 network inference pipeline (species tree→tree-of-blobs→network) with BROOQS used in the last step for resolving the tree of blobs. To infer the tree of blobs directly from a set of gene trees, we used the recent scalable method, TOB-QMC (Dai et al., 2026), combining the first two steps. We call this pipeline TOB-QMC+BROOQS. NetCS and NANUQ+ use the tree of blobs inferred by TOB-QMC and TINNIK (Allman et al., 2024b) by default in their full network inference pipeline; therefore, those trees of blobs were given as input to these methods. We also compared to two other standalone methods of network inference, CAMUS (Willson and Warnow, 2026) and SQUIRREL (Holtgrefe et al., 2025). CAMUS takes a displayed tree as input and adds reticulation edges to it to form cycles. We used the species tree inferred by ASTRAL-IV (Zhang et al., 2025) as input to CAMUS. Since CAMUS outputs a series of networks, with increasing numbers of reticulation events, we select the one that has the same number of reticulations as the tree of blobs inferred by TOB-QMC on the same set of gene trees. The input to SQUIRREL is a multi-sequence alignment (MSA) instead of a set of gene trees; we used the concatenation of the simulated sequences per gene as input to SQUIRREL, which is its intended way to run. All methods (except SQUIRREL) were run on both true and estimated gene trees and were given the 24-hour and 50GB memory constraints. We report HWCD and min nRF to evaluate the estimated networks for all methods. Since the tree of blobs can be wrong, we cannot report metrics relevant to blob resolution (Kendall-Tau and FNR).

#### 2.3.3 Real Data Analysis

To emphasize the scalability of the approach, we studied reticulation in two large phylogenetic datasets, one with many loci and another with a high number of species. For a high number of loci, we studied the avian dataset from Stiller et al. (2024) with 363 taxa and 63,430 gene trees. Previous studies suggest signs of hybridization at the base of Afroaves (Gatesy and Springer, 2022; Stiller et al., 2024), involving Accipitriformes (eagles, hawks, etc.), Strigiformes (owls), and Coraciimorphae (a diverse group of land birds including mousebirds, bee-eaters, woodpeckers, etc.). To examine large trees, we studied a plant dataset (Leebens-Mack et al., 2019) with 1,178 taxa and 410 gene trees. It has long been known that hybridization plays a major role in the evolution of plants (Anderson, 1948; Stull et al., 2023). The phylogenetic positions of several plant lineages (e.g., Gnetales) remain uncertain, with different analyses supporting conflicting placements (Wickett et al., 2014; Leebens-Mack et al., 2019). For each dataset, we ran TOB-QMC with the 3f1a setting (sampling *n* quartets around each branch) on the ASTRAL species tree and the estimated gene trees. We set the parameter for testing star quartets *β* = 0.95, and to correct for multiple tests inherent in testing many quartets around many branches, we use the Bonferroni correction and set the parameter for network quartet test *α* = *α*’/*n*^2^, where *n*^2^ is the total number of quartets sampled by TOB-QMC and *α* = 10^−2^ is the significance level used for both datasets (this leads to *α* = 7.6 × 10^−8^ for the avian and 7.2 × 10^−9^ for the plant dataset).

On the avian dataset, due to the large number of gene trees, statistical tests can become overly sensitive, picking up signals unrelated to hybridization due to slight deviations from symmetry. For example, running TOB-QMC on the full set of gene trees leads to 114 contracted branches and 31 blobs. To avoid such high levels of contractions (many of them likely false positives), we randomly split the gene trees into 11 sets, ran TOB-QMC on each set individually, and contracted a branch of the ASTRAL tree only if it was contracted by TOB-QMC in the majority of the 11 sets. This procedure resulted in 10 blobs (six 4-blobs and four with higher degrees). We ran BROOQS both on the full set of gene trees and on each of the 11 gene-tree partitions separately. For blobs of size 5 or more, we report the number of partitions that recover the same cyclic ordering and hybrid component as the analysis using the full set of gene trees. Due to its higher accuracy on simulated data and its nice visualizations, we only ran BROOQS in the spectral mode on both datasets.

We rooted the final tree of blobs output by TOB-QMC on the outgroup branch reported by the corresponding study. We then ran BROOQS on the estimated tree of blobs and the entire set of gene trees. For cycles with 5 or more components (*d >* 4), we report the *hybrid score* for each component *C*_*i*_ as 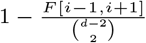. By Theorem 2, the component with the highest hybrid score is therefore picked as the hybrid component by BROOQS. For 4-blobs, we identified the hybrid component using the distance-based heuristic described earlier, applied as follows. We sampled 50 triplets of taxa randomly from the three non-root components and selected a component as the hybrid if the taxon sampled from that component was marked as a hybrid descendant in the majority of the triplets. We report the support for a hybrid in a 4-cycle as the proportion of the 50 triplets that agree with this choice.

## 3 Results

### 3.1 Experiment 1: True ToB

In Experiment 1, when the true tree of blobs was given to all methods, BROOQS-Spectral outperforms both NANUQ+ and NetCS across all simulated datasets and model conditions (Fig. 2). The magnitude of reduction in error depends on the dataset and model condition and can vary from 22% to 89% in HWCD, with higher gains observed in the more challenging cases. This advantage is achieved through both better hybrid component identification (evaluated by hybrid false negative rate) and more accurate cycle ordering (evaluated by Kendall-tau error). Comparing the two modes of BROOQS, BROOQS-PQtree is more accurate than the alternative methods in most conditions; however, it does not enjoy the accuracy and robustness of BROOQS-Spectral. Therefore, we focus on the spectral mode of BROOQS in subsequent sections.

**Figure 2.**
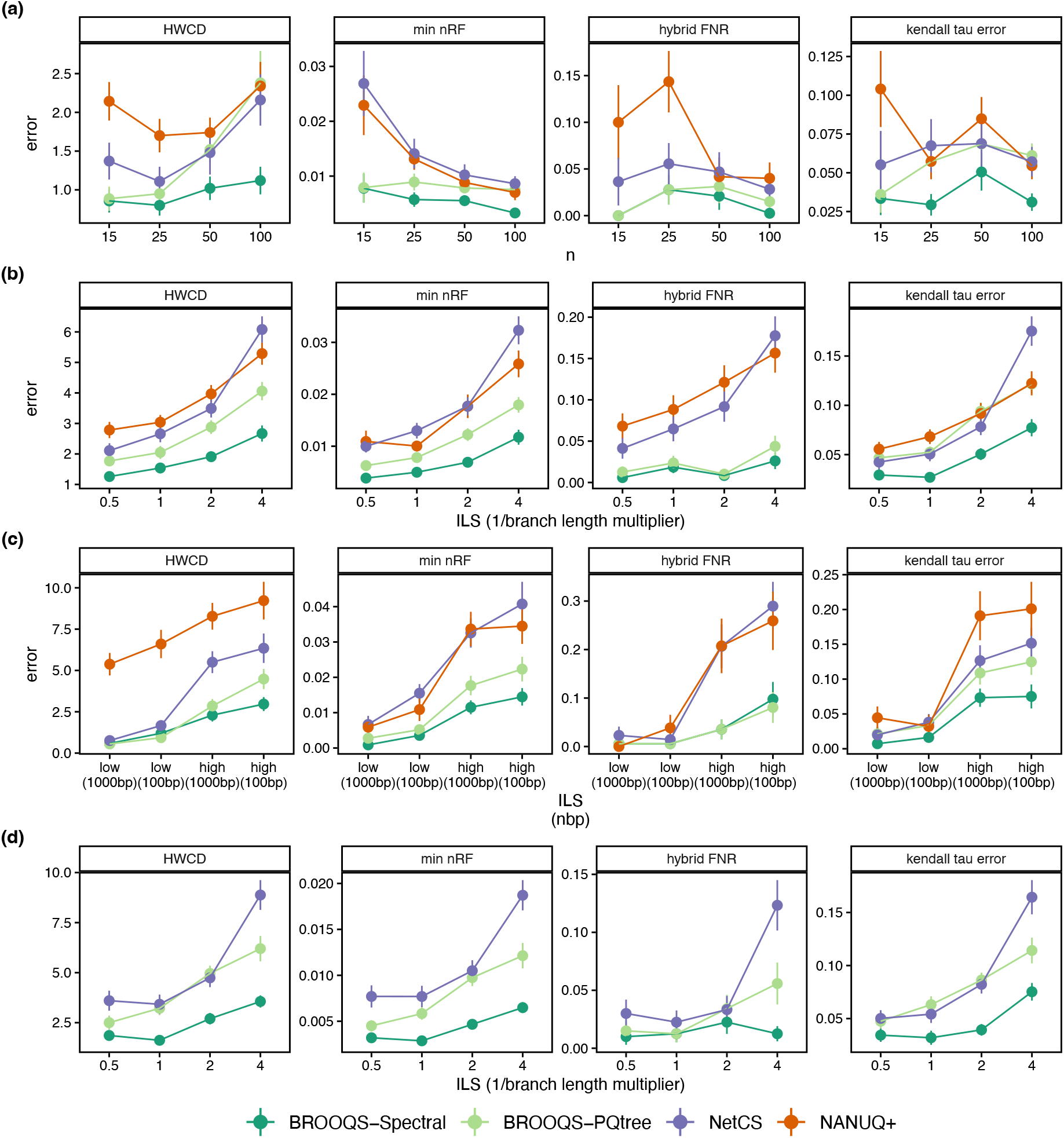
Experiment 1. Comparing BROOQS (in both Spectral and PQtree mode) to the alternative tree-of-blobs resolution methods, NANUQ+ and NetCS, on true tree of blobs. Panels compare error across three simulated datasets: Fixed-ILS **(a)**, Varying-ILS **(b,d)**, and Varying-ILS+GTEE **(c)**, and four error metrics: hard-wired cluster distance (HWCD), min nRF, False negative rate of hybrid component (FNR), and kendal-tau error. Panels **(a-c)** show *n* ≤ 100, where all three methods, BROOQS, NetCS, and NANUQ+ finished. Panel **(d)** shows *n* = 200, where only BROOQS and NetCS finished. Fig. S5 shows the same comparison on the other two simulated datasets. x-axis for the Varying-ILS **(b,d)** and Varying-ILS+GTEE **(c)** shows increased complexity. Results on true and estimated gene trees are merged for all datasets.

Focusing on the Fixed-ILS dataset with varying number of taxa, the HWCD remains stable for BROOQS-Spectral and increases only marginally for other methods as *n* increases (Fig. 2a). This is likely because the majority of the branches in the true tree of blobs are tree edges and do not contribute to HWCD. Similarly, min nRF decreases as the number of taxa increases, likely because the number of network edges does not increase linearly with the number of taxa. However, the identification of hybrid and the order also does not become worse with higher *n*, showing that BROOQS and the alternatives are robust to large tree sizes. Nevertheless, some of the biggest gains of BROOQS-Spectral are obtained for large *n*; HWCD error is reduced by 48% compared to NetCS, the second-best method, when *n* = 100.

Examining the Varying-ILS and Varying-ILS+GTEE datasets shows that BROOQS is not as sensitive to incomplete lineage sorting (ILS) as the other methods (Fig. 2b,c). While the performance of all methods suffers as levels of ILS increase, BROOQS, especially in spectral mode, is more robust to high ILS. NetCS, although slightly more accurate than NANUQ+, is the most sensitive method to ILS (Fig. 2b). Comparing the effects of ILS and gene tree estimation error (GTEE), controlled by sequence length, on the Varying-ILS+GTEE dataset shows that while both factors negatively affect the accuracy of all methods, the effect of ILS is much more pronounced (Fig. 2c). Like previous data, BROOQS is more robust to the effects of ILS than both NetCS and NANUQ+, and consistently outperforms both methods across all model conditions, with a greater advantage in the more challenging scenarios (high ILS and/or high GTEE).

On larger datasets (*n* > 100), we only compared BROOQS to NetCS. The advantage of BROOQS persists as the number of taxa increases, with BROOQS-Spectral substantially outperforming NetCS on the Varying-ILS dataset across all metrics and model conditions, particularly under the highest level of ILS, where BROOQS is 60% more accurate in terms of HWCD (Fig. 2d). This pattern is consistent across the other datasets (Fig. S5). While BROOQS-Spectral performs well in both hybrid component identification and cyclic ordering recovery, hybrid component identification is particularly robust to increasing ILS, with a hybrid FNR of 0.01 for BROOQS compared to 0.12 for NetCS at the highest level of ILS (Fig. 2d).

Not only is BROOQS significantly more accurate than the alternative methods, but it is also extremely efficient (Fig. 3). Empirically, our method has a subquadratic *O*(*n*^1.8^) running-time scaling with respect to *n* (Fig. S7a). Compared to the alternative methods, BROOQS is slightly slower than NetCS (eight minutes for BROOQS versus two for NetCS on *n* = 200), which has an empirical running-time scaling of *O*(*n*^1.6^) (Fig. S7a), but both methods easily extend to thousands of taxa. On the other hand, NANUQ+ did not extend to more than 100 taxa under our given constraints (24 hours and 50 GB memory), with an empirical running-time scaling of *O*(*n*^4.1^) (Fig. S7a). Similar to running time, BROOQS is also extremely efficient in memory usage. With linear memory usage with respect to n (Fig. S3c), it requires slightly less memory than NetCS, and both methods are drastically more memory-efficient than NANUQ+ (Fig. 3b).

**Figure 3.**
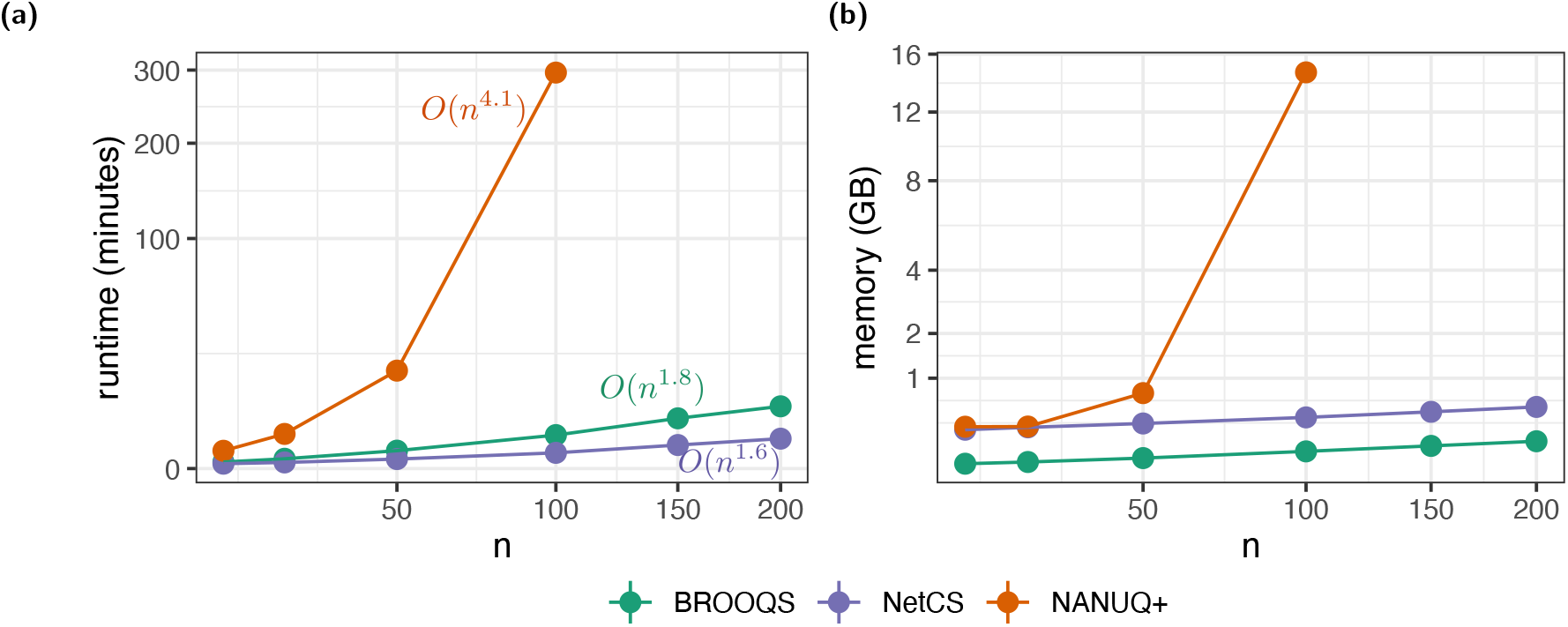
Experiment 1. Comparing the running time **(a)** and memory usage **(b)** of BROOQS to other blob resolution methods, NetCS and NANUQ+. Both axes are shown in square-root scale. Since both modes of BROOQS have the same runtime and memory complexity, we only show BROOQS-Spectral. Each method was given the true tree of blobs; therefore, the running time and memory usage of the ToB estimation step are not included for any of the methods. Note that NANUQ+ requires the output from TINNIK to resolve the tree of blobs, even when the true tree of blobs is provided as input. Thus, the running time of TINNIK is included for NANUQ+. Under the 24-hour and 50-GB memory constraints, NANUQ+ finished for *n* ≤ 100, while both BROOQS and NetCS ran for all numbers of taxa. For each *n*, true and estimated gene trees across all replicates from all three simulated datasets are merged. For the Varying ILS+GTEE dataset, only true gene trees are included (the number of estimated gene trees in this dataset varies drastically across replicates). The asymptotic running time was estimated using log-log slopes (see Figure S7a).

### 3.2 Experiment 2: Estimated ToB

When the estimated tree of blobs was given as input, both spectral and PQ-tree modes of BROOQS showed similar performances (Fig. S8). Therefore, we only show BROOQS-Spectral in comparison to other methods. On the Varying-ILS dataset, when given an estimated tree of blobs instead of the true tree of blobs, the accuracy of all three blob resolution methods drops drastically (Fig. S9), masking the gains from better tree-of-blobs resolution by BROOQS. Nevertheless, on the Varying-ILS dataset, BROOQS (run on TOB-QMC tree of blobs) is consistently among the best-performing methods across all metrics and model conditions (Fig. 4), a result further confirmed by the other simulated datasets (Figs. S11 and S12). BROOQS can be directly compared to NetCS since both methods use the same estimated tree of blobs by TOB-QMC. On the Varying-ILS dataset, BROOQS outperforms NetCS, although the improvements are small (Fig. 4). NANUQ+ tends to have higher error rates, especially at higher ILS levels.

**Figure 4.**
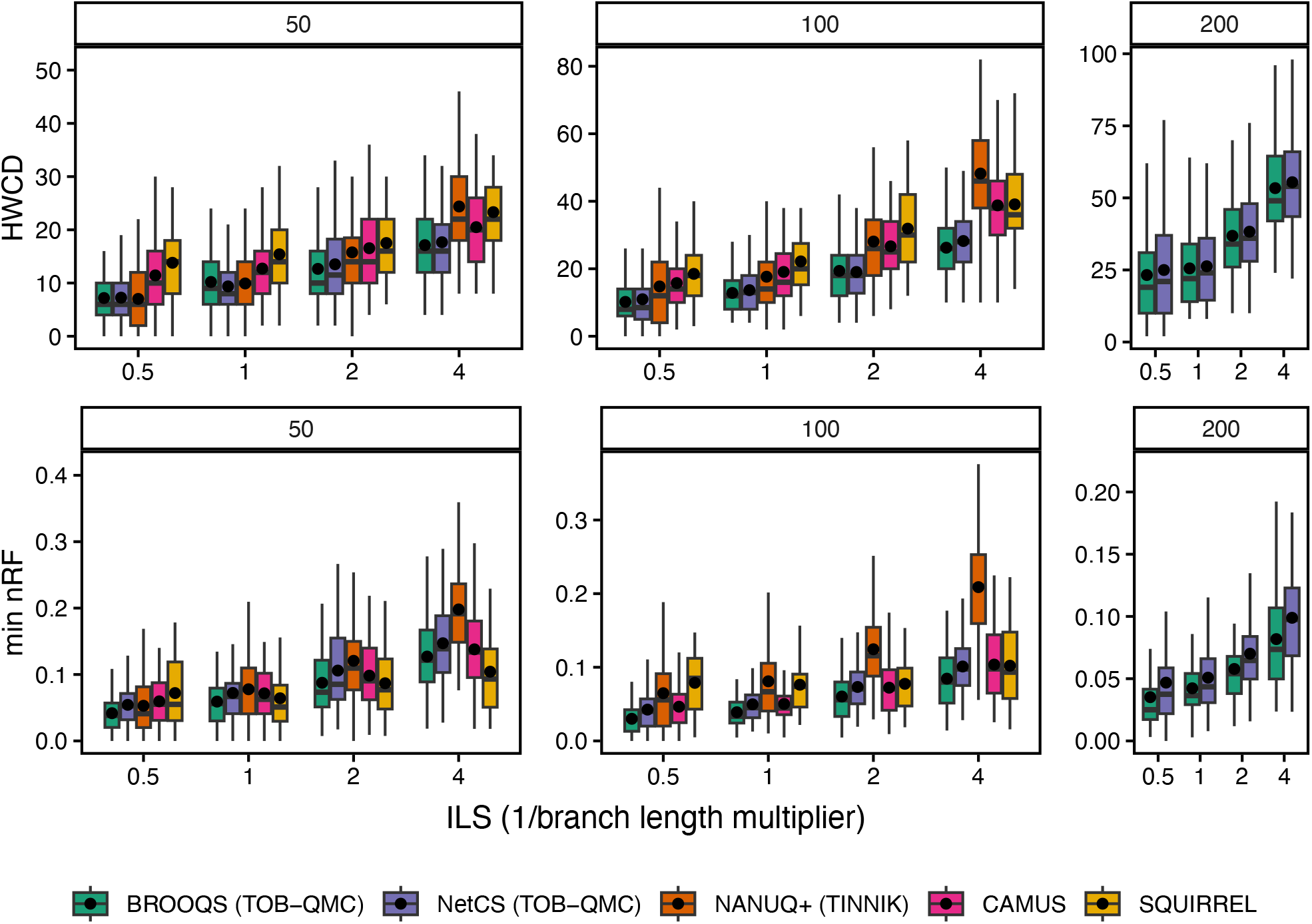
Experiment 2 (Varying-ILS dataset). Comparing BROOQS-Spectral on the estimated tree of blobs (TOB-QMC) with other network estimation methods. Methods are compared using two error metrics, HWCD and min nRF. Panels show the number of taxa (*n*). Under the 24-hour and 50-GB memory constraints, SQUIRREL, CAMUS and NANUQ+ finished for *n* ≤ 100. BROOQS and NetCS ran for all numbers of taxa. Also see Figs. S11 and S12.

Compared to the two standalone network inference methods, CAMUS and SQUIRREL, BROOQS consistently outperforms CAMUS. Note that we selected the CAMUS network with the same number of reticulations as the TOB-QMC tree of blobs, as CAMUS tends to substantially overestimate the number of reticulations with its default criterion (Fig. S10). SQUIRREL also overestimates the number of reticulations, although not as severely as CAMUS. This is reflected in high HWCD but low min nRF (Fig. 4); every extra reticulation is penalized by HWCD through additional false positive clusters, but as long as the true reticulations are captured closely, min nRF does not penalize extra reticulations (i.e., false positives). This shows that SQUIRREL has high recall but low precision.

In terms of scalability, BROOQS and NetCS remain the only methods that scale to 200 taxa under our computational constraints, even with the added running time of the tree-of-blobs estimation step by TOB-QMC. Both NetCS and BROOQS require less than one hour and 1GB of memory to finish on replicates with *n* = 100 (Fig. S7b). CAMUS scaled to replicates with *n* ≤ 150 under our constraints, while NANUQ+ and SQUIRREL only scaled up to *n* ≤ 100. SQUIRREL was the least scalable method in our analysis, requiring more than 12 hours and almost 25GB of memory for *n* = 100 (Fig. S7b).

### 3.3 Avian Dataset

In total, TOB-QMC identifies 10 blobs across the avian phylogeny (Fig. 5a-i). Six of these are 4-cycles, which we resolve using our proposed distance-based heuristic. Out of the 10 resolved cycles, six have an inferred hybrid source that is ancestral to the other source (compare H4 and H2 to H1 and H3 in Fig. 5 as examples). Such anachronistic hybridization events are often interpreted as at least one source of the reticulation being extinct or unsampled, a.k.a. a *ghost lineage* (Degnan, 2018; Ottenburghs, 2020). Among the 4-cycles, the support for the hybrid component (portion of triplets recovering it) varies from 0.4 to 0.8 across the reticulations (Fig. S13). All six 4-cycles are consistent with the ASTRAL tree from Stiller et al. (2024); that is, the ASTRAL tree is displayed within the inferred cycles. Because the strength of this heuristic is not our focus, we do not further investigate these reticulation events. We focus on the four larger cycles, marked as H1 to H4 in Fig. 5a-i. Of these four cycles, H1 and H4 are consistent with the ASTRAL tree, while H2 and H3 are not. Next, we examine each of these reticulation events.

**Figure 5.**
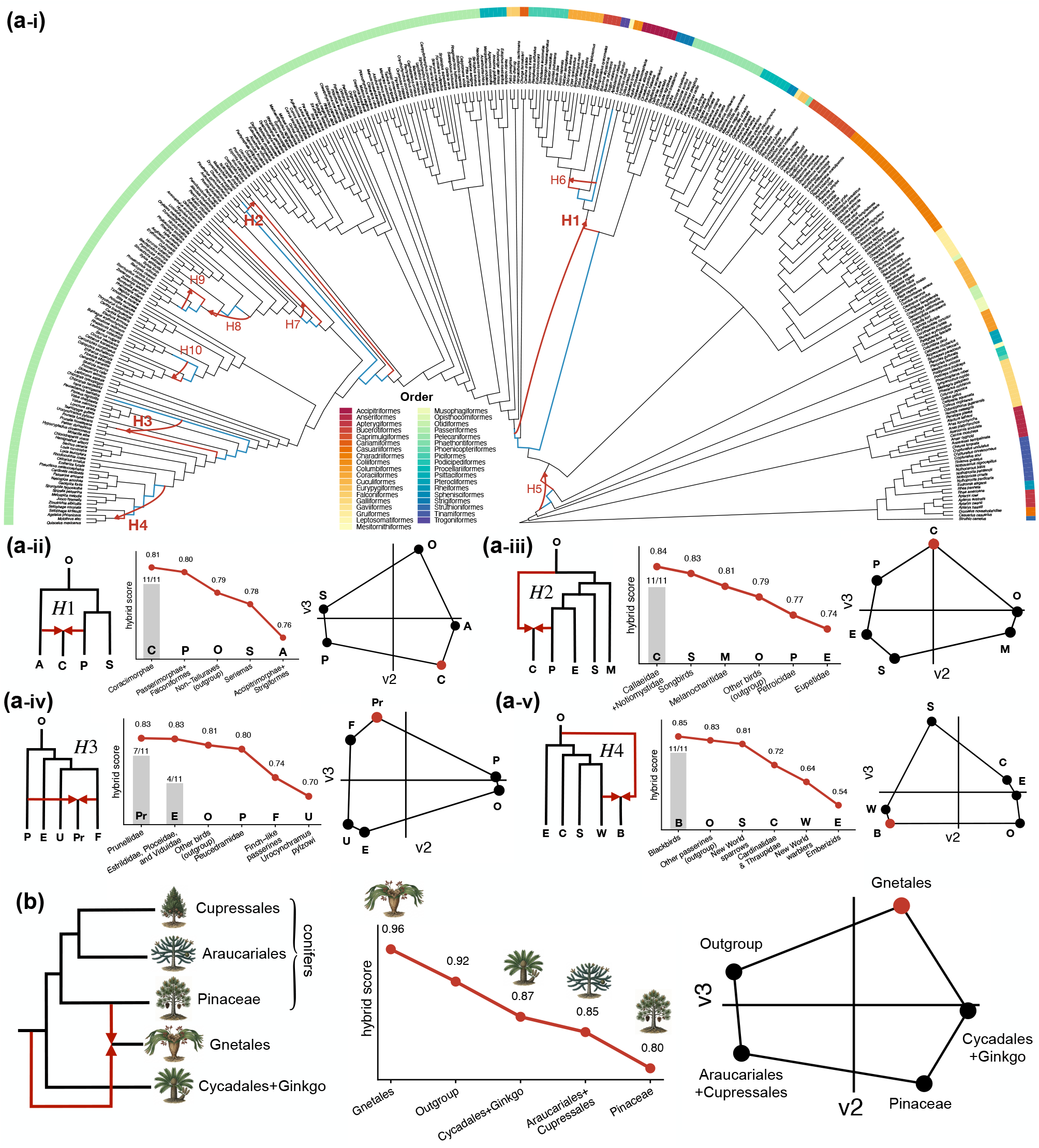
**(a)** Avian dataset. **(a-i)** Inferred phylogenetic network by BROOQS. Cycles are labeled from H1 to H10. For each cycle, the two hybrid edges are marked with red, and the rest of the edges in the cycle are marked with blue. See Fig. S13 for the support of the hybrid component of the 4-cycles H5-H10. The network is rooted at the common ancestor of Tinamus and Ostrich. Leaves are colored by taxonomic order. **(a-ii to a-v)** For each inferred cycle of size 5 or larger (H1-H4), we show the rooted topology of the inferred cycle by BROOQS, consistent with the network in **(a-i)**. We also show the hybrid score of each component in decreasing order. Highest score (i.e. arg min*h*(*F* [*h* −1,*h* + 1]) in *Methods*) is selected as hybrid by BROOQS. Bars show the number of 11 partitions that select each component as the hybrid. Note that all 11 partitions recover the same cyclic ordering for H1-H4. Finally, for each cycle, we show the embedding of each component on the eigenvector space induced by the corresponding F-matrix as well as the closed path that recovers the cyclic ordering. **(b)** Plants dataset. Rooted topology of the 5-cycle involving Gnetales, Pinaceae, the clade of Araucariales+Cupressales, and the clade of Cycadales+Ginkgo. We show the embedding of each component on the eigenvector space induced by the corresponding F-matrix as well as the closed path that recovers the cyclic ordering. We also show the hybrid score of each component in decreasing order. See Fig. S15 for the full tree with 1,178 taxa.

#### H1

Whether Strigiformes (owls) should be sister to Accipitriformes (eagles, hawks, etc.) or Coraciimorphae (a diverse set of birds, including mousebirds, woodpeckers, kingfishers, and trogons) has long been debated (Jarvis et al., 2014; Suh, 2016; Gatesy and Springer, 2022). Many studies have also reported signs of introgression within Strigiformes (Hanna et al., 2018). As a result, it has been hypothesized that Strigiformes may be a hybrid descendant of Accipitriformes and Coraciimorphae (Gatesy and Springer, 2022). In our analysis, however, TOB-QMC contracts the branch above Strigiformes+Accipitriformes in only 1 out of 11 gene tree partitions (average tree test p-value of 1.4*e*−6> α = 7.6*e* − 8; Fig. S14), which leads to Strigiformes and Accipitriformes remaining together as one component in the blob. Instead, Strigiformes+Accipitriformes appear in a 5-cycle along with Coraciimorphae, Passerimorphae+Falconiformes, and Seriemas (Fig. 5a-ii ). On this cycle, BROOQS identifies the Coraciimorphae group as the hybrid component, descending from Accipitriformes+Strigiformes and Passerimorphae+Falconiformes. Interestingly, the Accipitriformes+Strigiformes group has the lowest hybrid score according to BROOQS, making it the least likely candidate for the hybrid component. Moreover, all 11 gene tree partitions recover the same cyclic ordering and hybrid component (Fig. 5a-ii), providing strong support for this reticulation.

#### H2

The relationships among Petroicidae, Callaeidae, and Notiomystidae have been difficult to resolve, as these three families represent early-diverging lineages of Passerimorphae and previous studies have recovered conflicting relationships among them (Mackiewicz et al., 2019; Lubbe et al., 2022). In particular, previous analyses have provided evidence supporting both the Petroicidae+Notiomystidae and Callaeidae+Notiomystidae topologies (Mackiewicz et al., 2019; Lubbe et al., 2022). Despite this conflicting signal, the branch separating Callaeidae and Notiomystidae was not contracted by TOB-QMC (average tree test *p*-value of 0.004; Fig. S14). Nevertheless, BROOQS identifies a genetic contribution from Petroicidae to the branch resulting in Callaeidae+Notiomystidae (Fig. 5a-iii). This reticulation, which was supported by all 11 gene tree partitions, may explain part of the conflicting phylogenetic signal among these three lineages, including the independent gain of mitogenomic duplications in Petroicidae and Notiomystidae (Mackiewicz et al., 2019).

#### H3

We also resolve a blob among early branches of the Passerida radiation, involving Peucedramidae (olive warbler) and a diverse clade including the new World nine-primaried oscines and other families (which we call finch-like passerines, Fig. 5a-iv) to produce Prunellidae. Although early studies suggested Prunellidae and Peucedramidae as sisters (Ericson and Johansson, 2003), this relationship has not been consistently recovered by subsequent studies (Moyle et al., 2016; Stiller et al., 2024). The evidence supporting this reticulation is more mixed. Although all 11 gene tree partitions recover the same cyclic ordering, only 7 partitions identify Prunellidae as the hybrid descendant, while the remaining 4 partitions identify the common ancestor of Estrildidae, Ploceidae, and Viduidae as the hybrid (Fig. 5a-iv). This is also reflected in the hybrid scores, with the score for Prunellidae only 0.001 higher than that of the common ancestor of Estrildidae, Ploceidae, and Viduidae. We also observe that the embedding of the components of this cycle in the eigenvector space groups the components into three pairs that are close to each other within each pair but more distant across pairs (Fig. 5a-iv), a pattern that calls the veracity of the identified blob into question (more on this in the *Discussion*).

#### H4

Finally, we find the common ancestor of the blackbirds (Agelaius phoeniceus, Molothrus ater, and Quiscalus mexicanus) from the Icteridae family to be a descendant of a hybridization involving a ghost lineage (Fig. 5a-v). Both the cyclic ordering and hybrid component are supported by all 11 gene tree partitions. Although there have been previous reports of hybridization among several members of this group, including between Quiscalus mexicanus and Agelaius phoeniceus (Powell and Kirkley, 2021), we could not find a study reporting a reticulation event involving the common ancestor of these three species.

### 3.4 Plants Dataset

TOB-QMC finds 9 blobs on the plant dataset, of which only one is a 5-blob and the rest are 4-blobs (Fig. S15). The 5-blob involves the resolution of Gymnosperms, including Gnetales, Pinaceae, the ancestor of Cycadales and Ginkgo, and the rest of the conifers; i.e. Araucariales and Cupressales (Fig. 5b and *H4* in Fig. S15). The resolution of Gymnosperms broadly, and the position of Gnetales specifically, has been highly controversial. Various topologies (Gnetales+Pinaceae, Gnetales as a sister to all conifers, Gnetales as a sister to Araucariales and Cupressales, and Gnetales as a sister to conifers and Cycadales+Ginkgo) have been supported by previous analyses (Nickrent et al., 2000; Chaw et al., 2000; Leebens-Mack et al., 2019; Wickett et al., 2014). While ASTRAL recovers the ‘Gnetifer’ (Gnetales as a sister to all conifers) topology on this dataset, the short internal branches and asymmetry of minor CFs have been attributed to rapid diversification, gene-tree estimation biases, and potential gene flow (Leebens-Mack et al., 2019).

BROOQS infers Gnetales confidently as the hybrid component (0.04 higher hybrid score than the second candidate; Fig. 5b). This identification is consistent with the difficulty of placing Gnetales in the previous literature seeking to resolve Gymnosperms. The resulting network displays two of the debated topologies: Gnetales+Pinaceae, and Gnetales as sister to the rest of Gymnosperms. One valid interpretation of this reticulation event is that Gnetales are sister to pines, with some gene flow from an unknown ghost lineage that was sister to all present-day Gymnosperms. The fact that one of the contributors is ghost may explain why Gnetales as sister to all other Gymnosperms has been less commonly recovered by previous tree-based analyses (Wickett et al., 2014; Leebens-Mack et al., 2019). Consistent with our results, a similar displayed tree on the same dataset is recovered by InPhyNet, a divide-and-conquer approach for scaling likelihood-based network inference methods to larger datasets (Kolbow et al., 2026). However, InPhyNet supports the ‘Gnetifer’ hypothesis as the second alternative displayed tree.

TOB-QMC infers eight 4-blobs spanning several groups within the plant phylogeny, including asterids, lycophytes, and ferns (Fig. S15). We resolve these blobs using our distance-based heuristics. However, we do not report support for the inferred hybrid components, as many of these events involve only a few taxa, for which the heuristic used to identify the hybrid component and estimate its support is less reliable. Nonetheless, several of the hybrid descendants identified by BROOQS are consistent with previous findings. For example, for the reticulation within Lycopodiales, BROOQS identifies the genus *Diphasiastrum* as the hybrid descendant (*H1* in Fig. S15), which has previously been reported to be of hybrid origin (Bennert et al., 2011). In *Polypodium*, a genus with extensive reticulate evolution and numerous reported interspecific hybrids and allopolyploid species (Sigel et al., 2014a,b), BROOQS identifies *Polypodium hesperium* as the hybrid descendant (*H2*); this species was also identified as a hybrid descendant by InPhyNet (Kolbow et al., 2026). Finally, BROOQS identifies *Encephalartos barteri* as a hybrid descendant (*H3*), consistent with the complex relationships and numerous interspecific hybrids previously reported in *Encephalartos* (Habib et al., 2026).

## 4 Discussion

We introduced BROOQS, a method that resolves a tree of blobs into a level-1 semi-directed network using a quartet-based similarity matrix, *F*, which we show encodes the circular order of components around the blob by guaranteeing a set of inequalities. Unlike alternative methods such as NANUQ+ and NetCS, BROOQS does not rely on statistical tests on individual quartets. Nor does it subsample or list all quartets. Instead, BROOQS aggregates information from all quartets around a blob without sacrificing scalability or relying on quartet subsampling. It combines results into the similarity F-matrix, and uses the ordinal properties of this matrix to resolve each blob. As a result of its efficient methods to compute the F-matrix, BROOQS is extremely scalable. To recover the order from F-matrix, we use either a combinatorial or spectral method. The combinatorial version, which uses the PQ-tree algorithm of Armstrong et al. (2021), allows us to prove statistical consistency for blobs of size 5 to 15, while the spectral version is only conjectured to be consistent. Despite its stronger theoretical proofs, BROOQS-PQtree appeared less robust to noise in *F*, prompting us to use BROOQS-Spectral by default. The observation that a spectral method is more robust to noise than an iterative approach that merges the nearest neighboring components in each step is not surprising to us. Generally, spectral methods tend to be robust to noise, and this has been among their main appeals across several domains (Karoui and Wu, 2016; Fogel et al., 2016; Stephan and Massoulié, 2019). We hope that future work will be able to prove consistency of BROOQS-Spectral by proving Conjecture 1.

Our simulation results indicate that higher levels of ILS degrade the accuracy of all blob resolution methods, but BROOQS-Spectral is more robust than alternatives. High ILS dilutes the signal of quartet concordance factors (CFs), on which NANUQ+, NetCS, and BROOQS rely. However, algorithmic details matter, and BROOQS uses the diluted CF signal more efficiently. Beyond general robustness of spectral methods, two more specific reasons can be suggested here. First, when ILS is high, the CFs of individual quartets become closer to 1*/*3, reducing the power of the statistical tests used by alternative methods. BROOQS uses no such test. Second, higher ILS, paired with gene tree estimation error, can lead to noisy CFs that change the ranking of topologies. Quartets that include distant outgroups can further suffer from high levels of missing data, impacting CF rankings. Both NANUQ+ and NetCS use ranking of CFs of individual quartets to make decisions when resolving blobs. NANUQ+ scores a particular ordering based on all CF rankings of quartets, but each ranking can be noisy (note that ranking alone masks the magnitude of differences between CFs). NetCS constructs the cyclic ordering through a series of choices based on majority voting (hybrid component, neighboring components), where an incorrect decision at one step can affect subsequent decisions. BROOQS reduces these issues in two ways. It strengthens the signal by averaging the quartet frequencies across all quartets around a blob, avoiding reliance on noisy rankings for individual quartets. This averaging effect across quartets may also make it more robust to more subtle issues such as rogue taxa and missing data from distant outgroups. Furthermore, rather than inferring the cyclic ordering in a series of steps, the spectral version of BROOQS infers the ordering of all components simultaneously, without conditioning on a particular hybrid component. Comparing the two modes of our method, both of which use the same F-matrix, further highlights the benefit of inferring the orders jointly (BROOQS-Spectral) rather than iteratively (BROOQS-PQtree).

Across all simulations, given the true tree of blobs, BROOQS-Spectral outperforms alternative methods by a wide margin. When the tree of blobs is estimated, BROOQS also remains consistently among the best-performing methods, albeit with a lower gap. While we leave comparisons to the less scalable likelihood-based network inference methods to future work, our comparison to other quartet-based methods provides support for the general strategy of first estimating a tree of blobs and then resolving blobs into cycles. Although our results indicate that errors in the first step propagate to the second step, this two-step approach achieves comparable or better performance than alternative approaches on our data, particularly in estimating the number of reticulations (Fig. S10). Among the two alternative standalone network inference methods, SQUIRREL follows a similar strategy by first inferring a set of multifurcating trees (a.k.a. trees of blobs) and then resolving each one using quarnets inferred from the multiple sequence alignment. However, it does not model ILS, which may contribute to its overestimation of reticulation events (Fig. S10). Thus, while there is room for improvement in detecting blobs, the general strategy of the two-step approach appears promising.

Estimating an accurate tree of blobs, however, remains key. Our results suggest that, in the two-step approach, the error introduced when inferring the tree-of-blobs dominates the error introduced when resolving polytomies (Fig. S9). While this paper only focused on blob resolution, we believe that the signal in the F-matrix *F* and its embedding in the eigenspace are also informative of whether a set of components forms a blob in the underlying level-1 network. We illustrate this observation with an example (Fig. S16a,b). Given the true tree of blobs, the components in a *d*-cycle form a closed path with *d* distinct embeddings in the eigenspace (Fig. S16c). Contracting an additional tree edge adjacent to the blob (i.e., partitioning one component into two while keeping the remaining components intact) results in the two partitions to have near-identical embeddings (Fig. S16d). Thus, this type of signal may be able to identify when a blob is over-contracted. In the other extreme, contracting a set of adjacent *tree edges* into a completely incorrect blob results in embeddings that resemble a triangle with many overlapping components rather than a circle (Fig. S16e). These observations suggest that the geometry of the spectral embedding may contain information about whether the inferred components form a true blob. We leave a more extensive analysis of this observation and its potential for obtaining more accurate trees of blobs to future work.

The F-matrix used in this paper also has a direct connection to pairwise similarity matrices long used in distance-based tree inference. Although we compute the F-matrix over a set of components, each containing a partition of *n* taxa, the same matrix can be computed as an *n* × *n* pairwise similarity matrix between individual taxa. Previous work has shown that, although this quartet-based similarity matrix is not itself an additive distance matrix, a similar one defined based on the negative log transformation of CFs for each quartet is additive (Sayyari and Mirarab, 2016). This result is particularly interesting in light of a main finding by Jaffe et al. (2021): for a pairwise similarity matrix *R* = *e*^*−D*^, where *D* is an additive distance matrix on a tree, spectral algorithms combined with neighbor joining can recover the true tree. A recent study also shows that a divide-and-conquer approach using spectral algorithms on such matrices is statistically consistent under MSC (Reshef et al., 2026). Together with our observation that the F-matrix computed over a set of components with a tree-like structure separates the components into disjoint groups corresponding to the tripartitions around nodes in the true tree (Fig. S16e), these connections encourage further exploration of the F-matrix and spectral algorithms for phylogenetic network inference directly from F.

Several aspects of the blob resolution can also be improved. A main limitation of any method that relies on quartet CFs is that it cannot identify the hybrid component in resolving 4-blobs (Baños, 2019; Allman et al., 2024a). We suggested a distance-based heuristic to identify the hybrid in 4-blobs when the rooting is known and branch lengths are given. Our simulations show that this heuristic is promising, even when applied to estimated gene trees (Fig. S6b). However, the method remains a crude heuristic, which may not work well in the presence of substantial rate variation across loci. Future work should develop more robust methods for identifying the hybrid component in 4-cycles using branch lengths and, more broadly, incorporating gene tree branch lengths in resolving blobs. Moreover, we use a heuristic to correct for missing components in gene trees. Our simulation analysis shows the effectiveness of this heuristic (Fig. S4). On the avian dataset, on average less than 10% of gene trees have entire components missing per blob. Filtering out these gene trees when running BROOQS inferred the same network, suggesting the robustness of the heuristic on real datasets with a small amount of missing components. However, this heuristic assumes that missing components are independent across the blob, an assumption that may not always hold. Future work should explore more complex patterns of missing data. Furthermore, future work should develop statistical approaches to assess the confidence in inferred cycles. On the avian dataset, we report a measured support as the proportion of gene tree splits that agree with its cyclic ordering and hybrid identification (three of the four major cycles had 100% support). But more direct bootstrapping methods may provide more reliable support, particularly for datasets with fewer gene trees. Finally, future work should also explore inferring cycle parameters, such as the inheritance probability *γ* and branch lengths from the F-matrix. Such information may also be relevant to confidence of the inferred cycle. For example, an estimated *γ* very close to zero or near-zero branch lengths may indicate that the inferred cycle is weakly supported or may not represent a true reticulation.

Our analyses of real datasets demonstrated scalability of the TOB-QMC+BROOQS to modern phylogenomic datasets with large numbers of taxa and genes. BROOQS took only 2 minutes to run on the plant dataset with 1,178 taxa and 410 gene trees, and around 5 hours on the avian dataset with 363 taxa and 63,430 gene trees. And its results were in line with prior knowledge. It recovers reticulation patterns for several highly debated avian and plant groups that are consistent with previous studies, while also identifying some new potential hybrids. For example, the inferred placement of Gnetales with respect to conifers is consistent with previously reported relationships, whereas the inferred hybrid origin of Coraciimorphae, rather than Strigiformes, differs from some previous hypotheses and warrants further investigation. The results, therefore, indicate that the methodology laid out in BROOQS can be applied to real data as is, and that the approach can be extended in the future to infer the tree of blobs and network parameters.

## 6 Acknowledgments

We thank Prof. Tandy Warnow and The-Anh Vu-Le for carefully reading our paper and graciously pointing out a mistake in an earlier version of the paper, as well as a counterexample in Table S3. We also thank Dr. Erin K. Molloy and Junyan Dai for generously sharing their data and for providing assistance with the use of their tool, NetCS.

## 7 AI Use Statement

We used generative AI tools (specifically Claude Opus) for the numerical proof of Proposition 2, the extensive verification of our theory described in Section C.2, and the implementation of the PQ-tree algorithm by Armstrong et al. (2021). All other proofs are free of AI use. The AI work is checked by the authors.

## 8 Data and Code Availability

The software is available at github.com/shayesteh99/BROOQS. All data analyzed in this article are available on Dryad (https://doi.org/10.5061/dryad.6hdr7srgv) and on GitHub at github.com/shayesteh99/BROOQS-Data.

## References

Adams, R. H., D. R. Schield, D. C. Card, and T. A. Castoe. 2018. Assessing the impacts of positive selection on coalescent-based species tree estimation and species delimitation. Systematic Biology 67:1076–1090.

Allman, E. S., H. Baños, M. Garrote-Lopez, and J. A. Rhodes. 2024a. Identifiability of level-1 species networks from gene tree quartets.Bull Math Biol 86:110.

Allman, E. S., H. Baños, J. D. Mitchell, and J. A. Rhodes. 2022. The tree of blobs of a species network: identifiability under the coalescent. J Math Biol 86:10.

Allman, E. S., H. Baños, J. D. Mitchell, and J. A. Rhodes. 2024b. Tinnik: inference of the tree of blobs of a species network under the coalescent model. Algorithms for Molecular Biology 19:23.

Allman, E. S., H. Baños, and J. A. Rhodes. 2019. Nanuq: a method for inferring species networks from gene trees under the coalescent model. Algorithms Mol Biol 14:24.

Allman, E. S., H. Baños, J. A. Rhodes, and K. Wicke. 2025. Nanuq+: A divide-and-conquer approach to network estimation. Algorithms for Molecular Biology 20:14.

Allman, E. S., J. H. Degnan, and J. A. Rhodes. 2011. Identifying the rooted species tree from the distribution of unrooted gene trees under the coalescent. J Math Biol 62:833–862.

Anderson, E. 1948. Hybridization of the habitat. Evolution 2:1–9.

Armstrong, S., C. Guzmán, and C. A. Sing Long. 2021. An optimal algorithm for strict circular seriation. SIAM Journal on Mathematics of Data Science 3:1223–1250.

Baños, H. 2019. Identifying species network features from gene tree quartets under the coalescent model. Bulletin of Mathematical Biology 81:494–534.

Bennert, H. W., K. Horn, M. Kauth, J. Fuchs, I. S. B. Jakobsen, B. Ollgaard, M. Schnittler, M. Steinberg, and R. Viane. 2011. Flow cytometry confirms reticulate evolution and reveals triploidy in central european diphasiastrum taxa (lycopodiaceae, lycophyta). Ann Bot 108:867–876.

Bjornson, S., H. Verbruggen, N. S. Upham, and J. L. Steenwyk. 2024. Reticulate evolution: Detection and utility in the phylogenomics era. Molecular Phylogenetics and Evolution 201:108197.

Booth, K. S. and G. S. Lueker. 1976. Testing for the consecutive ones property, interval graphs, and graph planarity using pq-tree algorithms. Journal of Computer and System Sciences 13:335–379.

Bryant, D., J. Tsang, P. Kearney, and M. Li. 2000. Computing the quartet distance between evolutionary trees. Pages 285–286 in Proceedings of the Eleventh Annual ACM-SIAM Symposium on Discrete Algorithms SODA ‘00 Society for Industrial and Applied Mathematics, USA.

Cardona, G., M. Llabrés, F. Rosselló, and G. Valiente. 2009. Metrics for phylogenetic networks i: generalizations of the robinson-foulds metric. IEEE/ACM Trans Comput Biol Bioinform 6:46–61.

Chaw, S. M., C. L. Parkinson, Y. Cheng, T. M. Vincent, and J. D. Palmer. 2000. Seed plant phylogeny inferred from all three plant genomes: monophyly of extant gymnosperms and origin of gnetales from conifers. Proc Natl Acad Sci U S A 97:4086–4091.

Coifman, R. R., Y. Shkolnisky, F. J. Sigworth, and A. Singer. 2008. Graph laplacian tomography from unknown random projections.IEEE Trans Image Process 17:1891–1899.

Dai, J., Y. Han, and E. K. Molloy. 2026. Quartet-based species tree methods enable fast and consistent tree of blobs reconstruction under the network multispecies coalescent. Genome Research Page gr.282203.126.

Dai, J. and E. K. Molloy. 2026. Is Level-1 Blob Reconstruction Under the Network Multispecies Coalescent Easy? Pages 1:1–1:24 in 26th International Conference on Algorithms for Bioinformatics (WABI 2026) (N. El-Mabrouk and F. Vandin, eds.) vol. 390 of Leibniz International Proceedings in Informatics (LIPIcs) Schloss Dagstuhl – Leibniz-Zentrum für Informatik, Dagstuhl, Germany.

Degnan, J. H. 2018. Modeling hybridization under the network multispecies coalescent. Systematic Biology 67:786–799.

Ericson, P. G. and U. S. Johansson. 2003. Phylogeny of passerida (aves: Passeriformes) based on nuclear and mitochondrial sequence data. Molecular Phylogenetics and Evolution 29:126–138.

Felsenstein, J. 1978. Cases in which parsimony or compatibility methods will be positively misleading. Systematic Biology 27:401–410.

Fletcher, W. and Z. Yang. 2009. Indelible: A flexible simulator of biological sequence evolution. Molecular Biology and Evolution 26:1879–1888.

Fogel, F., A. d’Aspremont, and M. Vojnovic. 2016. Spectral ranking using seriation. Journal of Machine Learning Research 17:1–45.

Fogg, J., E. S. Allman, and C. Ané. 2023. Phylocoalsimulations: A simulator for network multispecies coalescent models, including a new extension for the inheritance of gene flow. Systematic Biology 72:1171–1179.

Frankel, L. E. and C. Ané. 2023. Summary tests of introgression are highly sensitive to rate variation across lineages. Systematic Biology 72:1357–1369.

Gatesy, J. and M. S. Springer. 2022. Phylogenomic coalescent analyses of avian retroelements infer zero-length branches at the base of neoaves, emergent support for controversial clades, and ancient introgressive hybridization in afroaves. Genes 13.

Gusfield, D., V. Bansal, V. Bafna, and Y. S. Song. 2007. A decomposition theory for phylogenetic networks and incompatible characters.Journal of Computational Biology 14:1247–1272.

Habib, S., A. Lindstrom, J. A. Clugston, Y. Gong, S. Dong, Y. Wang, D. Stevenson, C. Feng, and S. Zhang. 2026. Integrative phylogenomics and morphology reveal the evolution and biogeography of encephalartos (zamiaceae). Journal of Systematics and Evolution 64:295–312.

Hanna, Z. R., J. P. Dumbacher, R. C. K. Bowie, J. B. Henderson, and J. D. Wall. 2018. Whole-genome analysis of introgression between the spotted owl and barred owl (strix occidentalis and strix varia, respectively; aves: Strigidae) in western north america. G3 Genes—Genomes—Genetics 8:3945–3952.

He, C., M.-Y. Chen, and H. Zhu. 2026. Population size differences can lead to biases in phylogenetic inference and introgression detection in the presence of purifying selection. Systematic Biology 75:752–775.

He, C., D. Liang, and P. Zhang. 2020. Asymmetric distribution of gene trees can arise under purifying selection if differences in population size exist. Molecular Biology and Evolution 37:881–892.

Hejase, H. A. and K. J. Liu. 2016. A scalability study of phylogenetic network inference methods using empirical datasets and simulations involving a single reticulation. BMC Bioinformatics 17:422.

Holtgrefe, N., K. T. Huber, L. van Iersel, M. Jones, S. Martin, and V. Moulton. 2025. Squirrel: Reconstructing semi-directed phylogenetic level-1 networks from four-leaved networks or sequence alignments. Molecular Biology and Evolution 42:msaf067.

Huber, K. T., V. Moulton, C. Semple, and T. Wu. 2018. Quarnet inference rules for level-1 networks. Bulletin of Mathematical Biology 80:2137–2153.

Hubert, L., P. Arabie, and J. Meulman. 1998. Graph-theoretic representations for proximity matrices through strongly-anti-robinson or circular strongly-anti-robinson matrices. Psychometrika 63:341–358.

Huson, D. H., R. Rupp, and C. Scornavacca. 2010. Phylogenetic Networks: Concepts, Algorithms and Applications. Cambridge University Press.

Jaffe, A., N. Amsel, Y. Aizenbud, B. Nadler, J. T. Chang, and Y. Kluger. 2021. Spectral neighbor joining for reconstruction of latent tree models. SIAM Journal on Mathematics of Data Science 3:113–141.

Jarvis, E. D., S. Mirarab, A. J. Aberer, B. Li, P. Houde, C. Li, S. Y. W. Ho, B. C. Faircloth, B. Nabholz, J. T. Howard, A. Suh, C. C. Weber, R. R. da Fonseca, J. Li, F. Zhang, H. Li, L. Zhou, N. Narula, L. Liu, G. Ganapathy, B. Boussau, M. S. Bayzid, V. Zavidovych, S. Subramanian, T. Gabaldón, S. Capella-Gutiérrez, J. Huerta-Cepas, B. Rekepalli, K. Munch, M. Schierup, B. Lindow, W. C. Warren,D. Ray, R. E. Green, M. W. Bruford, X. Zhan, A. Dixon, S. Li, N. Li, Y. Huang, E. P. Derryberry, M. F. Bertelsen, F. H. Sheldon, R. T. Brumfield, C. V. Mello, P. V. Lovell, M. Wirthlin, M. P. C. Schneider, F. Prosdocimi, J. Samaniego, A. M. Vargas Velazquez, A. Alfaro-Núñez, P. F. Campos, B. Petersen, T. Sicheritz-Ponten, A. Pas, T. Bailey, P. Scofield, M. Bunce, D. M. Lambert, Q. Zhou, P. Perelman,A. C. Driskell, B. Shapiro, Z. Xiong, Y. Zeng, S. Liu, Z. Li, B. Liu, K. Wu, J. Xiao, X. Yinqi, Q. Zheng, Y. Zhang, H. Yang, J. Wang,L. Smeds, F. E. Rheindt, M. Braun, J. Fjeldsa, L. Orlando, F. K. Barker, K. A. Jønsson, W. Johnson, K.-P. Koepfli, S. O’Brien,D. Haussler, O. A. Ryder, C. Rahbek, E. Willerslev, G. R. Graves, T. C. Glenn, J. McCormack, D. Burt, H. Ellegren, P. Alström,S. V. Edwards, A. Stamatakis, D. P. Mindell, J. Cracraft, E. L. Braun, T. Warnow, W. Jun, M. T. P. Gilbert, and G. Zhang. 2014. Whole-genome analyses resolve early branches in the tree of life of modern birds. Science 346:1320–1331.

Jermiin, L. S., S. Y. Ho, F. Ababneh, J. Robinson, and A. W. Larkum. 2004. The biasing effect of compositional heterogeneity on phylogenetic estimates may be underestimated. Systematic Biology 53:638–643.

Justison, J. A., C. Solis-Lemus, and T. A. Heath. 2023. Siphynetwork: An r package for simulating phylogenetic networks. Methods in Ecology and Evolution 14:1687–1698.

Karoui, N. E. and H.-T. Wu. 2016. Graph connection Laplacian methods can be made robust to noise. The Annals of Statistics 44:346–372.

Kendall, M. G. 1938. A new measure of rank correlation. Biometrika 30:81–93.

Kolbow, N., S. Kong, and C. Solís-Lemus. 2026. A method for massively scalable inference of phylogenetic networks. bioRxiv .

Kong, S., C. Solís-Lemus, and G. P. Tiley. 2025. Phylogenetic networks empower biodiversity research. Proc Natl Acad Sci U S A 122:e2410934122.

Kück, P., J. Romahn, and K. Meusemann. 2022. Pitfalls of the site-concordance factor (scf) as measure of phylogenetic branch support.NAR Genom Bioinform 4:qac064.

Leebens-Mack, J. H., M. S. Barker, E. J. Carpenter, M. K. Deyholos, M. A. Gitzendanner, S. W. Graham, I. Grosse, Z. Li, M. Melkonian,S. Mirarab, M. Porsch, M. Quint, S. A. Rensing, D. E. Soltis, P. S. Soltis, D. W. Stevenson, K. K. Ullrich, N. J. Wickett, L. DeGironimo,P. P. Edger, I. E. Jordon-Thaden, S. Joya, T. Liu, B. Melkonian, N. W. Miles, L. Pokorny, C. Quigley, P. Thomas, J. C. Villarreal, M. M. Augustin, M. D. Barrett, R. S. Baucom, D. J. Beerling, R. M. Benstein, E. Biffin, S. F. Brockington, D. O. Burge, J. N. Burris, K. P. Burris, V. Burtet-Sarramegna, A. L. Caicedo, S. B. Cannon, Z. Çebi, Y. Chang, C. Chater, J. M. Cheeseman, T. Chen, N. D. Clarke,H. Clayton, S. Covshoff, B. J. Crandall-Stotler, H. Cross, C. W. dePamphilis, J. P. Der, R. Determann, R. C. Dickson, V. S. Di Stilio,S. Ellis, E. Fast, N. Feja, K. J. Field, D. A. Filatov, P. M. Finnegan, S. K. Floyd, B. Fogliani, N. García, G. Gâteblé, G. T. Godden,F. Q. Y. Goh, S. Greiner, A. Harkess, J. M. Heaney, K. E. Helliwell, K. Heyduk, J. M. Hibberd, R. G. J. Hodel, P. M. Hollingsworth,M. T. J. Johnson, R. Jost, B. Joyce, M. V. Kapralov, E. Kazamia, E. A. Kellogg, M. A. Koch, M. Von Konrat, K. Könyves, T. M. Kutchan,V. Lam, A. Larsson, A. R. Leitch, R. Lentz, F.-W. Li, A. J. Lowe, M. Ludwig, P. S. Manos, E. Mavrodiev, M. K. McCormick, M. McKain,T. McLellan, J. R. McNeal, R. E. Miller, M. N. Nelson, Y. Peng, P. Ralph, D. Real, C. W. Riggins, M. Ruhsam, R. F. Sage, A. K. Sakai,M. Scascitella, E. E. Schilling, E.-M. Schlösser, H. Sederoff, S. Servick, E. B. Sessa, A. J. Shaw, S. W. Shaw, E. M. Sigel, C. Skema,A. G. Smith, A. Smithson, C. N. Stewart, J. R. Stinchcombe, P. Szövényi, J. A. Tate, H. Tiebel, D. Trapnell, M. Villegente, C.-N. Wang, S. G. Weller, M. Wenzel, S. Weststrand, J. H. Westwood, D. F. Whigham, S. Wu, A. S. Wulff, Y. Yang, D. Zhu, C. Zhuang,J. Zuidof, M. W. Chase, J. C. Pires, C. J. Rothfels, J. Yu, C. Chen, L. Chen, S. Cheng, J. Li, R. Li, X. Li, H. Lu, Y. Ou, X. Sun, X. Tan,J. Tang, Z. Tian, F. Wang, J. Wang, X. Wei, X. Xu, Z. Yan, F. Yang, X. Zhong, F. Zhou, Y. Zhu, Y. Zhang, S. Ayyampalayam, T. J. Barkman, N.-p. Nguyen, N. Matasci, D. R. Nelson, E. Sayyari, E. K. Wafula, R. L. Walls, T. Warnow, H. An, N. Arrigo, A. E. Baniaga,S. Galuska, S. A. Jorgensen, T. I. Kidder, H. Kong, P. Lu-Irving, H. E. Marx, X. Qi, C. R. Reardon, B. L. Sutherland, G. P. Tiley, S. R. Welles, R. Yu, S. Zhan, L. Gramzow, G. Theißen, G. K.-S. Wong, and O. T. P. T. Initiative. 2019. One thousand plant transcriptomes and the phylogenomics of green plants. Nature 574:679–685.

Lubbe, P., N. J. Rawlence, O. Kardailsky, B. C. Robertson, R. Day, M. Knapp, and N. Dussex. 2022. Mitogenomes resolve the phylogeography and divergence times within the endemic new zealand callaeidae (aves: Passerida). Zoological Journal of the Linnean Society 196:1451–1463.

Mackiewicz, P., A. D. Urantówka, A. Kroczak, and D. Mackiewicz. 2019. Resolving phylogenetic relationships within passeriformes based on mitochondrial genes and inferring the evolution of their mitogenomes in terms of duplications. Genome Biology and Evolution 11:2824–2849.

Meurer, A., C. P. Smith, M. Paprocki, O. Čertík, S. B. Kirpichev, M. Rocklin, A. Kumar, S. Ivanov, J. K. Moore, S. Singh, T. Rathnayake,S. Vig, B. E. Granger, R. P. Muller, F. Bonazzi, H. Gupta, S. Vats, F. Johansson, F. Pedregosa, M. J. Curry, A. R. Terrel, Š. Roučka,A. Saboo, I. Fernando, S. Kulal, R. Cimrman, and A. Scopatz. 2017. Sympy: symbolic computing in python. PeerJ Computer Science 3:e103.

Mirarab, S., L. Nakhleh, and T. Warnow. 2021. Multispecies Coalescent: Theory and Applications in Phylogenetics. Annual Review of Ecology, Evolution, and Systematics 52:247–268.

Mirarab, S., R. Reaz, M. S. Bayzid, T. Zimmermann, M. S. Swenson, and T. Warnow. 2014. Astral: genome-scale coalescent-based species tree estimation. Bioinformatics 30:i541–i548.

Mitchell, J. D., E. S. Allman, and J. A. Rhodes. 2019. Hypothesis testing near singularities and boundaries. Electron J Stat 13:2150– 2193.

Moyle, R. G., C. H. Oliveros, M. J. Andersen, P. A. Hosner, B. W. Benz, J. D. Manthey, S. L. Travers, R. M. Brown, and B. C. Faircloth.2016. Tectonic collision and uplift of wallacea triggered the global songbird radiation. Nature Communications 7:12709.

Nakhleh, L. 2011. Evolutionary Phylogenetic Networks: Models and Issues Pages 125–158. Springer US, Boston, MA.

Nickrent, D. L., C. L. Parkinson, J. D. Palmer, and R. J. Duff. 2000. Multigene phylogeny of land plants with special reference to bryophytes and the earliest land plants. Mol Biol Evol 17:1885–1895.

Ottenburghs, J. 2020. Ghost introgression: Spooky gene flow in the distant past. Bioessays 42:e2000012.

Powell, A. F. L. A. and J. S. Kirkley. 2021. An instance of hybridization with common grackle (quiscalus quiscula) and male parental care by great-tailed grackle (q. mexicanus). The Wilson Journal of Ornithology 133:73–81.

Price, M. N., P. S. Dehal, and A. P. Arkin. 2010. Fasttree 2 – approximately maximum-likelihood trees for large alignments. PLOS ONE 5:1–10.

Recanati, A., T. Kerdreux, and A. d’Aspremont. 2018. Reconstructing latent orderings by spectral clustering. CoRR abs/1807.07122.

Reshef, O., O. Glassman, O. Zuk, Y. Aizenbud, B. Nadler, and A. Jaffe. 2026. Sdsr: A spectral divide-and-conquer approach for species tree reconstruction.

Rhodes, J. A., H. Baños, J. D. Mitchell, and E. S. Allman. 2021. Mscquartets 1.0: quartet methods for species trees and networks under the multispecies coalescent model in r. Bioinformatics 37:1766–1768.

Robinson, D. and L. Foulds. 1981. Comparison of phylogenetic trees. Mathematical Biosciences 53:131–147.

Sayyari, E. and S. Mirarab. 2016. Anchoring quartet-based phylogenetic distances and applications to species tree reconstruction. BMC Genomics 17:783.

Sigel, E. M., M. D. Windham, C. H. Haufler, and K. M. Pryer. 2014a. Phylogeny, divergence time estimates, and phylogeography of the diploid species of the polypodium vulgare complex (polypodiaceae). Systematic Botany 39:1042–1055.

Sigel, E. M., M. D. Windham, and K. M. Pryer. 2014b. Evidence for reciprocal origins in polypodium hesperium (polypodiaceae): a fern model system for investigating how multiple origins shape allopolyploid genomes. Am J Bot 101:1476–1485.

Solís-Lemus, C. and C. Ané. 2016. Inferring phylogenetic networks with maximum pseudolikelihood under incomplete lineage sorting.PLOS Genetics 12:1–21.

Solís-Lemus, C., M. Yang, and C. Ané. 2016. Inconsistency of species tree methods under gene flow. Systematic Biology 65:843–851.

Stephan, L. and L. Massoulié. 2019. Robustness of spectral methods for community detection. Pages 2831–2860 in Proceedings of the Thirty-Second Conference on Learning Theory (A. Beygelzimer and D. Hsu, eds.) vol. 99 of Proceedings of Machine Learning Research PMLR.

Stiller, J., S. Feng, A.-A. Chowdhury, I. Rivas-González, D. A. Duchêne, Q. Fang, Y. Deng, A. Kozlov, A. Stamatakis, S. Claramunt,J. M. T. Nguyen, S. Y. W. Ho, B. C. Faircloth, J. Haag, P. Houde, J. Cracraft, M. Balaban, U. Mai, G. Chen, R. Gao, C. Zhou, Y. Xie,Z. Huang, Z. Cao, Z. Yan, H. A. Ogilvie, L. Nakhleh, B. Lindow, B. Morel, J. Fjeldså, P. A. Hosner, R. R. da Fonseca, B. Petersen, J. A. Tobias, T. Székely, J. D. Kennedy, A. H. Reeve, A. Liker, M. Stervander, A. Antunes, D. T. Tietze, M. F. Bertelsen, F. Lei, C. Rahbek,G. R. Graves, M. H. Schierup, T. Warnow, E. L. Braun, M. T. P. Gilbert, E. D. Jarvis, S. Mirarab, and G. Zhang. 2024. Complexity of avian evolution revealed by family-level genomes. Nature 629:851–860.

Stull, G. W., K. K. Pham, P. S. Soltis, and D. E. Soltis. 2023. Deep reticulation: the long legacy of hybridization in vascular plant evolution. The Plant Journal 114:743–766.

Suh, A. 2016. The phylogenomic forest of bird trees contains a hard polytomy at the root of neoaves. Zoologica Scripta 45:50–62.

von Luxburg, U. 2007. A tutorial on spectral clustering. CoRR abs/0711.0189.

Wen, D., Y. Yu, J. Zhu, and L. Nakhleh. 2018. Inferring phylogenetic networks using phylonet. Systematic Biology 67:735–740.

Wickett, N. J., S. Mirarab, N. Nguyen, T. Warnow, E. Carpenter, N. Matasci, S. Ayyampalayam, M. S. Barker, J. G. Burleigh, M. A. Gitzendanner, B. R. Ruhfel, E. Wafula, J. P. Der, S. W. Graham, S. Mathews, M. Melkonian, D. E. Soltis, P. S. Soltis, N. W. Miles,C. J. Rothfels, L. Pokorny, A. J. Shaw, L. DeGironimo, D. W. Stevenson, B. Surek, J. C. Villarreal, B. Roure, H. Philippe, C. W. dePamphilis, T. Chen, M. K. Deyholos, R. S. Baucom, T. M. Kutchan, M. M. Augustin, J. Wang, Y. Zhang, Z. Tian, Z. Yan, X. Wu,X. Sun, G. K.-S. Wong, and J. Leebens-Mack. 2014. Phylotranscriptomic analysis of the origin and early diversification of land plants. Proceedings of the National Academy of Sciences 111:E4859–E4868.

Willson, J. and T. Warnow. 2026. Camus: scalable phylogenetic network estimation. Bioinformatics 42:btag245.

Wong, T. K. F., N. Ly-Trong, H. Ren, P. Demotte, H. Baños, A. J. Roger, E. Susko, C. Bielow, N. De Maio, N. Goldman, M. W. Hahn,M. dos Reis, L. S. Vinh, G. Huttley, R. Lanfear, and B. Q. Minh. 2026. Iq-tree 3: phylogenomic inference software using complex evolutionary models. Molecular Biology and Evolution 43:msag117.

Wu, Y. 2012. Coalescent-based species tree inference from gene tree topologies under incomplete lineage sorting by maximum likelihood. Evolution 66:763–775.

Yu, Y., J. H. Degnan, and L. Nakhleh. 2012. The probability of a gene tree topology within a phylogenetic network with applications to hybridization detection. PLOS Genetics 8:1–10.

Yu, Y. and L. Nakhleh. 2015. A maximum pseudo-likelihood approach for phylogenetic networks. BMC Genomics 16:S10.

Zhang, C., R. Nielsen, and S. Mirarab. 2025. ASTER: A Package for Large-Scale Phylogenomic Reconstructions. Molecular Biology and Evolution 42:msaf172.eprint: https://academic.oup.com/mbe/article-pdf/42/8/msaf172/63779203/msaf172.pdf.

Zhu, J. and L. Nakhleh. 2018. Inference of species phylogenies from bi-allelic markers using pseudo-likelihood. Bioinformatics 34:i376– i385.

