## Supplementary Materials for "BROOQS: Quartet-based Resolution of Level-1 Hybridization Cycles without Tests of Symmetry"

### Supplementary Material

#### A Supplementary Figures and Tables

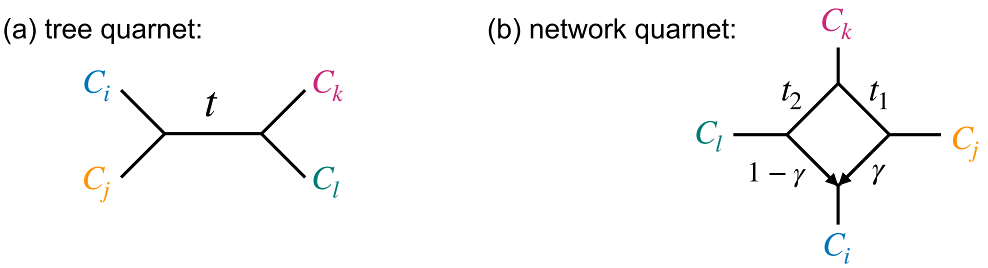


$$CF(C_iC_j|C_kC_l) = 1 - \frac{2}{3}e^{-t} \qquad CF(C_iC_j|C_kC_l) = \gamma(1 - \frac{2}{3}e^{-t_1}) + (1-\gamma)(\frac{1}{3}e^{-t_2})$$

$$CF(C_iC_k|C_jC_l) = \frac{1}{3}e^{-t} \qquad CF(C_iC_k|C_jC_l) = \gamma(\frac{1}{3}e^{-t_1}) + (1-\gamma)(\frac{1}{3}e^{-t_2})$$

$$CF(C_iC_l|C_jC_k) = \frac{1}{3}e^{-t} \qquad CF(C_iC_l|C_jC_k) = \gamma(\frac{1}{3}e^{-t_1}) + (1-\gamma)(1 - \frac{2}{3}e^{-t_2})$$

**Figure S1** The two classes of metric quarnets on four distinct components $C_i, C_j, C_k, C_l \in \mathcal{C}$. **(a)** The induced metric is a *tree quarnet* if $C_i, C_j, C_k, C_l$ are all non-hybrid components. In the example shown, $CF(C_i, C_j \mid C_k, C_l) > CF(C_i, C_k \mid C_j, C_l) = CF(C_i, C_l \mid C_j, C_k)$. **(b)** The induced metric is a *network quarnet* if one component is a hybrid component. In the example shown, $C_i$ is the hybrid component and $CF(C_i, C_j \mid C_k, C_l) > CF(C_i, C_k \mid C_j, C_l)$, $CF(C_i, C_l \mid C_j, C_k) > CF(C_i, C_k \mid C_j, C_l)$.

**Table S1** Summary statistics for each of the simulation datasets. For each number of taxa $n$, average number of true blobs, average size of each blob and maximum blob size is shown.

| $n$ | Mean num | Mean size | Max size |
|---|---|---|---|
| 15 | 1.09 | 6.72 | 10 |
| 25 | 1.04 | 7.95 | 14 |
| 50 | 1.22 | 9.73 | 18 |
| 100 | 1.64 | 10.7 | 20 |
| 150 | 2.62 | 11.0 | 21 |
| 200 | 2.06 | 13.6 | 26 |

(a) Fixed-ILS dataset

| $n$ | Mean num | Mean size | Max size |
|---|---|---|---|
| 50 | 1.50 | 7.20 | 17 |
| 100 | 1.80 | 10.5 | 22 |
| 200 | 2.10 | 14.2 | 24 |

(b) Varying-ILS dataset

| $n$ | Mean num | Mean size | Max size |
|---|---|---|---|
| 25 | 1.55 | 7.26 | 13 |
| 50 | 2.15 | 8.72 | 18 |
| 100 | 1.85 | 10.9 | 28 |
| 200 | 2.28 | 13.2 | 24 |

(c) Varying-ILS+GTEE dataset

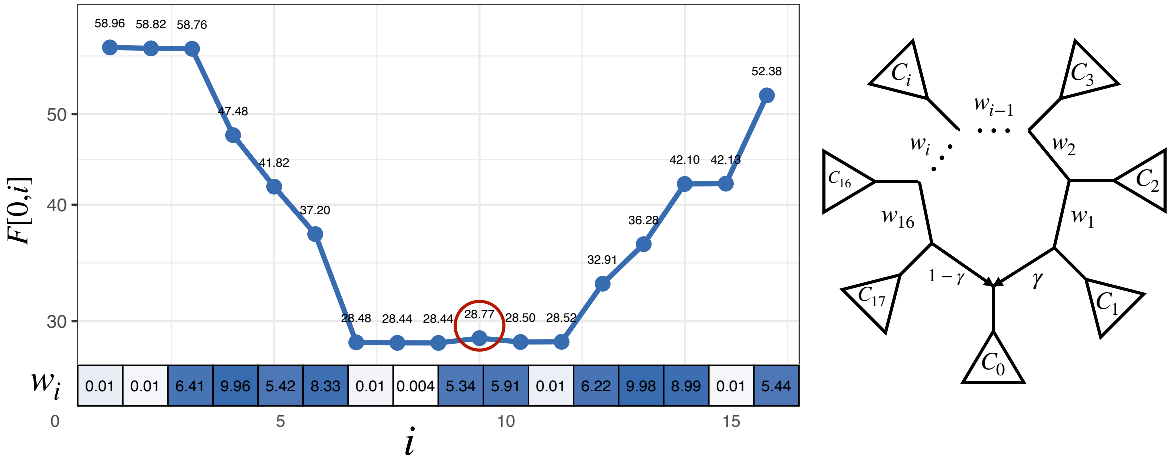


**Figure S2** A counterexample violating Theorem 1. Consider the 18-cycle on the right with the true ordering $\pi^*(\mathcal{C}) = [C_0, C_1, \ldots, C_{16}, C_{17}]$. Theorem 1 states that each row and column of the correctly permuted F-matrix $\pi^*(F)$ must be unimodal; i.e., decreasing up to a minimum and increasing afterward. This property can be violated for cycles of size 16 or more (Table S2), for the hybrid row $[F[0,1], F[0,2], \ldots, F[0,d-1]]$ under some choice of parameters. The panel on the left shows an example of this violation for the 18-cycle on the right, where $F[0,10]$ is a local maximum (marked in red), violating unimodality. The branch lengths $w_i$ in this example vary drastically, ranging from $0.004$ to $\sim 10$, creating a challenging case. For this example, the inheritance probability is $\gamma = 0.5759$.

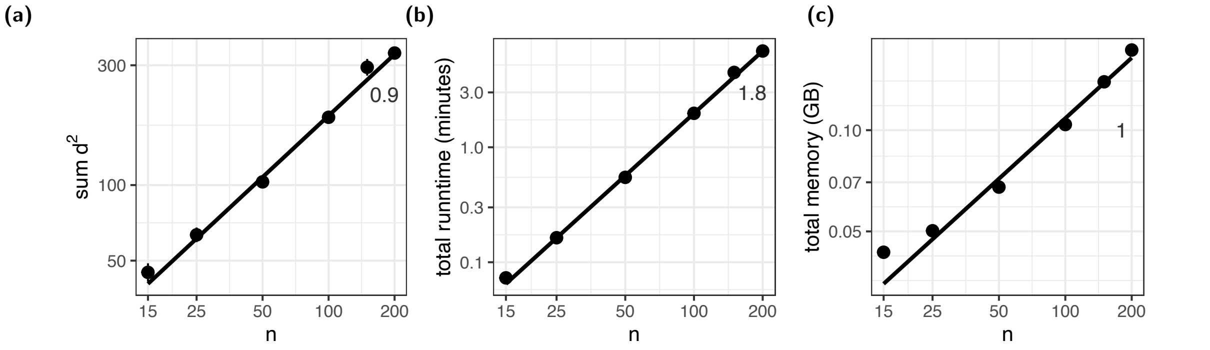


**Figure S3** Total complexity of BROOQS. The total running time of BROOQS is $O(mnD)$ where $D = \sum_c d_c^2$ is the sum of squared blob degrees. The empirical value of $D$ **(a)** and the running time of BROOQS **(b)** are shown, as well as the slope of the line in log-log scale (slope is estimated for $n \geq 50$). **(c)** shows empirical memory complexity. Empirical analysis merges all simulated datasets and both true and estimated gene trees for each dataset, except for the Varying ILS+GTEE dataset, where only true gene trees are included (the number of estimated gene trees in this dataset varies drastically across replicates).

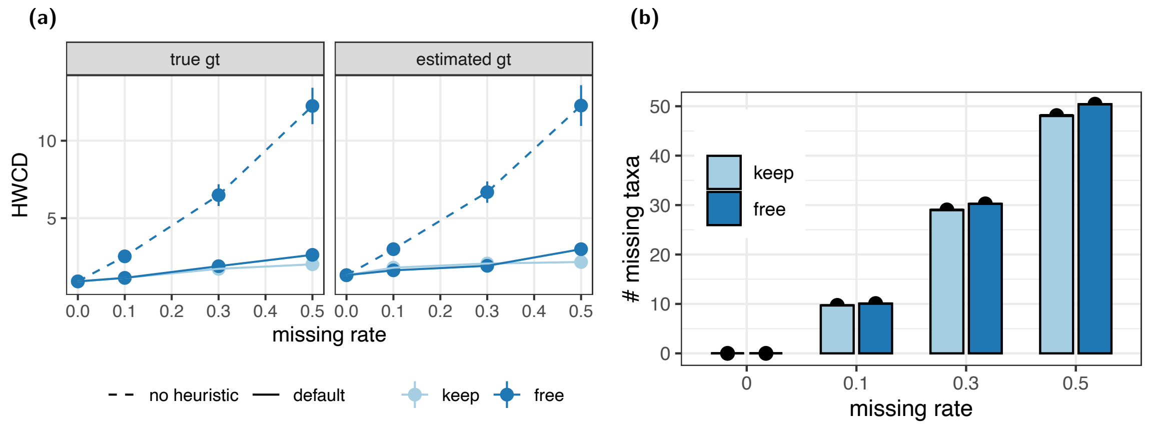


**Figure S4** Fixed-ILS dataset ($n = 100$). For $p \in \{0, 0.1, 0.3, 0.5\}$, we removed taxa from the true and estimated gene trees at random in two ways: removing each taxa with probability $p$ (free), and removing each taxa with probability $p$ but ensuring at least one taxon from each component is included in every gene tree (keep). We then ran BROOQS-Spectral on the pruned gene trees, and handled missing components using the heuristic described in Section B.5.1. To evaluate our heuristic for the free mode, we also ran BROOQS-Spectral once without any heuristics to handle missing components. We show estimation error as HWCD **(a)** and average number of missing taxa per gene tree **(b)** for each rate $p$. While the keep mode has more constraints for removing taxa, both modes prune similar numbers of taxa from the gene trees, confirming that the difference in the estimation error is due to the heuristics used to handle missing components.

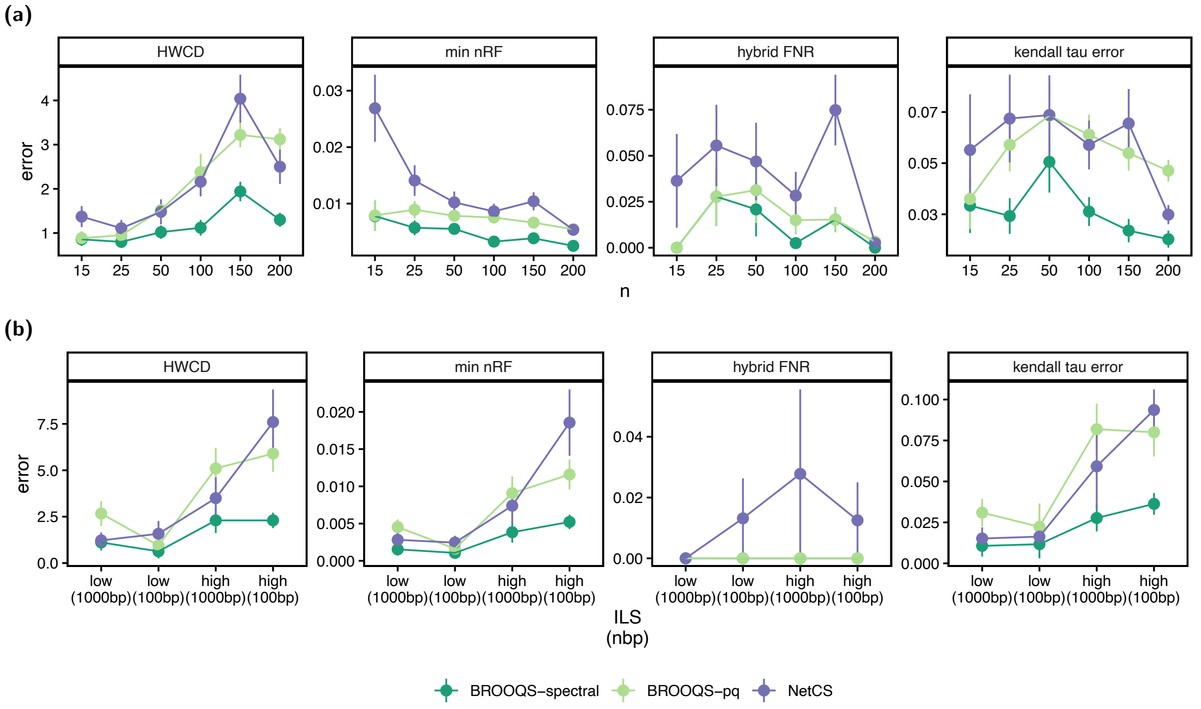


**Figure S5** Experiment 1. Comparing BROOQS to NetCS on the Fixed-ILS **(a)** and Varying-ILS+GTEE **(b)** datasets on true tree of blobs. Panels show four error metrics: hard-wired cluster distance (HWCD), min nRF, False negative rate of hybrid component (FNR), and kendal-tau error. The Varying-ILS+GTEE dataset **(b)** is limited to $n = 200$ (10 replicates per model condition). See Figure 2 for the comparison on smaller sizes.

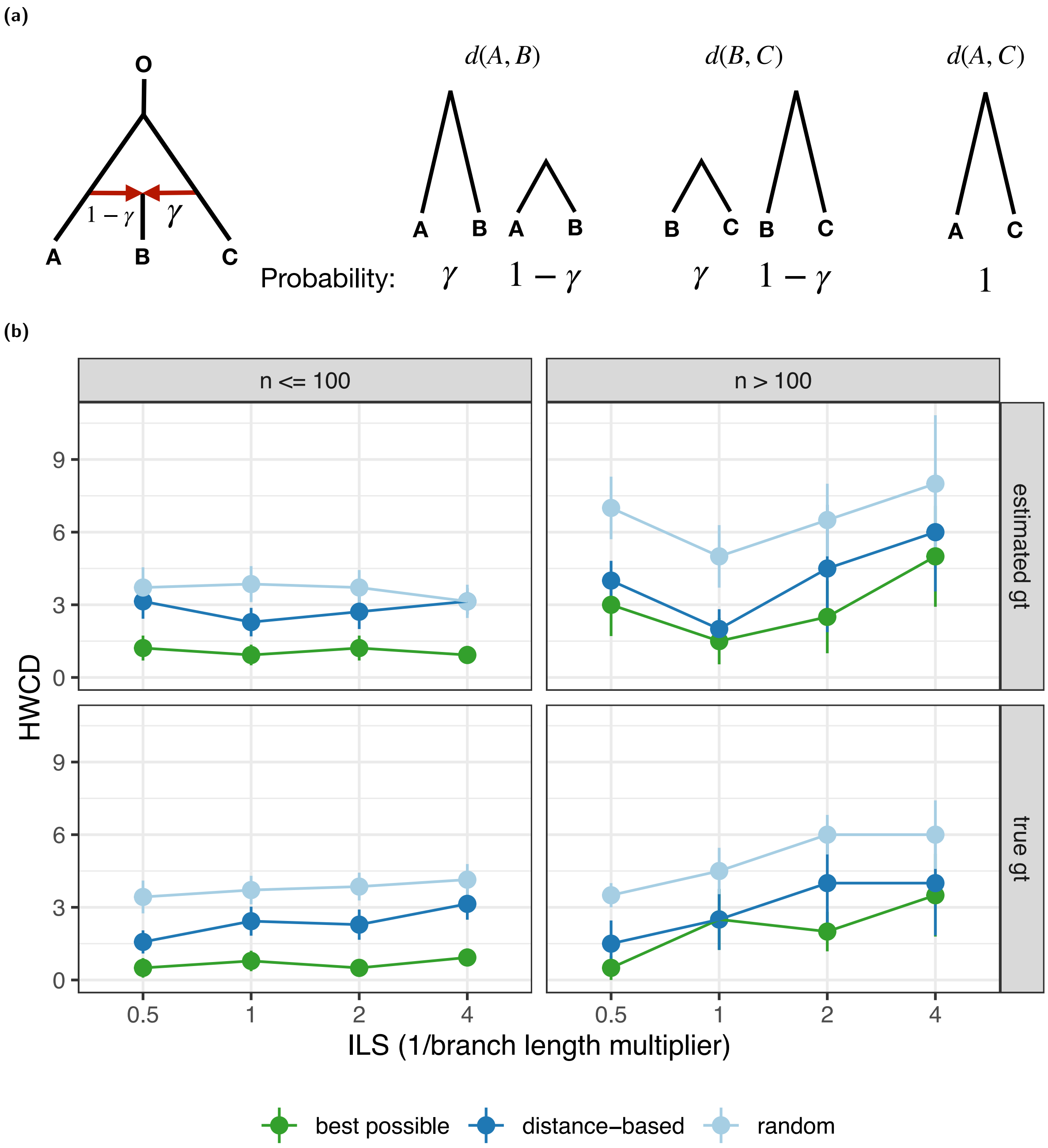


**Figure S6** **(a)** For the 4-cycle on the left, we show all probable rooted topologies for each pair. Any pair containing the hybrid taxon **B** displays two rooted topologies with different pairwise distances. The pair not containing the hybrid (**A**, **C**) is the only pair with one rooted topology. Therefore, while $d(A, B)$ and $d(B, C)$ are drawn from two different distributions, and therefore are expected to have higher variance across a set of gene trees, $d(A, C)$ is drawn from one distribution. Note that these illustrations and conclusions completely ignore rate variation across gene trees. **(b)** Varying-ILS dataset. Evaluating the distance-based heuristic for identifying the hybrid component in 4-cycles. We only show replicates with at least one 4-cycle, and evaluate against random hybrid assignment (the default setting in BROOQS). For the heuristic mode, we sample $50$ triplets randomly, and choose the hybrid component by majority voting. We use the true 4-cycle as the baseline (referred to as best possible in the legend). Note that the baseline error is not zero, as many replicates contain other cycles of degree $5$ or more, that are resolved independently by BROOQS and contribute to the overall error.

**(a)**

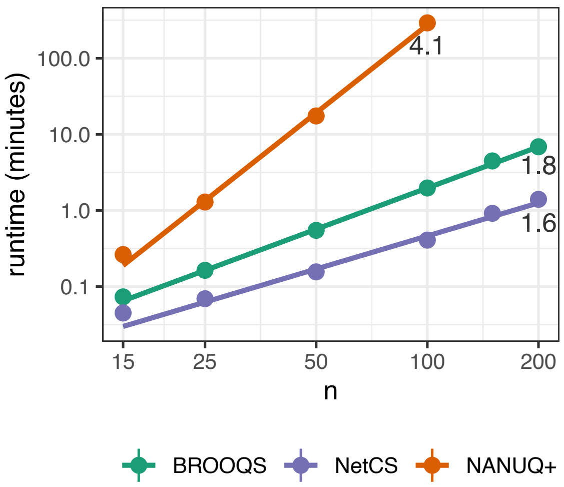


**(b)**

| Method | Runtime (minutes) | Memory (GB) |
|---|---|---|
| TOB-QMC+BROOQS | 53.89 | 0.104 |
| TOB-QMC+NetCS | 53.56 | 0.405 |
| NANUQ+ | 296 | 14.7 |
| ASTRAL+CAMUS | 118 | 1.27 |
| SQUIRREL | 767 | 24.9 |

**Figure S7** **(a)** Experiment 1. Comparing the running time of BROOQS to other blob resolution methods, NetCS and NANUQ+. Both axes are shown on a logarithmic scale, as well as the slope of the fitted line. Since both modes of BROOQS have the same runtime and memory complexity, we only show BROOQS-Spectral. Each method was given the true tree of blobs; therefore, the running time and memory usage of the ToB estimation step are not included for any of the methods. Note that NANUQ+ requires the output from TINNIK to resolve the tree of blobs, even when the true tree of blobs is provided as input. Thus, the running time of TINNIK is included for NANUQ+. Under the 24-hour and 50-GB memory constraints, NANUQ+ finished for $n \leq 100$, while both BROOQS and NetCS ran for all numbers of taxa. For each $n$, true and estimated gene trees across all replicates from all three simulated datasets are merged. For the Varying ILS+GTEE dataset, only true gene trees are included (the number of estimated gene trees in this dataset varies drastically across replicates). **(b)** Experiment 2 ($n = 100$). Comparing running time and memory of all evaluated methods for full network inference pipeline. Running times of tree of blobs estimation (TOB-QMC for BROOQS and NetCS, and TINNIK for NANUQ+), and ASTRAL tree inference (only for CAMUS) are added to total running times of the corresponding methods. For each $n$, true and estimated gene trees across all replicates from all three simulated datasets are merged. For the Varying ILS+GTEE dataset, only true gene trees are included (the number of estimated gene trees in this dataset varies drastically across replicates). $n = 100$ was the largest dataset, for which all methods successfully finished within our constraints.

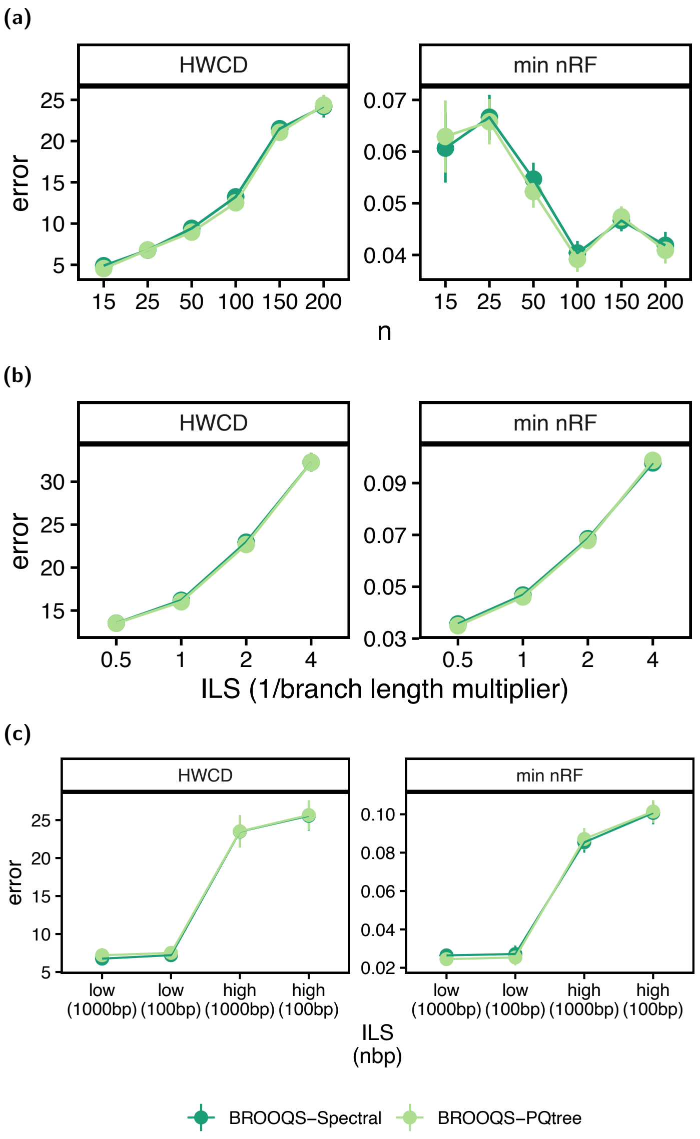


**Figure S8** Experiment 2. Comparing BROOQS-Spectral and BROOQS-PQtree on the TOB-QMC estimated tree of blobs on **(a)** fixed-ILS, **(b)** varying-ILS, and **(c)** varying-ILS+GTEE datasets. Two error metrics HWCD and min nRF are shown.

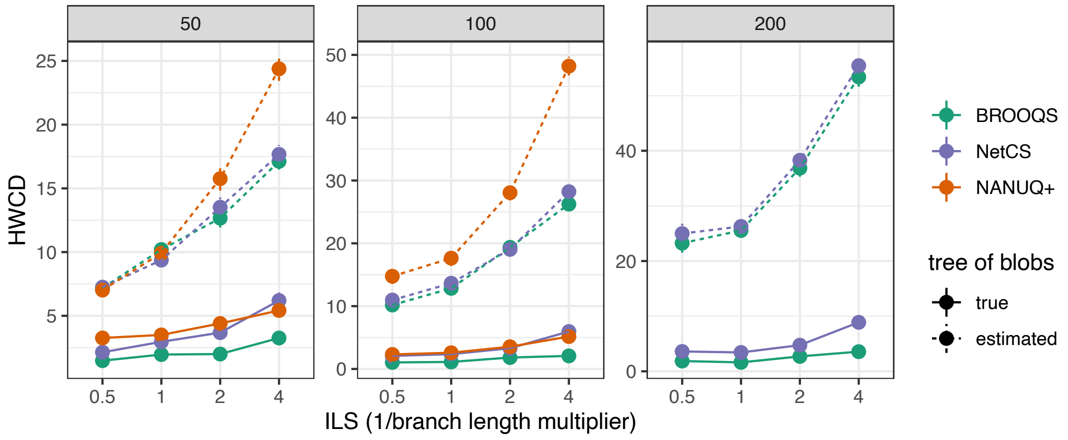


**Figure S9** Experiment 2 (Varying-ILS dataset). Comparing the network estimation error of the three methods, for true vs. estimated tree of blobs as input. The difference between the dotted line and the solid lines is an indication of how much of the error comes from the identification of TOB versus its resolution. BROOQS and NetCS use the TOB-QMC estimated tree of blobs, whereas NANUQ+ uses TINNIK's tree of blobs. The performance of all methods is heavily affected by the construction of the tree of blobs.

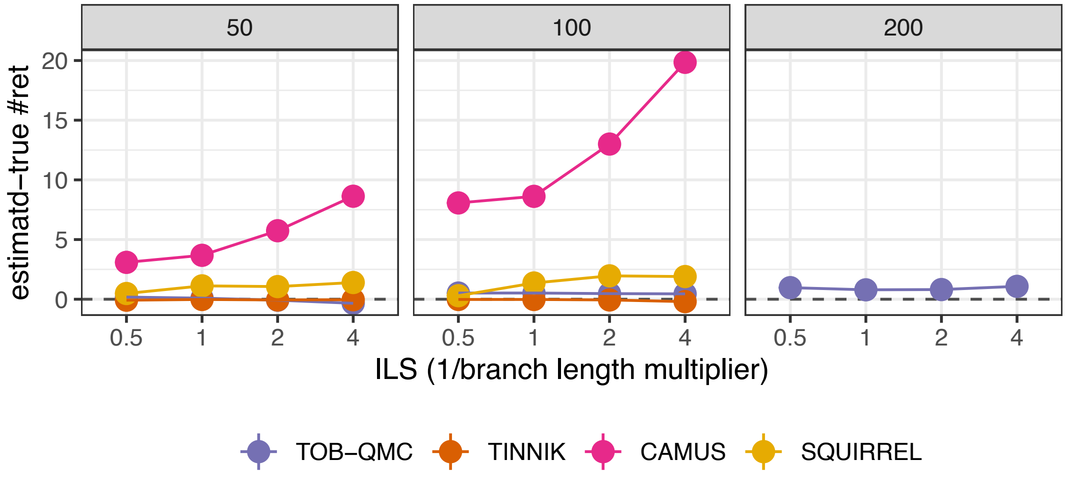


**Figure S10** Experiment 2 (Varying-ILS dataset). Comparing number of estimated − true reticulation events for two tree of blobs estimation methods TOB-QMC and TINNIK, as well as two standalone network inference tools CAMUS and SQUIRREL. Above zero (dashed line) means overestimation of reticulation events. The tree of blobs estimated by TOB-QMC is used as input to both BROOQS and NetCS; therefore, these methods share the same number of reticulations. Similarly, the tree of blobs estimated by TINNIK is used by NANUQ+. CAMUS severely overestimates the number of reticulations; therefore, in our analyses we pick the CAMUS network with the same number of reticulations as TOB-QMC to give CAMUS an advantage.

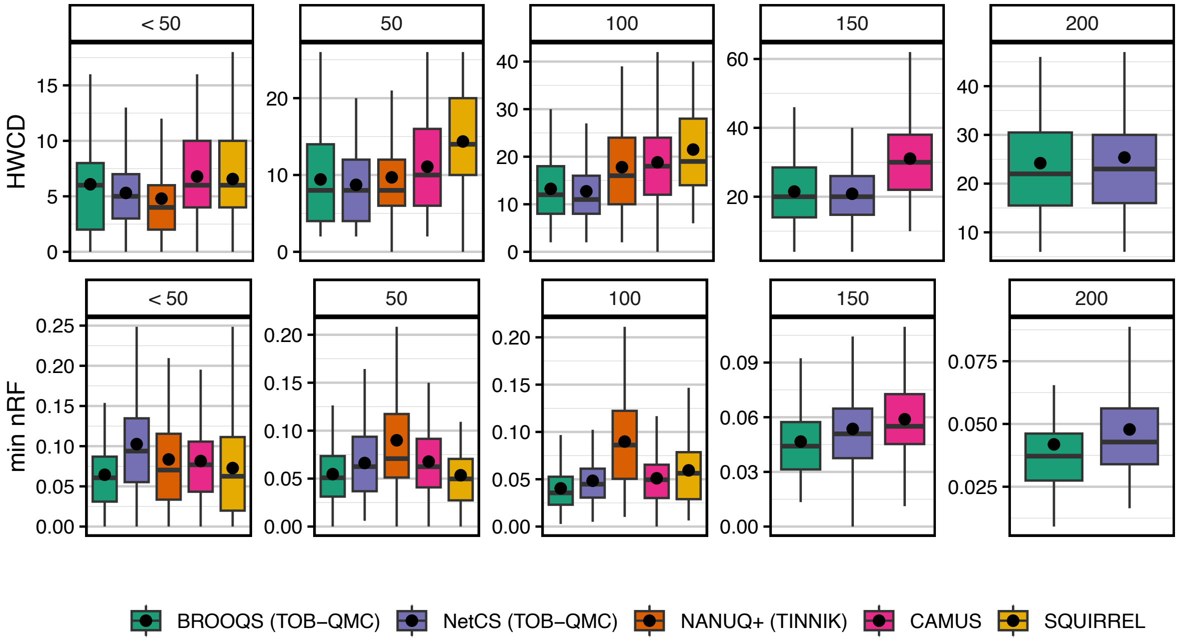


**Figure S11** Experiment 2 (Fixed-ILS dataset). Comparing BROOQS on the estimated tree of blobs (TOB-QMC) with other network estimation methods. Methods are compared using two error metrics, HWCD and min nRF. Panels show the number of taxa ($n$). Under the 24-hour and 50-GB memory constraints, SQUIRREL and NANUQ+ finished for $n \leq 100$, while CAMUS finished for $n \leq 150$. BROOQS and NetCS ran for all numbers of taxa.

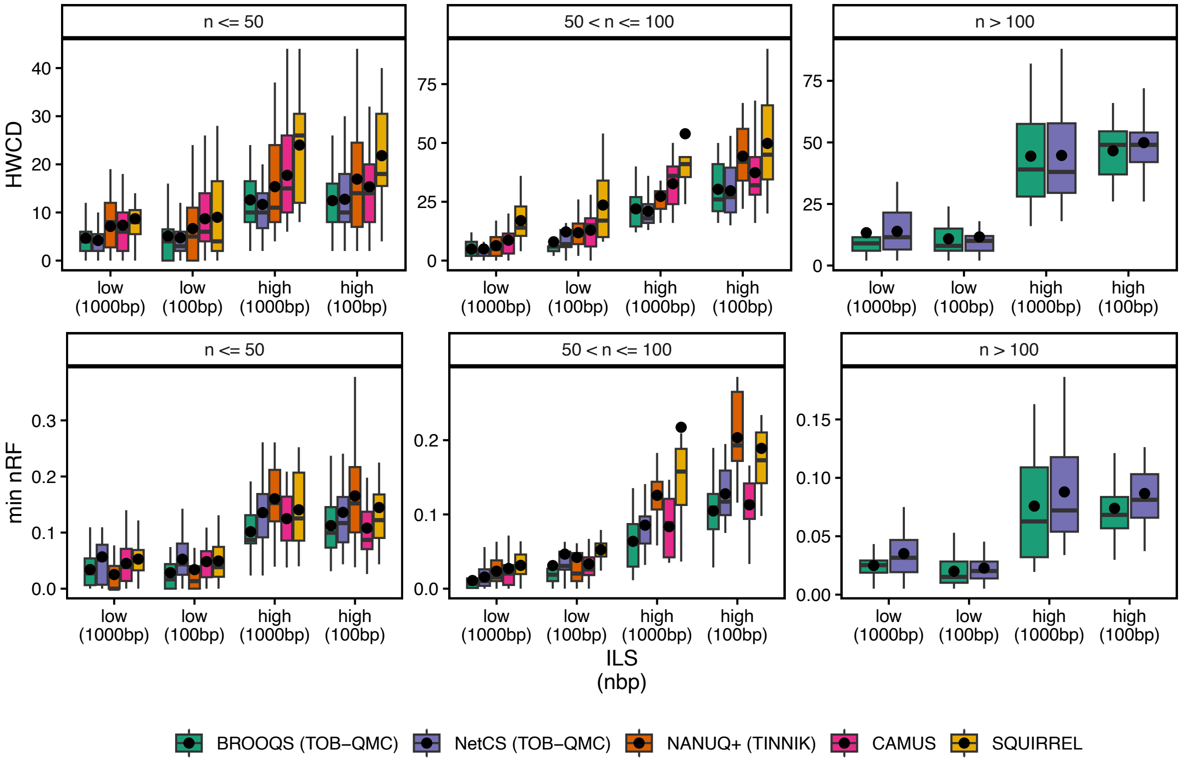


**Figure S12** Experiment 2 (Varying-ILS+GTEE dataset). Comparing BROOQS on the estimated tree of blobs (TOB-QMC) with other network estimation methods. x-axis shows increased complexity. Methods are compared using two error metrics, HWCD and min nRF. Panels show the number of taxa ($n$). Under the 24-hour and 50-GB memory constraints, SQUIRREL, CAMUS, and NANUQ+ finished for $n \leq 100$. BROOQS and NetCS ran for all numbers of taxa.

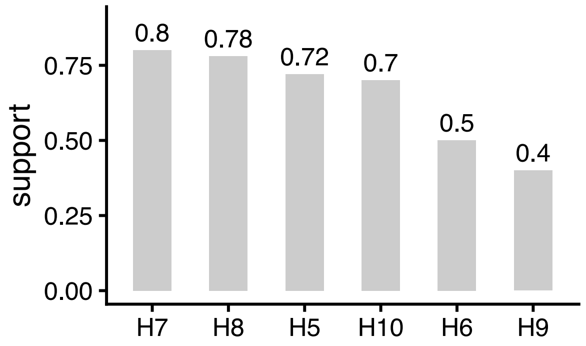


**Figure S13** Avian dataset. Support for the hybrid component for all the 4-cycles inferred by TOB-QMC+BROOQS. Support is computed as the proportion of the $50$ sampled triplets around each blob, that agreed with the choice of the hybrid, and by definition is between $\frac{1}{3}$ and $1$. See Fig. 5a-i for the full network with labelled cycles.

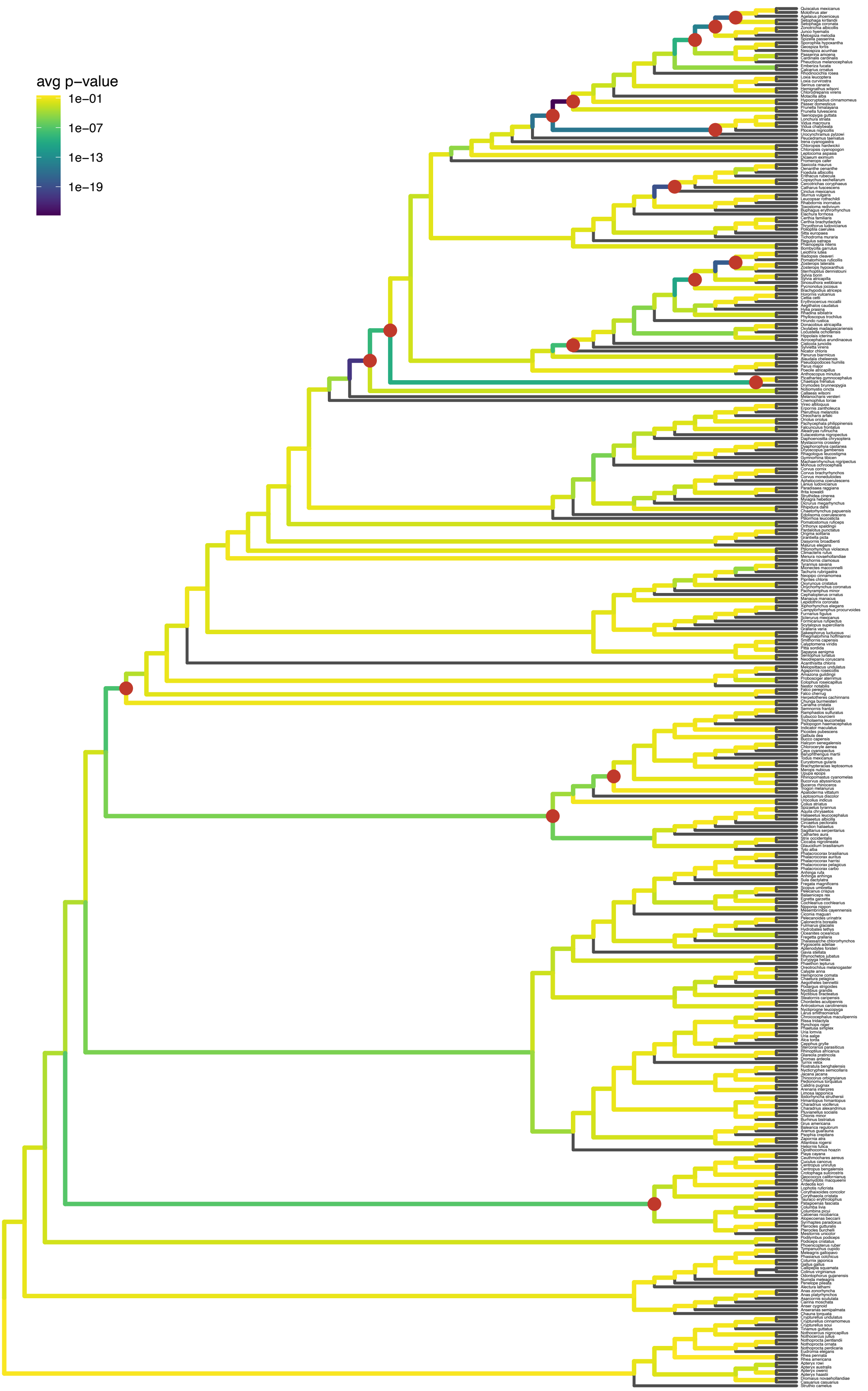


**Figure S14** Avian dataset. ASTRAL species tree from Stiller et al. (2024), which was used as input tree to TOB-QMC in our analysis. Internal branches are colored by the average p-value of the quartet tree test reported by TOB-QMC over 11 gene tree splits. Contracted branches in the final tree of blobs are marked with red at the tip of the branch.

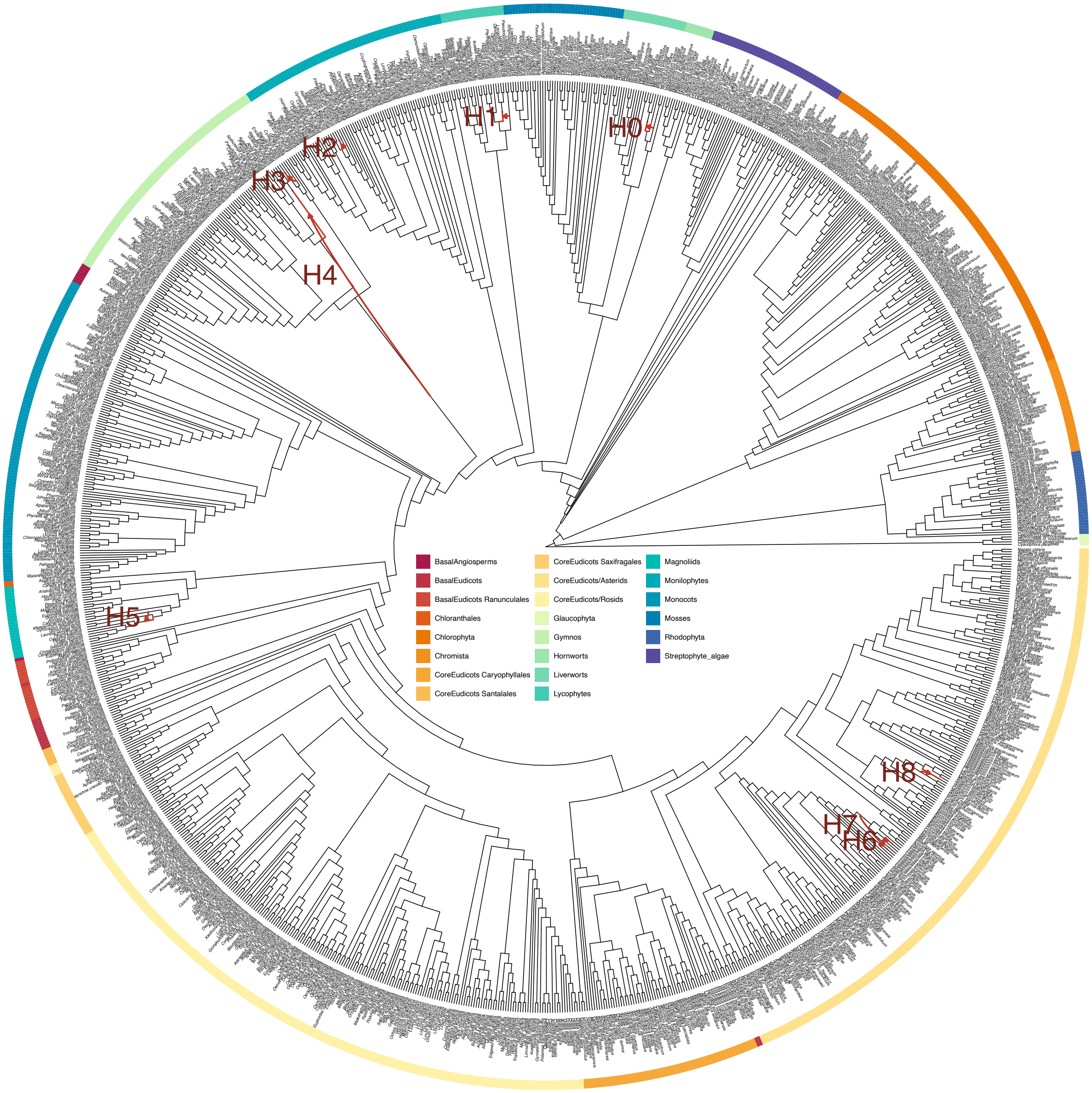


**Figure S15** Plants dataset. Entire phylogenetic network inferred by TOB-QMC+BROOQS. The 9 cycles are annotated by H0 to H8. Except for H4, which is a 5-blob, all cycles are of size 4.

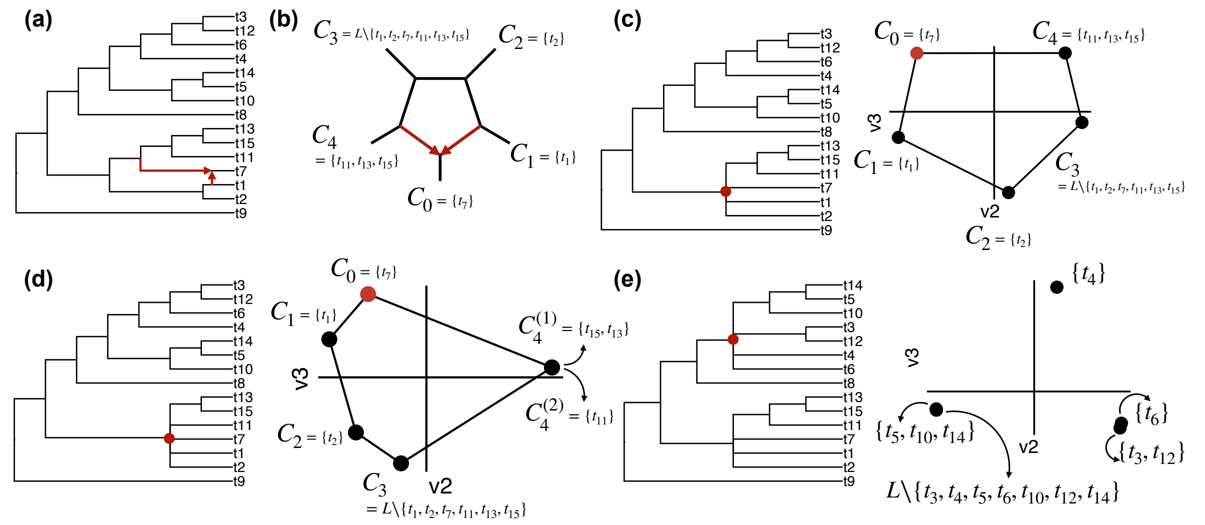

**Figure S16** An example of a network on the leaf set $L = \{t_1, t_2, \ldots, t_{15}\}$ with a 5-cycle (Fixed-ILS dataset). **(a)** True network with the hybrid edges marked with red. **(b)** The 5-cycle $\Omega$ from the true network in **(a)**. Components $\mathcal{C}$ are labeled from $C_0$ to $C_4$, with $C_0$ as the hybrid component. The true cyclic ordering is $\pi^*(\mathcal{C}) = [C_0, C_1, \ldots, C_4]$. **(c)** The true tree of blobs from the network in **(a)**. The polytomy corresponding to the cycle shown in **(b)** is marked in red. This polytomy induces the same set of components. Embedding the F-matrix $F$ computed from the true tree of blobs and true gene trees, results in a closed-path on the components of $\Omega$ with the correct cyclic ordering. **(d)** Contracting an extra branch around the cycle $\Omega$ results in a 6-blob where four components $C_0, C_1, C_2, C_3$ are unchanged, and one component is partitioned into two new components $C_4^{(1)} \cup C_4^{(2)} = C_4$. Running BROOQS on this tree of blobs and true gene trees results in the embedding presented on the right. The two partitions of $C_4$, $C_4^{(1)}$ and $C_4^{(2)}$, are fully overlapping in the embedding. The correct cyclic ordering between the rest of the components is preserved; however, the eigenvectors presenting these components (i.e., embeddings shown) are distorted due to division of one component into two. **(e)** Randomly contracting two adjacent tree-edges in the network in **(a)**, results in an additional 5-blob in the tree of blobs (marked in red). This blob does not correspond to any cycles in the true network, and therefore, no cyclic ordering exists on the components induced by this blob. Running BROOQS on this tree of blobs and true gene trees results in a triangle-like embedding dividing the components into three groups, with overlapping embeddings for the components in each group.

### B Supplementary Claims and Proofs

Throughout the proofs, for ease of notation, we use the conventions below.

**Simplified notation:**

- Recall that $\pi^*(\mathcal{C}) = [C_{\pi^*(0)}, C_{\pi^*(1)}, \ldots, C_{\pi^*(d-1)}]$ is one of the equivalent true orderings where $C_{\pi^*(0)}$ is the hybrid component. We will assume (w.o.l.g) that $\pi^*(i) = i$ everywhere, as shown in Figure S17. This will free us from having to write $\pi^*(i)$ in all expressions, for example allowing us to write $CF(C_iC_j \mid C_kC_l)$ instead of $CF(C_{\pi^*(i)}C_{\pi^*(j)} \mid C_{\pi^*(k)}C_{\pi^*(l)})$ and $F[i,j]$ instead of $F[\pi^*(i), \pi^*(j)]$. Since all proofs are only concerned with the true order, this adds no ambiguity.
- To further simplify notation, we use the shorthand $CF_{i,j|k,l}$ interchangeably for $CF(C_iC_j \mid C_kC_l)$.

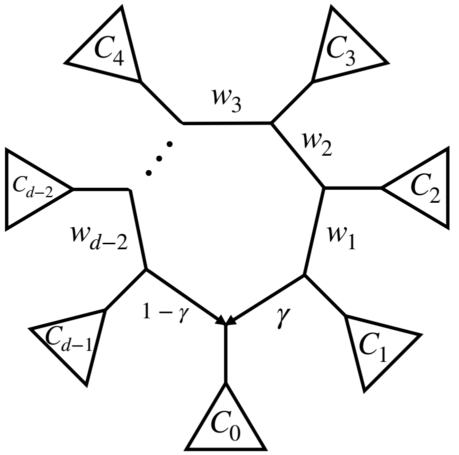


**Figure S17** A metric cycle $\Omega$ with components $\mathcal{C}$. To simplify notation, we assume the true order is $\pi^*(\mathcal{C}) = [C_0, C_1, \ldots, C_{d-1}]$, where $C_0$ is the hybrid component, and $\gamma$ and $1-\gamma$ are the inheritance probabilities of each hybrid edge. $w_1, w_2, \ldots, w_{d-2}$ are the CU lengths of the branches around $\Omega$ as shown in the figure.

#### B.1 Proof of Theorem 1

**Theorem 1:** Let $\Omega$ be a $d$-cycle where $4 \le d \le 15$ with components $\mathcal{C}$ and true cyclic ordering $\pi^*(\mathcal{C})$. Then, assuming NMSC, the permuted F-matrix $\pi^*(F)$ is a strict circular Robinson similarity matrix.

The proof relies on a series of lemmas and remarks provided below. We first state the overall proof of Theorem 1, followed by the formal statement of all the intermediate results and their proofs.

*Proof of Theorem 1.* Recall $\gamma \in (0,1)$ and $1-\gamma$ denote the inheritance probabilities of the two hybrid edges entering the hybrid node (Figure S17), and let $w_1, w_2, \ldots, w_{d-2}$ denote the edge lengths of the branches around $\Omega$ (see Figure S17), where $w_i > 0$ for all $0 < i < d-1$. Theorem 1 states that for every component $C_i$ $(0 \le i < d)$, the corresponding row of the F-matrix is strictly circularly unimodal. Recalling our simplified notation ($\pi^*(\mathcal{C}) = [C_0, C_1, \ldots, C_{d-1}]$ and $\pi^*(F) = F$), this is equivalent to saying that the sequence $[F[i, (i+1) \bmod d],\ F[i, (i+2) \bmod d],\ \ldots,\ F[i, (i+d-1) \bmod d]]$ is strictly decreasing up to a minimum and strictly increasing thereafter (i.e., is *strictly unimodal*).

We prove this property separately for the hybrid and non-hybrid components. The proof for non-hybrid components holds for every $d \ge 4$, whereas the proof for the hybrid component requires $d < 16$. Consequently, Theorem 1 is stated only for cycles with $4 \le d < 16$.

**Case 1 – Non-hybrids:** In Section B.1.1, we will show in Lemma 2 that for two non-hybrid components $C_i$ and $C_j$ for $0 < i < j < d-1$,

$$F[i,j] > F[i,j+1]\ .$$

Then we show in Remark 3 (building on Lemmas 3 and 4) that for every component $1 < i < d-1$,

$$F[i,0] < \max(F[i,1], F[i,d-1])\,. \tag{S3}$$

Combining these two easily proves the case for non-hybrids. From Lemma 2, we have:

$$F[i,i+1] > F[i,i+2] > \cdots > F[i,d-1]$$

and

$$F[i,1] < F[i,2] < \cdots < F[i,i-1].$$

From these and Equation (S3), it follows that the sequence $[F[i,i+1], F[i,i+2], \cdots, F[i,d-1], F[i,0], F[i,1], \cdots, F[i,i-1]]$ is strictly unimodal with at most two consecutive minimums. For each $C_i$, the minimum F-score corresponds to either $F[i,0]$, $F[i,1]$, or $F[i,d-1]$. Note that by Equation (S3), the three values cannot be equal, and therefore, the above sequence has at most consecutive minimums.

**Case 2 – hybrid:** For the hybrid component, Theorem 1 states that the sequence

$$[F[0,1], F[0,2], \ldots, F[0,d-1]]$$

is strictly unimodal. Let

$$\Delta_i := F[0,i+1] - F[0,i], \qquad 1 \le i \le d-2.$$

It is sufficient to show that the signs of $\Delta_1, \ldots, \Delta_{d-2}$ change from negative to positive exactly once.

**Definition 4** We define the following nonnegative quantities:

$$A_i = \sum_{k=i+2}^{d-1} e^{-(w_{i+1}+\cdots+w_{k-1})}, \quad B_i = \sum_{l=1}^{i-1} (l-1)\, e^{-(w_l+\cdots+w_{i-1})}, \quad C_i = \sum_{k=i+2}^{d-1} (d-1-k)\, e^{-(w_{i+1}+\cdots+w_{k-1})}, \quad D_i = \sum_{l=1}^{i-1} e^{-(w_l+\cdots+w_{i-1})},$$

together with

$$P_i = \tfrac{1}{3}(i-1)A_i + \tfrac{2}{3}B_i + (i-1), \qquad Q_i = \tfrac{2}{3}C_i + \tfrac{1}{3}(d-2-i)D_i + (d-2-i). \tag{S4}$$

It is easy to see that $P_1 = 0$ and $P_i > 0$ for $1 < i \le d-2$, and $Q_{d-2} = 0$ and $Q_i > 0$ for $1 \le i < d-2$.

Next, in Lemma 5, we show that

$$\Delta_i = \left(1 - e^{-w_i}\right) \Big[(1-\gamma)P_i - \gamma\, Q_i\Big].$$

Since $w_i > 0$ and $1 - e^{-w_i} > 0$, the sign of $\Delta_i$ is determined by the sign of $(1-\gamma)P_i - \gamma Q_i$. In Remark 4, we take care of the two neighbors of the hybrid component, stating that

$$\Delta_1 = -\gamma(1 - e^{-w_1})Q_1 < 0,$$

and

$$\Delta_{d-2} = (1-\gamma)(1 - e^{-w_{d-2}})P_{d-2} > 0.$$

which is what we need for the unimodal order. For other nodes, we define

$$\theta := \frac{\gamma}{1-\gamma} \in (0,\infty), \qquad R_i := \frac{P_i}{Q_i}, \quad 1 \le i \le d-3.$$

Since $P_1 = 0$, we have $R_1 = 0$, and we set $R_{d-2} = +\infty$, which naturally follows from $P_{d-2} > 0$ and $Q_{d-2} = 0$. In Lemma 6, we prove that

$$R_1 < R_2 < \ldots < R_{d-2}$$

is a sufficient condition for the sequence

$$[F[0,1], F[0,2], \ldots, F[0,d-1]]$$

to be unimodal. Then, in Proposition 2, we use a computer program that casts $R_{i+1} - R_i$ as a polynomial in variables defined based on branch lengths $w$ and $\lambda$ to prove that as long as $d \leq 15$, the polynomial is strictly positive, showing that $R_1 < R_2 < \ldots < R_{d-2}$, which completes the proof. □

##### B.1.1 Auxiliary results for proof for non-hybrid components

---

**Lemma 2** [monotonicity.] Let $C_i$ and $C_j$ be two non-hybrid components in $\mathcal{C}$ such that $0 < i < j < d-1$. Then, for the non-hybrid component $C_{j+1}$, $F[i,j] > F[i,j+1]$.

*Proof* By definition,

$$F[i,j] = \sum_{\substack{k<l \\ k,l \neq i,j}} CF_{i,j|k,l}$$

To compare $F[i,j]$ and $F[i,j+1]$, we compare each summand in their definitions. Specifically, for every choice of two additional components $C_k$ and $C_l$ distinct from $C_i$, $C_j$, and $C_{j+1}$, we compare the corresponding quartet CFs $CF_{i,j|k,l}$ and $CF_{i,j+1|k,l}$. We show that $CF_{i,j|k,l} \geq CF_{i,j+1|k,l}$ for every possible arrangement of the four components, with the inequality being strict for at least one arrangement. Summing over all choices of $C_k$ and $C_l$ then yields $F[i,j] > F[i,j+1]$.

The possible arrangements are shown in Figure S18 together with the corresponding induced metric quartet/quarnet.

- **Case 1.** Suppose $i < j < j+1 < k < l < d$. Then

$$CF_{i,j|k,l} = 1 - \frac{2}{3} e^{-(w_j + w_{j+1} + \cdots + w_{k-1})},$$

and

$$CF_{i,j+1|k,l} = 1 - \frac{2}{3} e^{-(w_{j+1} + \cdots + w_{k-1})}.$$

Since $w_j + w_{j+1} + \cdots + w_{k-1} > w_{j+1} + \cdots + w_{k-1}$, we obtain $CF_{i,j|k,l} > CF_{i,j+1|k,l}$.

- **Case 2.** Suppose $0 < k < l < i < j$. Then

$$CF_{i,j|k,l} = 1 - \frac{2}{3} e^{-(w_l + \cdots + w_{i-1})},$$

and

$$CF_{i,j+1|k,l} = 1 - \frac{2}{3} e^{-(w_l + \cdots + w_{i-1})}.$$

Therefore, $CF_{i,j|k,l} = CF_{i,j+1|k,l}$.

- **Case 3.** Suppose $0 < k < i$ and $j+1 < l < d$. Then

$$CF_{i,j|k,l} = \frac{1}{3} e^{-(w_i + \cdots + w_{j-1})},$$

and

$$CF_{i,j+1|k,l} = \frac{1}{3} e^{-(w_i + \cdots + w_{j-1} + w_j)}.$$

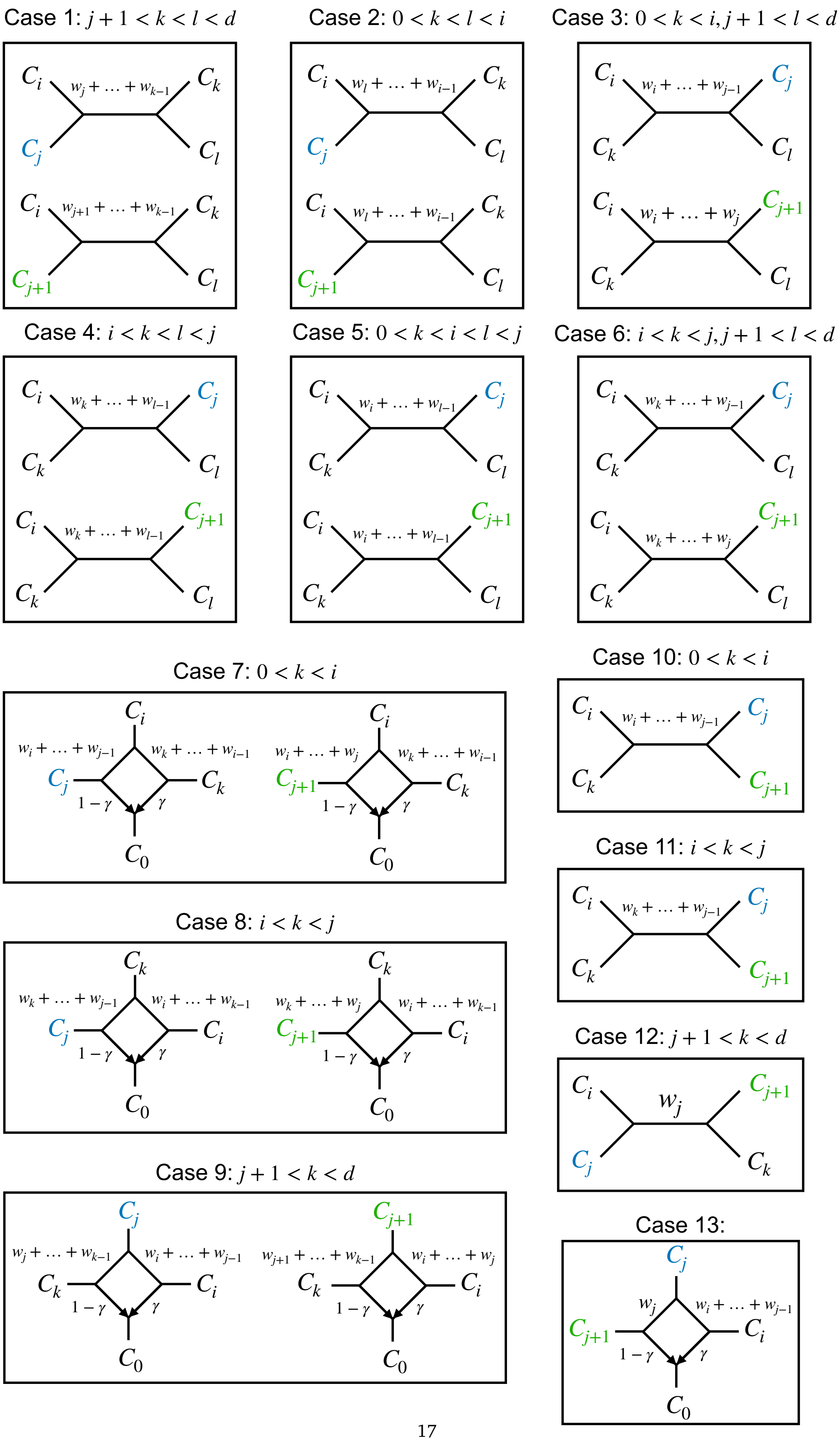




**Figure S18** Every possible choice of four components including $C_i$ and $C_j/C_{j+1}$. Corresponding metric quartet/quarnet is shown for each case.

Since $w_i + \cdots + w_{j-1} < w_i + \cdots + w_{j-1} + w_j$, we obtain $CF_{i,j|k,l} > CF_{i,j+1|k,l}$.

- **Case 4.** Suppose $i < k < l < j$. Then

$$CF_{i,j|k,l} = \frac{1}{3}e^{-(w_k+\cdots+w_{l-1})},$$

and

$$CF_{i,j+1|k,l} = \frac{1}{3}e^{-(w_k+\cdots+w_{l-1})}.$$

Therefore, $CF_{i,j|k,l} = CF_{i,j+1|k,l}$.

- **Case 5.** Suppose $0 < k < i < l < j$. Then

$$CF_{i,j|k,l} = \frac{1}{3}e^{-(w_i+\cdots+w_{l-1})},$$

and

$$CF_{i,j+1|k,l} = \frac{1}{3}e^{-(w_i+\cdots+w_{l-1})}.$$

Therefore, $CF_{i,j|k,l} = CF_{i,j+1|k,l}$.

- **Case 6.** Suppose $i < k < j$ and $j+1 < l < d$. Then

$$CF_{i,j|k,l} = \frac{1}{3}e^{-(w_k+\cdots+w_{j-1})},$$

and

$$CF_{i,j+1|k,l} = \frac{1}{3}e^{-(w_k+\cdots+w_{j-1}+w_j)}.$$

Since $w_k + \cdots + w_{j-1} < w_k + \cdots + w_{j-1} + w_j$, we obtain $CF_{i,j|k,l} > CF_{i,j+1|k,l}$.

- **Case 7.** Suppose $l = 0 < k < i < j$. Then

$$CF_{i,j|k,l} = CF_{i,j|k,0} = \gamma[1 - \frac{2}{3}e^{-(w_k+\cdots+w_{i-1})}] + (1-\gamma)[\frac{1}{3}e^{-(w_i+\cdots+w_{j-1})}],$$

and

$$CF_{i,j+1|k,l} = CF_{i,j+1|k,0} = \gamma[1 - \frac{2}{3}e^{-(w_k+\cdots+w_{i-1})}] + (1-\gamma)[\frac{1}{3}e^{-(w_i+\cdots+w_{j-1}+w_j)}],$$

Since $w_i + \cdots + w_{j-1} < w_i + \cdots + w_{j-1} + w_j$, we obtain $CF_{i,j|k,0} > CF_{i,j+1|k,0}$.

- **Case 8.** Suppose $l = 0 < i < k < j$. Then

$$CF_{i,j|k,0} = \gamma[\frac{1}{3}e^{-(w_i+\cdots+w_{k-1})}] + (1-\gamma)[\frac{1}{3}e^{-(w_k+\cdots+w_{j-1})}],$$

and

$$CF_{i,j+1|k,0} = \gamma[\frac{1}{3}e^{-(w_i+\cdots+w_{k-1})}] + (1-\gamma)[\frac{1}{3}e^{-(w_k+\cdots+w_{j-1}+w_j)}].$$

Since $w_k + \cdots + w_{j-1} < w_k + \cdots + w_{j-1} + w_j$, we obtain $CF_{i,j|k,0} > CF_{i,j+1|k,0}$.

- **Case 9.** Suppose $l = 0 < i < j < j+1 < k < d$. Then

$$CF_{i,j|k,0} = \gamma[\frac{1}{3}e^{-(w_i+\cdots+w_{j-1})}] + (1-\gamma)[1 - \frac{2}{3}e^{-(w_j+w_{j+1}+\cdots+w_{k-1})}],$$

and

$$CF_{i,j+1|k,0} = \gamma[\frac{1}{3}e^{-(w_i+\cdots+w_{j-1}+w_j)}] + (1-\gamma)[1-\frac{2}{3}e^{-(w_{j+1}+\cdots+w_{k-1})}].$$

Since $w_i + \cdots + w_{j-1} < w_i + \cdots + w_{j-1} + w_j$ and $w_j + w_{j+1} + \cdots + w_{k-1} > w_{j+1} + \cdots + w_{k-1}$, we obtain $CF_{i,j|k,0} > CF_{i,j+1|k,0}$.

- **Case 10.** Suppose $0 < k < i$ and $l = j+1$ for $CF_{i,j|k,l}$ and $j$ for $CF_{i,j+1|k,l}$ . Then

$$CF_{i,j|k,j+1} = \frac{1}{3}e^{-(w_i+\cdots+w_{j-1})},$$

and

$$CF_{i,j+1|k,j} = \frac{1}{3}e^{-(w_i+\cdots+w_{j-1})}.$$

Therefore, $CF_{i,j|k,j+1} = CF_{i,j+1|k,j}$.

- **Case 11.** Suppose $i < k < j$ and $l = j+1$ for $CF_{i,j|k,l}$ and $l = j$ for $CF_{i,j+1|k,l}$. Then

$$CF_{i,j|k,j+1} = \frac{1}{3}e^{-(w_k+\cdots+w_{j-1})},$$

and

$$CF_{i,j+1|k,j} = \frac{1}{3}e^{-(w_k+\cdots+w_{j-1})}.$$

Therefore, $CF_{i,j|k,j+1} = CF_{i,j+1|k,j}$.

- **Case 12.** Suppose $j+1 < k < d$ and $l = j+1$ for $CF_{i,j|k,l}$ and $l = j$ for $CF_{i,j+1|k,l}$.

$$CF_{i,j|k,j+1} = 1 - \frac{2}{3}e^{-w_j},$$

and

$$CF_{i,j+1|k,j} = \frac{1}{3}e^{-w_j}.$$

Therefore, $CF_{i,j|k,j+1} > CF_{i,j+1|k,j}$.

- **Case 13.** Suppose $k = 0$ and $l = j+1$ for $CF_{i,j|k,l}$ and $l = j$ for $CF_{i,j+1|k,l}$. Then

$$CF_{i,j|j+1,0} = \gamma[\frac{1}{3}e^{-(w_i+\cdots+w_{j-1})}] + (1-\gamma)[1-\frac{2}{3}e^{-w_j}],$$

and

$$CF_{i,j+1|j,0} = \gamma[\frac{1}{3}e^{-(w_i+\cdots+w_{j-1})}] + (1-\gamma)[\frac{1}{3}e^{-w_j}].$$

Therefore, $CF_{i,j|j+1,0} > CF_{i,j+1|j,0}$.

The cases above exhaust all possible relative positions of $C_i$, $C_j$, $C_k$, and $C_l$ on the cycle. Since $d \geq 4$ and $C_i$, $C_j$, and $C_{j+1}$ are all non-hybrid components ($0 < i < j < d-1$), the hybrid component $C_0$ is distinct from all three components. Consequently, although not every case occurs for every cycle and every choice of $C_i$ and $C_j$, Case 13 is always included among the summands of $F[i,j]$. In this case, $CF_{i,j|k,0} > CF_{i,j+1|k,0}$. Since all remaining cases satisfy $CF_{i,j|k,l} \geq CF_{i,j+1|k,l}$, it follows that $F[i,j] > F[i,j+1]$, for every cycle $\Omega$ and every choice of $C_i$ and $C_j$. □

**Lemma 3** (Hybrid CF) Let $C_0$ be the hybrid component of $\Omega$, and let $\gamma$ and $1-\gamma$ denote the inheritance probabilities of the two hybrid edges adjacent to the components $C_1$ and $C_{d-1}$, respectively (Figure S17). For any distinct components $C_i, C_k, C_l \notin \{C_0, C_1, C_{d-1}\}$,

$$CF_{0,i|k,l} = \gamma\, CF_{1,i|k,l} + (1-\gamma)\, CF_{d-1,i|k,l}.$$

*Proof* Let $\{C_0C_i \mid C_kC_l\}$ denote the event that a gene tree induced on four taxa $t_0 \in C_0$, $t_i \in C_i$, $t_k \in C_k$, and $t_l \in C_l$ has topology $t_0t_i \mid t_kt_l$. Then, $CF_{0,i|k,l}$ is the probability of this event. Under the NMSC model, the lineage sampled from the hybrid component $C_0$ independently chooses the parental edge adjacent to $C_1$ and $C_{d-1}$ with probabilities $\gamma$ and $1-\gamma$. Let $H_1$ and $H_2$ denote these two events, respectively. Conditioning on this choice and applying the law of total probability gives

$$CF_{0,i|k,l} = \Pr(\{C_0C_i \mid C_kC_l\}) = \Pr(\{C_0C_i \mid C_kC_l\} \mid H_1)\Pr(H_1) + \Pr(\{C_0C_i \mid C_kC_l\} \mid H_2)\Pr(H_2),$$

Conditioned on $H_1$, the induced metric quartet is identical to the one obtained by replacing $C_0$ with $C_1$. Likewise, conditioned on $H_2$, the induced metric quartet is identical to the one obtained by replacing $C_0$ with $C_{d-1}$. Since quartet concordance factors depend only on the induced metric quartet Baños (2019),

$$\Pr(\{C_0C_i \mid C_kC_l\} \mid H_1) = CF_{1,i|k,l},$$

and

$$\Pr(\{C_0C_i \mid C_kC_l\} \mid H_2) = CF_{d-1,i|k,l}.$$

Substituting these probabilities together with $\Pr(H_1) = \gamma$ and $\Pr(H_2) = 1-\gamma$ completes the proof. □

---

**Lemma 4** (non-neighbors) For any component $C_i$, where $1 < i < d-1$, we have

$$F[i,0] < \gamma F[i,1] + (1-\gamma)F[i,d-1].$$

*Proof* We partition the summands of $F[i,0]$, $F[i,1]$, and $F[i,d-1]$ into the following three classes. First,

$$F[i,0] = \sum_{\substack{k<l \\ k,l\notin\{i,0,1,d-1\}}} CF_{i,0|k,l} + \sum_{k\notin\{i,0,1,d-1\}} \left(CF_{i,0|k,1} + CF_{i,0|k,d-1}\right) + CF_{i,0|1,d-1},$$

$$F[i,1] = \sum_{\substack{k<l \\ k,l\notin\{i,0,1,d-1\}}} CF_{i,1|k,l} + \sum_{k\notin\{i,0,1,d-1\}} \left(CF_{i,1|k,0} + CF_{i,1|k,d-1}\right) + CF_{i,1|0,d-1},$$

$$F[i,d-1] = \sum_{\substack{k<l \\ k,l\notin\{i,0,1,d-1\}}} CF_{i,d-1|k,l} + \sum_{k\notin\{i,0,1,d-1\}} \left(CF_{i,d-1|k,1} + CF_{i,d-1|k,0}\right) + CF_{i,d-1|1,0}.$$

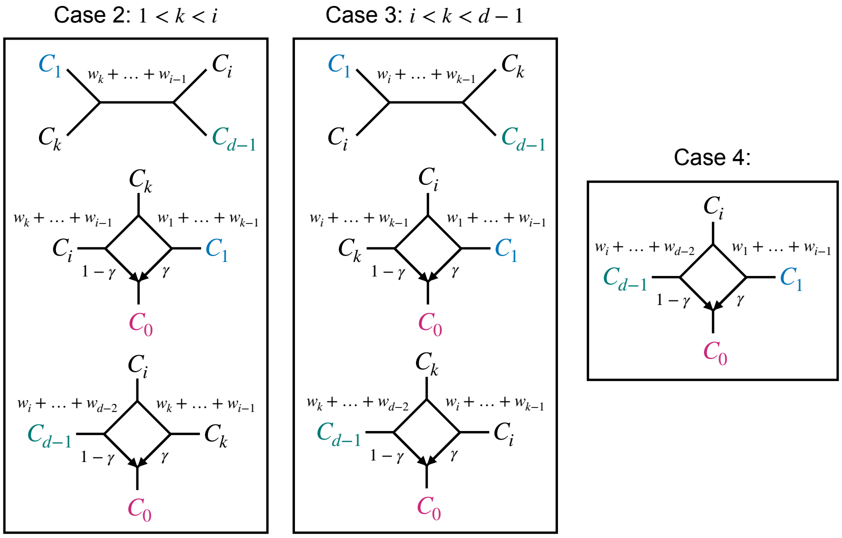


**Figure S19** Every possible choice of four components including $C_i$ and two of $C_0$, $C_1$, $C_{d-1}$. Corresponding metric quartet/quarnet is shown for each case.

The stated lemma is equivalent to showing that

$$
\begin{aligned}
0 <& \gamma F[i,1] + (1-\gamma)F[i,d-1] - F[i,0] = \\
&\overbrace{\sum_{\substack{k<l \\ k,l \notin \{i,0,1,d-1\}}} \gamma CF_{i,1|k,l} + (1-\gamma)CF_{i,d-1|k,l} - CF_{i,0|k,l}}^{\text{case 1}} + \\
&\overbrace{\sum_{k \neq i,0,1,d-1} \gamma\left(CF_{i,1|k,0} + CF_{i,1|k,d-1}\right) + (1-\gamma)\left(CF_{i,d-1|k,1} + CF_{i,d-1|k,0}\right) - \left(CF_{i,0|k,1} + CF_{i,0|k,d-1}\right)}^{\text{case 2 } (k<i) \text{ and case 3} (k>i)} + \\
&\overbrace{\gamma CF_{i,1|0,d-1} + (1-\gamma)CF_{i,d-1|1,0} - CF_{i,0|1,d-1}}^{\text{case 4}}.
\end{aligned}
$$

For each case, we show that its contribution is either positive or zero. Figure S19 shows all non-trivial cases, each of which we next examine.

- **Case 1.** Let $k < l$, where $k, l \neq i, 0, 1, d-1$. By Lemma 3,

$$CF_{0,i|k,l} = \gamma CF_{1,i|k,l} + (1-\gamma)CF_{d-1,i|k,l}.$$

  Therefore, the contribution of this case to $\gamma F[i,1] + (1-\gamma)F[i,d-1] - F[i,0]$ is 0.
- **Case 2.** Let $1 < k < i$. From the metric quartets we have

$$
\begin{aligned}
CF_{1,i|k,0} + CF_{1,i|k,d-1} &= \gamma[\frac{1}{3}e^{-(w_1+\cdots+w_{k-1})}] + (1-\gamma)[\frac{1}{3}e^{-(w_k+\cdots+w_{i-1})}] + \frac{1}{3}e^{-(w_k+\cdots+w_{i-1})} \\
CF_{d-1,i|k,1} + CF_{d-1,i|k,0} &= 1 - \frac{2}{3}e^{-(w_k+\cdots+w_{i-1})} + \gamma[1 - \frac{2}{3}e^{-(w_k+\cdots+w_{i-1})}] + (1-\gamma)[\frac{1}{3}e^{-(w_i+\cdots+w_{d-2})}] \\
CF_{i,0|k,1} + CF_{i,0|k,d-1} &= \gamma[\frac{1}{3}e^{-(w_1+\cdots+w_{k-1})}] + (1-\gamma)[1 - \frac{2}{3}e^{-(w_k+\cdots+w_{i-1})}] + \gamma[\frac{1}{3}e^{-(w_k+\cdots+w_{i-1})}] + (1-\gamma)[\frac{1}{3}e^{-(w_i+\cdots+w_{d-2})}].
\end{aligned}
$$

We can compute the contribution of $k$ to $\gamma F[i,1]+(1-\gamma)F[i,d-1]-F[i,0]$ denoted by $\Delta$ as:

$$
\begin{aligned}
\Delta = \gamma\Big(&CF_{1,i|k,0} + CF_{1,i|k,d-1}\Big) \\
&+ (1-\gamma)\Big(CF_{d-1,i|k,1} + CF_{d-1,i|k,0}\Big) \\
&- \Big(CF_{0,i|k,1} + CF_{0,i|k,d-1}\Big) \\
= &\gamma^2\left[\frac{1}{3}e^{-(w_1+\cdots+w_{k-1})}\right] + \gamma(1-\gamma)\left[\frac{1}{3}e^{-(w_k+\cdots+w_{i-1})}\right] + \gamma[\frac{1}{3}e^{-(w_k+\cdots+w_{i-1})}] \\
&+ \gamma(1-\gamma)\left[1-\frac{2}{3}e^{-(w_k+\cdots+w_{i-1})}\right] + (1-\gamma)^2\left[\frac{1}{3}e^{-(w_i+\cdots+w_{d-2})}\right] + (1-\gamma)[\frac{1}{3}e^{-(w_i+\cdots+w_{d-2})}] \\
&- \gamma\left[\frac{1}{3}e^{-(w_1+\cdots+w_{k-1})}\right] - (1-\gamma)\left[\frac{1}{3}e^{-(w_i+\cdots+w_{d-2})+}\right] - \gamma[\frac{1}{3}e^{-(w_k+\cdots+w_{i-1})}] - (1-\gamma)[\frac{1}{3}e^{-(w_i+\cdots+w_{d-2})}] \\
= &\gamma(1-\gamma)\left[1-\frac{1}{3}e^{-(w_k+\cdots+w_{i-1})}\right] - \gamma(1-\gamma)\left[\frac{1}{3}e^{-(w_1+\cdots+w_{k-1})}\right] \\
&- \gamma(1-\gamma)\left[\frac{1}{3}e^{-(w_i+\cdots+w_{d-2})}\right] \\
= &\gamma(1-\gamma)\left[1-\frac{1}{3}e^{-(w_k+\cdots+w_{i-1})} - \frac{1}{3}e^{-(w_1+\cdots+w_{k-1})} - \frac{1}{3}e^{-(w_i+\cdots+w_{d-2})}\right].
\end{aligned}
$$

Since $\gamma(1-\gamma)>0$ and $\frac{1}{3}e^{-x}<\frac{1}{3}$ for every $x>0$, this contribution is strictly positive ($\Delta>0$).

- **Case 3.** Let $i<k<d-1$. The proof of this case is identical to Case 2, with variables $i$ and $k$ swapped (see Figure S19), but leading to the same strictly positive term at the end.
- **Case 4.** This is the unique quarnet on $C_0, C_1, C_i, C_{d-1}$. For this quarnet, we have:

$$
\begin{aligned}
CF_{0,i|1,d-1} &= \gamma[\frac{1}{3}e^{-(w_1+\cdots+w_{i-1})}] + (1-\gamma)[\frac{1}{3}e^{-(w_i+\cdots+w_{d-2})}] \\
CF_{1,i|0,d-1} &= \gamma[1-\frac{2}{3}e^{-(w_1+\cdots+w_{i-1})}] + (1-\gamma)[\frac{1}{3}e^{-(w_i+\cdots+w_{d-2})}] \\
CF_{d-1,i|0,1} &= \gamma[\frac{1}{3}e^{-(w_1+\cdots+w_{i-1})}] + (1-\gamma)[1-\frac{2}{3}e^{-(w_i+\cdots+w_{d-2})}].
\end{aligned}
$$

By the CFs of a network quarnet (Fig. S1),

$$CF_{0,i|1,d-1} < CF_{1,i|0,d-1}$$

and

$$CF_{0,i|1,d-1} < CF_{d-1,i|0,1},$$

it follows immediately that

$$CF_{0,i|1,d-1} < \gamma CF_{1,i|0,d-1} + (1-\gamma)CF_{d-1,i|0,1}.$$

The four cases above exhaust all summands in the three F-scores. Case 1 contributes 0, whereas Cases 2–4 contribute a strictly positive amount. Since Case 4 is always present, the total contribution is strictly positive. Therefore,

$$\gamma F[i,1]+(1-\gamma)F[i,d-1]-F[i,0]>0,$$

which completes the proof. □

---

**Remark 3** Since

$$\gamma F[i,1] + (1-\gamma)F[i,d-1] \le \max(F[i,1], F[i,d-1])\,,$$

Lemma 4 immediately implies that

$$F[i,0] < \max(F[i,1], F[i,d-1])\,.$$

##### B.1.2 Auxiliary Results for the Proof for the hybrid component

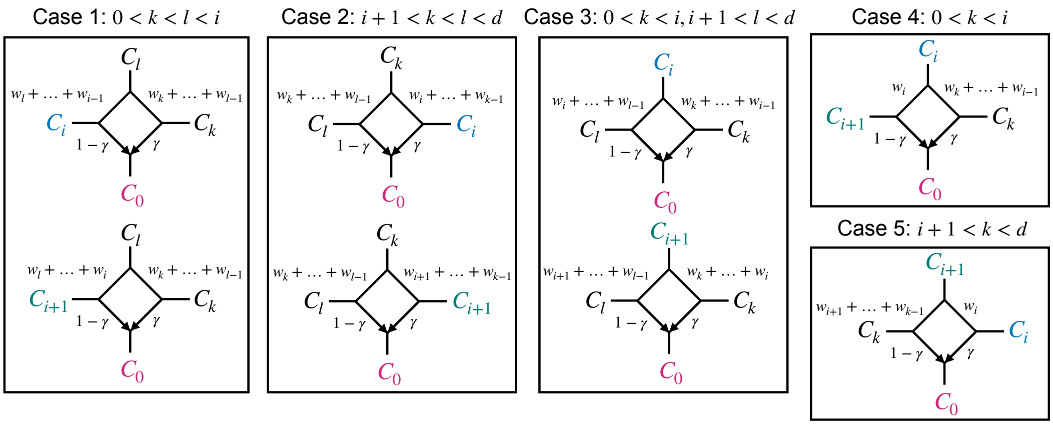


**Figure S20** Every possible choice of four components including $C_0$ and $C_i/C_{i+1}$. Corresponding metric quarnets are shown for each case.

---

**Lemma 5** ($\Delta$ for hybrids) For every $1 \le i \le d-2$,

$$\Delta_i = F[0,i+1] - F[0,i] \;=\; \left(1 - e^{-w_i}\right)\Big[(1-\gamma)P_i - \gamma Q_i\Big]. \tag{S5}$$

*Proof* We have

$$F[0,i] = \sum_{k<l;k,l\notin\{0,i,i+1\}} CF_{0,i|k,l} + \sum_{k\notin\{0,i,i+1\}} CF_{0,i|i+1,k}.$$

Similarly,

$$F[0,i+1] = \sum_{k<l;k,l\notin\{0,i,i+1\}} CF_{0,i+1|k,l} + \sum_{k\notin\{0,i,i+1\}} CF_{0,i+1|i,k}.$$

For each choice of four components, we compute its contribution to $\Delta_i$. Figure S20 shows all possible cases.

- **Case 1.** Suppose $0 < k < l < i$. For any choice of $k, l$ we have

$$CF_{0,i|k,l} = \gamma[\frac{1}{3}e^{-(w_k+\cdots+w_{l-1})}] + (1-\gamma)[1 - \frac{2}{3}e^{-(w_l+\cdots+w_{i-1})}]$$

$$CF_{0,i+1|k,l} = \gamma[\frac{1}{3}e^{-(w_k+\cdots+w_{l-1})}] + (1-\gamma)[1 - \frac{2}{3}e^{-(w_l+\cdots+w_i)}].$$

Therefore,

$$CF_{0,i+1|k,l} - CF_{0,i|k,l} = (1-\gamma)(1-e^{-w_i})[\frac{2}{3}e^{-(w_l+\cdots+w_{i-1})}].$$

Summing over all choices of $0 < k < l < i$, we get

$$\Delta_i = \sum_{l=1}^{i-1}\sum_{k=1}^{l-1}(1-\gamma)(1-e^{-w_i})[\frac{2}{3}e^{-(w_l+\cdots+w_{i-1})}] = \frac{2}{3}(1-\gamma)(1-e^{-w_i})\sum_{l=1}^{i-1}(l-1)e^{-(w_l+\cdots+w_{i-1})} = \frac{2}{3}(1-\gamma)(1-e^{-w_i})B_i.$$

- **Case 2.** Suppose $i+1 < k < l < d$. For any choice of $k, l$ we have

$$CF_{0,i|k,l} = \gamma[1-\frac{2}{3}e^{-(w_i+\cdots+w_{k-1})}] + (1-\gamma)[\frac{1}{3}e^{-(w_k+\cdots+w_{l-1})}]$$
$$CF_{0,i+1|k,l} = \gamma[1-\frac{2}{3}e^{-(w_{i+1}+\cdots+w_{k-1})}] + (1-\gamma)[\frac{1}{3}e^{-(w_k+\cdots+w_{l-1})}]$$

Therefore,

$$CF_{0,i+1|k,l} - CF_{0,i|k,l} = -\gamma(1-e^{-w_i})[\frac{2}{3}e^{-(w_{i+1}+\cdots+w_{k-1})}].$$

Summing over all choices of $i+1 < k < l < d$, we get

$$\Delta_i = \sum_{k=i+2}^{d-1}\sum_{l=k+1}^{d-1} -\gamma(1-e^{-w_i})[\frac{2}{3}e^{-(w_{i+1}+\cdots+w_{k-1})}] = -\frac{2}{3}\gamma(1-e^{-w_i})\sum_{k=i+2}^{d-1}(d-1-k)e^{-(w_{i+1}+\cdots+w_{k-1})} = -\frac{2}{3}\gamma(1-e^{-w_i})C_i.$$

- **Case 3.** Suppose $0 < k < i$ and $i+1 < l < d$. For any choice of $k, l$ we have

$$CF_{0,i|k,l} = \gamma[\frac{1}{3}e^{-(w_k+\cdots+w_{i-1})}] + (1-\gamma)[\frac{1}{3}e^{-(w_i+\cdots+w_{l-1})}]$$
$$CF_{0,i+1|k,l} = \gamma[\frac{1}{3}e^{-(w_k+\cdots+w_i)}] + (1-\gamma)[\frac{1}{3}e^{-(w_{i+1}+\cdots+w_{l-1})}].$$

Therefore,

$$CF_{0,i+1|k,l} - CF_{0,i|k,l} = -\gamma(1-e^{-w_i})[\frac{1}{3}e^{-(w_k+\cdots+w_{i-1})}] + (1-\gamma)(1-e^{-w_i})[\frac{1}{3}e^{-(w_{i+1}+\cdots+w_{l-1})}].$$

Summing over all choices of $0 < k < i$ and $i+1 < l < d$, we get

$$\begin{aligned}\Delta_i &= \sum_{1\le k<i}\sum_{i+1<l<d}\left(-\gamma(1-e^{-w_i})\frac{1}{3}e^{-(w_k+\cdots+w_{i-1})} + (1-\gamma)(1-e^{-w_i})\frac{1}{3}e^{-(w_{i+1}+\cdots+w_{l-1})}\right)\\ &= -\frac{1}{3}\gamma(1-e^{-w_i})(d-2-i)\sum_{k=1}^{i-1}e^{-(w_k+\cdots+w_{i-1})} + \frac{1}{3}(1-\gamma)(1-e^{-w_i})(i-1)\sum_{l=i+2}^{d-1}e^{-(w_{i+1}+\cdots+w_{l-1})}\\ &= -\frac{1}{3}\gamma(1-e^{-w_i})(d-2-i)D_i + \frac{1}{3}(1-\gamma)(1-e^{-w_i})(i-1)A_i.\end{aligned}$$

- **Case 4.** Suppose $0 < k < i$. For any choice of $k$, we have

$$CF_{0,i|i+1,k} = \gamma[\frac{1}{3}e^{-(w_k+\cdots+w_{i-1})}] + (1-\gamma)[\frac{1}{3}e^{-w_i}]$$
$$CF_{0,i+1|i,k} = \gamma[\frac{1}{3}e^{-(w_k+\cdots+w_{i-1})}] + (1-\gamma)[1-\frac{2}{3}e^{-w_i}].$$

Therefore,

$$CF_{0,i+1|i,k} - CF_{0,i|i+1,k} = (1-\gamma)(1-e^{-w_i}).$$

Summing over all choices of $0 < k < i$, we get

$$\Delta_i = \sum_{0<k<i} (1-\gamma)(1-e^{-w_i}) = (1-\gamma)(1-e^{-w_i})(i-1).$$

- **Case 5.** Suppose $i+1 < k < d$. For any choice of $k$, we have

$$\begin{aligned} CF_{0,i|i+1,k} &= \gamma[1 - \frac{2}{3}e^{-w_i}] + (1-\gamma)[\frac{1}{3}e^{-(w_{i+1}+\cdots+w_{k-1})}] \\ CF_{0,i+1|i,k} &= \gamma[\frac{1}{3}e^{-w_i}] + (1-\gamma)[\frac{1}{3}e^{-(w_{i+1}+\cdots+w_{k-1})}] \end{aligned}$$

Therefore,

$$CF_{0,i+1|i,k} - CF_{0,i|i+1,k} = -\gamma(1-e^{-w_i}).$$

Summing over all choices of $i+1 < k < d$, we get

$$\Delta_i = \sum_{i+1<k<d} -\gamma(1-e^{-w_i}) = -\gamma(1-e^{-w_i})(d-2-i).$$

Summing the contributions of all five cases gives

$$\begin{aligned} \Delta_i &= \frac{2}{3}(1-\gamma)(1-e^{-w_i})B_i - \frac{2}{3}\gamma(1-e^{-w_i})C_i \\ &\quad - \frac{1}{3}\gamma(1-e^{-w_i})(d-2-i)D_i + \frac{1}{3}(1-\gamma)(1-e^{-w_i})(i-1)A_i \\ &\quad + (1-\gamma)(1-e^{-w_i})(i-1) - \gamma(1-e^{-w_i})(d-2-i) \\ &= (1-e^{-w_i})\Big[(1-\gamma)P_i - \gamma Q_i\Big], \end{aligned}$$

which completes the proof of Lemma 5. □

---

**Remark 4** For every $1 \le i \le d-2$, we have

$$P_i \ge i-1 \qquad \text{and} \qquad Q_i \ge d-2-i.$$

In particular,

$$P_1 = 0, \qquad Q_{d-2} = 0,$$

whereas $P_i > 0$ for $i \ge 2$ and $Q_i > 0$ for $i \le d-3$. Consequently, plugging in Equation (S5),

$$\begin{aligned} \Delta_1 &= -\gamma(1-e^{-w_1})Q_1 < 0, \\ \Delta_{d-2} &= (1-\gamma)(1-e^{-w_{d-2}})P_{d-2} > 0. \end{aligned}$$

---

**Lemma 6** Let

$$\theta := \frac{\gamma}{1-\gamma} \in (0,\infty), \qquad R_i := \frac{P_i}{Q_i}, \quad 1 \le i \le d-3.$$

If $R_1 < R_2 < \cdots < R_{d-2}$, then the row of the F-matrix corresponding to $C_0$ is unimodal.

*Proof* Since $P_1 = 0$, we have $R_1 = 0$, and we set $R_{d-2} = +\infty$, which follows naturally from $P_{d-2} > 0$ and $Q_{d-2} = 0$. For $1 \le i \le d-3$, we have $Q_i > 0$, and therefore

$$S_i := Q_i\big((1-\gamma)R_i - \gamma\big) = (1-\gamma)Q_i\big(R_i - \theta\big).$$

Based on Lemma 5, $\Delta_i = \left(1 - e^{-w_i}\right)\Big[(1-\gamma)P_i - \gamma\, Q_i\Big]$ and thus, the sign of $\Delta_i$ is determined by the sign of $R_i - \theta$. For $i = d-2$, we have $\Delta_{d-2} > 0$ by Remark 4, which is consistent with $R_{d-2} = +\infty > \theta$.

Since

$$R_1 = 0 < \theta < R_{d-2},$$

and the sequence $R_1, \ldots, R_{d-2}$ is strictly increasing by assumption of the lemma, the set

$$\{\, i : R_i \ge \theta \,\}$$

is a non-empty terminal segment

$$\{m, m+1, \ldots, d-2\},$$

for some $2 \le m \le d-2$. Therefore,

$$\Delta_i < 0 \quad \text{for } i < m,$$

$$\Delta_i \ge 0 \quad \text{for } i \ge m,$$

with equality possible for at most one index. Consequently,

$$F[0,1], F[0,2], \ldots, F[0,d-1]$$

is non-increasing up to $F[0,m]$ and non-decreasing thereafter. □

---

**Proposition 2** For every $d \le 15$, every $1 \le i \le d-3$, and all branch lengths $w_1, \ldots, w_{d-2} > 0$, we have $R_{i+1} > R_i$.

*Proof* Let $\nabla_i := P_{i+1}Q_i - P_iQ_{i+1}$. Since $Q_i > 0$ for $1 \le i \le d-3$, cross multiplication shows that

$$R_{i+1} > R_i \Rightarrow P_{i+1}Q_i > P_iQ_{i+1} \Rightarrow \nabla_i := P_{i+1}Q_i - P_iQ_{i+1} > 0, \qquad 1 \le i \le d-3.$$

Thus, to show the sequence $R_i$ is strictly increasing, we only need to show that $\nabla_i > 0$. The boundary cases hold immediately:

$$\nabla_1 = P_2 Q_1 > 0,$$

because $P_1 = 0$, and

$$\nabla_{d-3} = P_{d-2} Q_{d-3} > 0,$$

because $Q_{d-2} = 0$.

We substitute $x_m := e^{-w_m} \in (0,1)$ in $\nabla_i$, which turns every exponential term into a monomial since $e^{-(w_k+\cdots+w_l)} = \prod_{m=k}^{l} x_m$. Treating $\nabla$ as a function of the unknown $x$, clearly $\nabla_i(x)$ becomes a multivariate polynomial in $x = (x_1, \ldots, x_{d-2})$ with rational coefficients. For any given $i$ and $d$, those coefficients can be calculated by carrying out the multiplications implied by:

$$\begin{aligned}
P_i &= (i-1) + \frac{1}{3}(i-1)\sum_{k=i+2}^{d-1} x_{i+1}\ldots x_{k-1} + \frac{2}{3}\sum_{l=1}^{i-1}(l-1)\,x_l\ldots x_{i-1}, \\
Q_i &= (d-2-i) + \frac{2}{3}\sum_{k=i+2}^{d-1}(d-1-k)\,x_{i+1}\cdots x_{k-1} + \frac{1}{3}(d-2-i)\sum_{l=1}^{i-1} x_l\cdots x_{i-1}. \\
\nabla_i &= P_{i+1}Q_i - P_i Q_{i+1}
\end{aligned} \tag{S6}$$

Such calculations would be cumbersome but can be automated easily.

Proving $\nabla_i > 0$ is equivalent to showing that the polynomial $\nabla_i(x)$ is positive on the box $x \in (0,1)^{d-2}$. It is not immediately clear whether $\nabla_i(x)$ is positive due to the presence of some negative coefficients from the minus operation in $P_{i+1}Q_i - P_i Q_{i+1}$. However, by applying the following Möbius transformation, we convert the positivity on a box $x \in (0,1)^{d-2}$ to positivity on the positive range $y \in (0,\infty)^{d-2}$, and therefore, prove $\nabla_i(y) > 0$ by showing that all coefficients of this transformation are positive. We perform the change of variables

$$x_m = \frac{1}{1+y_m}, \qquad y_m = e^{w_m} - 1,$$

which maps $(0,1)^{d-2}$ bijectively onto $(0,\infty)^{d-2}$. Let $\delta_m$ be the largest exponent of $x_m$ appearing in $\nabla_i(x)$; then, $\nabla_i(y)$ has a term with $\frac{1}{(1+y_m)^{\delta_m}}$. To remove these terms from the denominator and turn $\nabla_i(y)$ into a polynomial in $y$, we multiply $\nabla_i(y)$ by the largest exponent of each $x_m$:

$$N_i(y) = 9\nabla_i(y)\prod_{m=1}^{d-2}(1+y_m)^{\delta_m}. \tag{S7}$$

Since $y_m > 0$ for all $0 < m < d-1$, we can conclude $N_i(y) > 0$ if all its coefficients are positive. The strictly positive factor $9\prod_m(1+y_m)^{\delta_m}$ cancels all the denominators, so that $N_i(y)$ becomes a polynomial with integer coefficients and the same sign as $\nabla_i$. It therefore suffices to show that every coefficient of $N_i$ is positive. For any given $d$ and $i$, computing these integer coefficients is a finite computation over the integers, which can be performed automatically using a computer program using symbolic algebra libraries such as SymPy. Algorithm S1 constructs $N_i$ exactly (using rational and integer arithmetic only) and returns the smallest coefficient for any given $d$ and $i$. We implemented this procedure using help from Claude Opus (code is available at github.com/shayesteh99/BROOQS-Data/Scripts/ushape.py). The smallest coefficient $\mu(d) = \min_i \min_\alpha [y^\alpha -$ coefficient of $N_i]$ across all $1 \le i \le d-3$ is reported for every $d \le 20$ in Table S2; for $d \le 15$, this number is strictly positive.

Since every coefficient of $N_i$ is positive and every monomial is positive on $(0,\infty)^{d-2}$ for $d \le 15$, we conclude that $N_i(y) > 0$ throughout the orthant. Therefore, $\nabla_i > 0$ for $d \le 15$, completing the proof. □

**Table S2** The smallest coefficient $\mu(d) = \min_i \min_\alpha [y^\alpha\text{-coefficient of } N_i]$ of the polynomial defined in Equation (S7), computed exactly using Algorithm S1 over all $1 \le i \le d-3$. For $d \le 15$, every value is positive. For $d \ge 16$ cycles, the minimum can be negative. See Fig. S2 for an example of a 18-cycle where the hybrid F-scores are not unimodal and Section C.2 for empirical evaluation of $d \ge 16$.

| $d$ | 4 | 5 | 6 | 7 | 8 | 9 | 10 | 11 | 12 | 13 | 14 | 15 | 16 | 17 | 18 | 19 | 20 |
|---|---|---|---|---|---|---|---|---|---|---|---|---|---|---|---|---|---|
| $\mu(d)$ | 9 | 18 | 27 | 36 | 45 | 54 | 63 | 72 | 76 | 66 | 47 | 6 | -2032 | -15840 | -108612 | -387324 | -1206835 |

**Algorithm S1** MinCoefficient: Exact computation of the minimum coefficient of $N_i$. The implementation uses the symbolic algebra library SymPy Meurer et al. (2017) for polynomial construction, expansion, and coefficient extraction.

**function** MinCoefficient$(d, i)$
  Create symbolic variables $x_1, \ldots, x_{d-2}$ and $y_1, \ldots, y_{d-2}$, where $x_m = (1 + y_m)^{-1}$.
  **for** $s \in \{i, i+1\}$ **do**
    Construct $P_s$ and $Q_s$ from Equation (S6) as polynomials in $x = (x_1, \ldots, x_{d-2})$.
  **end for**
  Compute and expand $\nabla(x) = P_{i+1}Q_i - P_iQ_{i+1}$ as a polynomial in $x$.
  **for** $m = 1, \ldots, d-2$ **do**
    Let $\delta_m$ be the largest exponent of $x_m$ appearing in any monomial of $\nabla(x)$.
  **end for**
  Substitute $x_m = \frac{1}{(1+y_m)}$ in $\nabla$ and multiply by $\prod_{m=1}^{d-2}(1+y_m)^{\delta_m}$ to obtain

$$N = 9\nabla\left(\frac{1}{1+y_1}, \ldots, \frac{1}{1+y_{d-2}}\right) \prod_{m=1}^{d-2} (1+y_m)^{\delta_m}.$$

  Expand $N$ as a polynomial in $y = (y_1, \ldots, y_{d-2})$.
  Return the smallest coefficient among all monomials of $N$.
**end function**

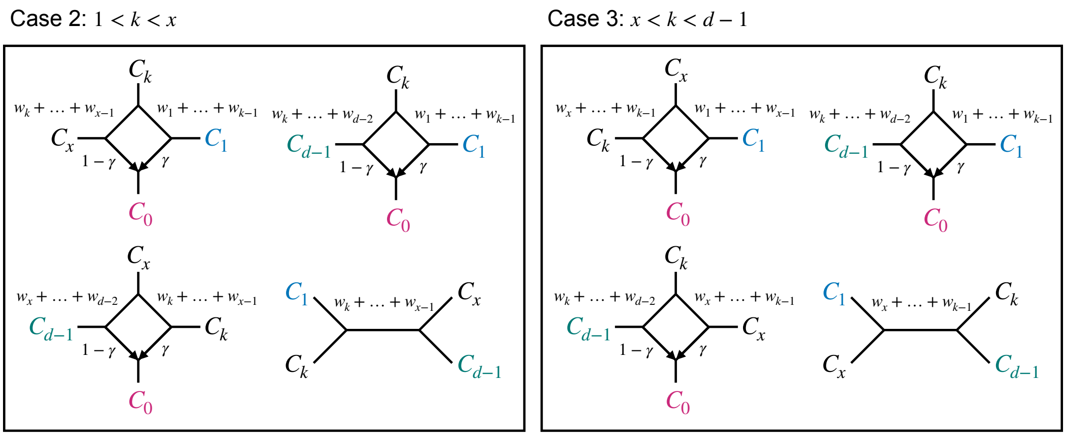


**Figure S21** All possible choices of four components containing $C_0$, a fixed component $C_x \notin \{C_1, C_{d-1}\}$, and $C_1$ and/or $C_{d-1}$. The corresponding metric quartet/quarnets are shown for each case.

#### B.2 Proof of Theorem 2

Recall:

**Theorem 2:** Let $\Omega$ be a $d$-cycle in $\mathcal{N}$, where $d > 4$, with true component ordering $\pi^*(\mathcal{C}) = [C_{\pi^*(0)}, C_{\pi^*(1)}, \ldots, C_{\pi^*(d-1)}]$, where $C_{\pi^*(0)}$ is the hybrid component and $C_{\pi^*(1)}$ and $C_{\pi^*(d-1)}$ are the neighboring components to $C_{\pi^*(0)}$. Then $F[\pi^*(1), \pi^*(d-1)] < F[\pi^*(i), \pi^*(j)]$ for every unordered pair $(i,j) \neq (1, d-1)$. Equivalently, $F[\pi^*(1), \pi^*(d-1)] = \min_{i \neq j} F[i,j]$, and this minimum is unique.

*Proof* We again use the simplified notation assuming $\pi^*(\mathcal{C}) = [C_0, C_1, \ldots, C_{d-1}]$, which makes $C_0$ the hybrid component, and makes $C_1$ and $C_{d-1}$ its neighboring components. Then, we can restate Theorem 2 as

$$F[1, d-1] = \min_{i \neq j} F[i,j].$$

We prove by contradiction. Suppose that $(x, y) \neq (1, d-1)$ satisfies

$$F[x, y] = \min_{i \neq j} F[i,j].$$

By Lemma 2, among all pairs of non-hybrid components,

$$F[1, d-1] = \min_{i,j \neq 0} F[i,j].$$

Therefore, at least one of $x$ or $y$ must be the hybrid component. Without loss of generality, let $y = 0$, and let $x = \arg\min_i F[0, i]$. By Remark 4, $\Delta_1 = F[0,2] - F[0,1] < 0$, therefore, $F[0,2] < F[0,1]$. Similarly, $F[0, d-2] < F[0, d-1]$. As a result, $x \notin \{1, d-1\}$. We show that $F[1, d-1] < F[0, x]$, which contradicts the choice of $(x, 0)$ as a minimizer. By definition,

$$F[0,x] = \sum_{\substack{k<l \\ k,l \notin \{0,x,1,d-1\}}} CF_{0,x|k,l} + \sum_{k \notin \{0,x,1,d-1\}} \left(CF_{0,x|k,1} + CF_{0,x|k,d-1}\right) + CF_{0,x|1,d-1},$$

and

$$F[1,d-1] = \sum_{\substack{k<l \\ k,l\notin\{0,x,1,d-1\}}} CF_{1,d-1|k,l} + \sum_{k\notin\{0,x,1,d-1\}} \left(CF_{1,d-1|k,0} + CF_{1,d-1|k,x}\right) + CF_{1,d-1|0,x}.$$

We compare each corresponding summand in the equations above and show that $F[1,d-1] < F[0,x]$ or $F[1,d-1] = F[0,x]$.

- **Case 1.** Let $k < l$, where $k,l \notin \{0,x,1,d-1\}$. By Lemma 3,

  $$CF_{0,x|k,l} = \gamma CF_{1,x|k,l} + (1-\gamma)CF_{d-1,x|k,l}.$$

  We have

  $$CF_{1,x|k,l} \geq CF_{1,d-1|k,l}$$

  and

  $$CF_{d-1,x|k,l} \geq CF_{1,d-1|k,l}.$$

  At least one of these inequalities hold strictly. If $k < l < x$, we have $CF_{d-1,x|k,l} > CF_{1,d-1|k,l}$; if $x < k < l$, we have $CF_{1,x|k,l} \geq CF_{1,d-1|k,l}$; and if $k < x < l$, both inequalities hold strictly. Therefore, we obtain

  $$CF_{0,x|k,l} > \gamma CF_{1,d-1|k,l} + (1-\gamma)CF_{1,d-1|k,l} = CF_{1,d-1|k,l}.$$

- **Case 2.** Let $1 < k < x$. The contribution of $k$ to $F[0,x]$ is

  $$CF_{0,x|k,1} + CF_{0,x|k,d-1}.$$

  Similarly, the contribution of $k$ to $F[1,d-1]$ is

  $$CF_{1,d-1|k,0} + CF_{1,d-1|k,x}.$$

  We compute both quantities and compare them:

  $$\begin{aligned} S_1 &= CF_{0,x|k,1} + CF_{0,x|k,d-1} \\ &= \gamma\left[\frac{1}{3}e^{-(w_1+\cdots+w_{k-1})}\right] + (1-\gamma)\left[1-\frac{2}{3}e^{-(w_k+\cdots+w_{x-1})}\right] + \gamma\left[\frac{1}{3}e^{-(w_k+\cdots+w_{x-1})}\right] + (1-\gamma)\left[\frac{1}{3}e^{-(w_x+\cdots+w_{d-2})}\right], \\ S_2 &= CF_{1,d-1|k,0} + CF_{1,d-1|k,x} \\ &= \gamma\left[\frac{1}{3}e^{-(w_1+\cdots+w_{k-1})}\right] + (1-\gamma)\left[\frac{1}{3}e^{-(w_k+\cdots+w_{d-2})}\right] + \frac{1}{3}e^{-(w_k+\cdots+w_{x-1})}. \end{aligned}$$

  Subtracting $S_2$ from $S_1$ gives

  $$S_1 - S_2 = (1-\gamma)\left[1 - e^{-(w_k+\cdots+w_{x-1})}\right] > 0.$$

  Therefore, $S_1 > S_2$.

- **Case 3.** Let $x < k < d-1$. The proof is symmetric to Case 2 (see Figure S21).

- **Case 4.** This case corresponds to the unique quarnet on $C_0, C_1, C_x, C_{d-1}$. In this case,

$$CF_{0,x|1,d-1} = CF_{1,d-1|0,x}.$$

The four cases above exhaust all summands in the definitions of $F[0,x]$ and $F[1,d-1]$. Case 4 contributes equally to both F-scores, whereas Cases 1–3 contribute strictly less to $F[1,d-1]$ than to $F[0,x]$. When $d=4$, only Case 4 is present, implying

$$F[0,x] = F[1,d-1] = \min_{i \neq j} F[i,j].$$

When $d > 4$, at least one summand from Cases 1–3 is present, so at least one inequality is strict. Therefore,

$$F[1,d-1] < F[0,x],$$

contradicting the choice of $(x,0)$ as a minimizer. Hence, $F[1,d-1]$ is the unique minimum whenever $d > 4$. □

##### B.3 Description of the PQ-tree algorithm of Armstrong et al. (2021)

The PQ-tree algorithm of Armstrong et al. (2021) takes a dissimilarity matrix $D \in \mathbb{R}^{d\times d}$ that is a permuted strict circular Robinson dissimilarity matrix, and returns a tree that represents all circular orderings consistent with $D$, up to rotation and reversal (Armstrong et al., 2021, Theorem 5.17). The algorithm relies on a data structure called a PQ-tree (Booth and Lueker, 1976), a rooted tree with two types of internal nodes: *P nodes*, whose children may be permuted freely, and *Q nodes*, whose children have a fixed order up to reversal. The algorithm only uses Q nodes; the leaves of each Q node always form a contiguous block (an arc) of the ordering.

The algorithm starts from $d$ trees, each consisting of a single leaf (Algorithm S2; line 3). At each step of the recursion, the algorithm first computes, for every pair of trees $T_1$ and $T_2$, the dissimilarity $D_{\min}[T_1, T_2] = \min_{x\in T_1, y\in T_2} D[x, y]$, as well as the set of pairs $D_{\arg\min}[T_1, T_2]$ that minimize this dissimilarity (line 6). For each $(x, y) \in D_{\arg\min}[T_1, T_2]$, since $x$ is the leaf of $T_1$ closest to $T_2$, it must sit at one end of $T_1$ in any consistent ordering. The algorithm therefore *orients* $T_1$ toward $x$: for each Q node in the path from the root of $T_1$ to $x$, it keeps or reverses their ordering so that $x$ becomes an end leaf of $T_1$ (similarly for $T_2$ and $y$; lines 8-9). Once a Q node is oriented, the order of its children can no longer change, so the node is removed and its children are attached directly to its parent node in the same order.

Next, the algorithm builds a *nearest-neighbor* graph, in which each tree is connected to its nearest tree(s) according to $D_{\min}$, and extracts its connected components (line 12). Since nearest neighbors must be adjacent in every ordering consistent with $D$ (Armstrong et al., 2021, Lemma 5.2), each connected component corresponds to a set of trees that are adjacent in *any* ordering consistent with $D$. The trees of each component are therefore merged into a new tree by attaching their roots, in path order of the nearest-neighbor graph, to a new Q node (line 21). The inner trees of the component, i.e., all except the first and last, are then oriented with respect to their two neighbors in the path, by comparing the dissimilarities between their border candidates and those of the neighbors (line 22). A tree whose orientation cannot be determined in this way is kept as a Q node, as both of its orientations are consistent with $D$. The two end trees of the component are oriented later, once the component is merged at a higher level of the recursion.

Since every tree is merged with at least its nearest neighbor, the number of trees at least halves at each step, and the recursion has depth at most $\log_2 d$. The algorithm stops when the nearest-neighbor graph has a single connected component (line 13). A final orientation is then performed on the root, whose children are treated as a ring, so that the first and last children are also neighbors (line 15). This fixes the ordering of the root's children up to rotation and reflection. In the resulting tree, at most one Q node remains that cannot be oriented (Armstrong et al., 2021, Lemma 5.19); reversing it yields the only other ordering consistent with $D$. The total time and memory complexity of this algorithm is $O(d^2)$.

**Algorithm S2** PQ-tree algorithm of Armstrong et al. (2021)

**function** PQtreeOrdering($F$) ▷ Alg. 5.2 in Armstrong et al. (2021)
    $D \leftarrow \binom{d-2}{2}(\mathbf{1}_{d\times d} - I_d) - F$
    $\mathcal{T} \leftarrow \{\{x\} : x \in [d]\}$ ▷ one single-leaf tree per element
    **while true do**
        **for all** ordered distinct pairs $(T_1, T_2)$ in $\mathcal{T}$ **do**
            $\big(D_{\min}[T_1, T_2],\, D_{\arg\min}[T_1, T_2]\big) \leftarrow \text{TreeDissimilarity}(T_1, T_2, D)$
            **for all** $(x, y) \in D_{\arg\min}[T_1, T_2]$ **do**
                ExternalOrientation($T_1, x$) ▷ Alg. 5.7 in Armstrong et al. (2021)
                ExternalOrientation($T_2, y$)
            **end for**
        **end for**
        $\mathcal{C} \leftarrow \text{ArcPartition}(D_{\min}, \mathcal{T})$ ▷ Alg. B.1 in Armstrong et al. (2021)
        **if** $|\mathcal{C}| = 1$ **then**
            $R \leftarrow$ new Q node whose children are the trees of the component, in path order
            FinalOrientation($R$) ▷ Alg. 5.6 in Armstrong et al. (2021)
            $\Pi \leftarrow$ the set of orderings corresponding to $R$
            **return** $\pi = \arg\max_{\pi\in\Pi} \sum_{i=0}^{d-1} F[\pi(i), \pi(i+1 \mod d)]$
        **end if**
        $\mathcal{T}' \leftarrow \emptyset$
        **for all** components $(T_1, \ldots, T_k) \in \mathcal{C}$ **do**
            $T \leftarrow$ new Q node with children $T_1, \ldots, T_k$
            CompleteInternalOrientation($T$) ▷ Alg. 5.5 in Armstrong et al. (2021)
            $\mathcal{T}' \leftarrow \mathcal{T}' \cup \{T\}$
        **end for**
        $\mathcal{T} \leftarrow \mathcal{T}'$
    **end while**
**end function**

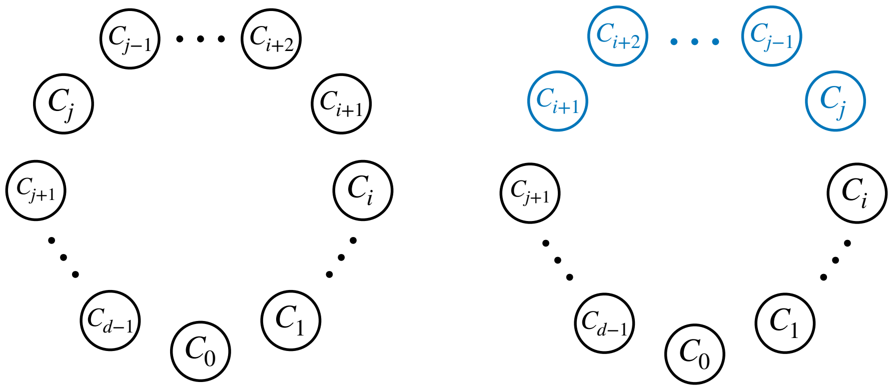

**Figure S22** The true ordering $\pi^*(\mathcal{C})$ (left) corresponding to the cycle in Fig. S17, and a partial reversal of it (right). The sequence $[C_{i+1}, C_{i+2}, \ldots, C_{j-1}, C_j]$ is reversed in the partial reversal.

#### B.4 Proof of correctness for PQ-tree algorithm

**Definition 5** Let $\pi = [\pi(0), \pi(1), \ldots, \pi(d-1)]$ be a circular ordering on $d$ elements. We say an ordering $\pi' \neq \pi$ is a *partial reversal* of $\pi$, if there exist two indices $0 \leq i < j \leq d-1$, such that $j > i+1$ and $i \neq (j+1) \mod d$ and $\pi' = [\pi(0), \ldots, \pi(i), \pi(j), \pi(j-1), \ldots, \pi(i+1), \pi(j+1), \ldots, \pi(d-1)]$. Informally, $\pi'$ is obtained by reversing the sequence $[\pi(i+1), \ldots, \pi(j)]$ in $\pi$. Note that this ordering is equivalent to reversing the sequence $[\pi(j+1), \ldots, \pi(d-1), \pi(0), \ldots, \pi(i)]$ in $\pi$, and therefore, the constraint $i < j$ covers all the partial reversals of $\pi$.

**Lemma 1:** Let $\Omega$ be a $d$-cycle where $4 \leq d \leq 15$ with components $\mathcal{C}$ and true cyclic ordering $\pi^*(\mathcal{C})$ with permuted F-matrix $\pi^*(F)$. Then, the PQ-tree algorithm of Armstrong et al. (2021) on the dissimilarity matrix $D = \binom{d-2}{2}(\mathbf{1}_{d\times d} - I_d) - F$ returns at most two inequivalent orderings $\Pi = \{\pi_1, \pi_2\}$, such that $\pi^*(\mathcal{C}) = \arg\max_{\pi\in\Pi} \sum_{i=0}^{d-1} F[\pi(i), \pi(i+1 \mod d)]$.

*Proof* By Theorem 1, $\pi^*(F)$ is a strict Robinson similarity matrix. It follows immediately that $D$ is a strict Robinson *dissimilarity* matrix; that is, elements in every row and column of $D$ strictly increase up to one or two consecutive maximum(s) and then strictly decrease. By Lemma 5.19 of Armstrong et al. (2021), PQ-tree algorithm returns at most two inequivalent orderings $\Pi = \{\pi_1, \pi_2\}$ up to rotation and reversal. By their Theorem 5.17, $\pi^*(\mathcal{C}) \in \Pi$, and by Lemma 5.19, the other ordering is a partial reversal of $\pi^*(\mathcal{C})$. In Lemma 7, we prove that under NMSC, for any partial reversal $\pi$ of the true ordering $\pi^*(\mathcal{C})$,

$$\sum_{i=0}^{d-1} F[\pi(i), \pi(i+1 \mod d)] < \sum_{i=0}^{d-1} F[\pi^*(i), \pi^*(i+1 \mod d)].$$

Therefore,

$$\pi^*(\mathcal{C}) = \arg\max_{\pi\in\Pi} \sum_{i=0}^{d-1} F[\pi(i), \pi(i+1 \mod d)].$$

□

**Lemma 7** For a $d-$cycle $\Omega$ with components $\mathcal{C}$ and the true ordering $\pi^*(\mathcal{C})$, let $\pi$ be a partial reversal of $\pi^*(\mathcal{C})$. Then,

$$\sum_{i=0}^{d-1} F[\pi(i), \pi(i+1 \mod d)] < \sum_{i=0}^{d-1} F[\pi^*(i), \pi^*(i+1 \mod d)].$$

*Proof* For simplicity of notation, let $\pi^*(\mathcal{C}) = [C_0, C_1, \ldots, C_{d-1}]$, where $C_0$ is the hybrid component (Fig. S17). Let $\pi$ be a partial reversal of $\pi^*(\mathcal{C})$ at positions $i$ and $j$, obtained by reversing the sequence $[C_{i+1}, \ldots, C_j]$ in $\pi^*(\mathcal{C})$ (Fig. S22). Let

$$\Delta(\pi, \pi^*(\mathcal{C})) = \sum_{a=0}^{d-1} F[\pi^*(a), \pi^*(a+1 \mod d)] - F[\pi(a), \pi(a+1 \mod d)].$$

Therefore, we need to show $\Delta > 0$. Because $F$ is symmetric, for two elements adjacent in both $\pi^*(\mathcal{C})$ and $\pi$, they contribute 0 to $\Delta$. Out of the $d$ terms in the summation, only two terms are not equal, which correspond to the two sets of components not adjacent in both orderings. Therefore,

$$\Delta = F[i, i+1] + F[j, j+1] - F[i, j] - F[i+1, j+1].$$

Since $i \neq (j+1) \mod d$ (by Definition 5), $i = 0$ and $j = d-1$ is not a valid partial reversal. Also, as noted in Definition 5, each reversal is equivalent to the reversal of the complement of the elements reversed. Thus, we can arbitrarily choose element 0 to stay in the non-reversed part, and designate the segment excluding 0 as the reversed. Therefore, we need to show $\Delta > 0$ for the two cases of $i > 0, j < d-1$ and $i = 0, j < d-1$. The proof of $j = d-1, i > 0$ follows from the latter case by symmetry.

**The case of $i > 0$ and $j < d-1$ (no hybrid or neighbors):**

Since all $i, i+1, j, j+1$ are non-hybrid components, by Lemma 2, we have:

$$F[i, i+1] > F[i, i+2] > \ldots > F[i, j],$$

and

$$F[j+1, j] > F[j+1, j-1] > \ldots > F[j+1, i+1].$$

Therefore, $\Delta = F[i, i+1] + F[j, j+1] - F[i, j] - F[i+1, j+1] > 0$.

**The case of $i = 0$ and $j < d-1$ (neighbor included in reversal):**

We have $\Delta = F[0, 1] + F[j, j+1] - F[0, j] - F[1, j+1]$. We prove $\Delta > 0$ for every $2 \leq j < d-1$. We expand each term:

$$F[0,1] = \sum_{k<l;k,l \notin \{0,1,j,j+1\}} CF_{0,1|k,l} + \sum_{k \notin \{0,1,j,j+1\}} CF_{0,1|j,k} + \sum_{k \notin \{0,1,j,j+1\}} CF_{0,1|j+1,k} + CF_{0,1|j,j+1},$$

$$F[j,j+1] = \sum_{k<l;k,l \notin \{0,1,j,j+1\}} CF_{j,j+1|k,l} + \sum_{k \notin \{0,1,j,j+1\}} CF_{j,j+1|0,k} + \sum_{k \notin \{0,1,j,j+1\}} CF_{j,j+1|1,k} + CF_{j,j+1|0,1},$$

$$F[0,j] = \sum_{k<l;k,l \notin \{0,1,j,j+1\}} CF_{0,j|k,l} + \sum_{k \notin \{0,1,j,j+1\}} CF_{0,j|1,k} + \sum_{k \notin \{0,1,j,j+1\}} CF_{0,j|k,j+1} + CF_{0,j|1,j+1},$$

$$F[1,j+1] = \sum_{k<l;k,l \notin \{0,1,j,j+1\}} CF_{1,j+1|k,l} + \sum_{k \notin \{0,1,i,j+1\}} CF_{1,j+1|0,k} + \sum_{k \notin \{0,1,j,j+1\}} CF_{1,j+1|k,j} + CF_{1,j+1|0,j}.$$

For each choice of four components, we compute its contribution to $F[0,1], F[j,j+1], F[0,j], F[1,j+1]$ and therefore, $\Delta$. Figure S23 shows all possible cases.

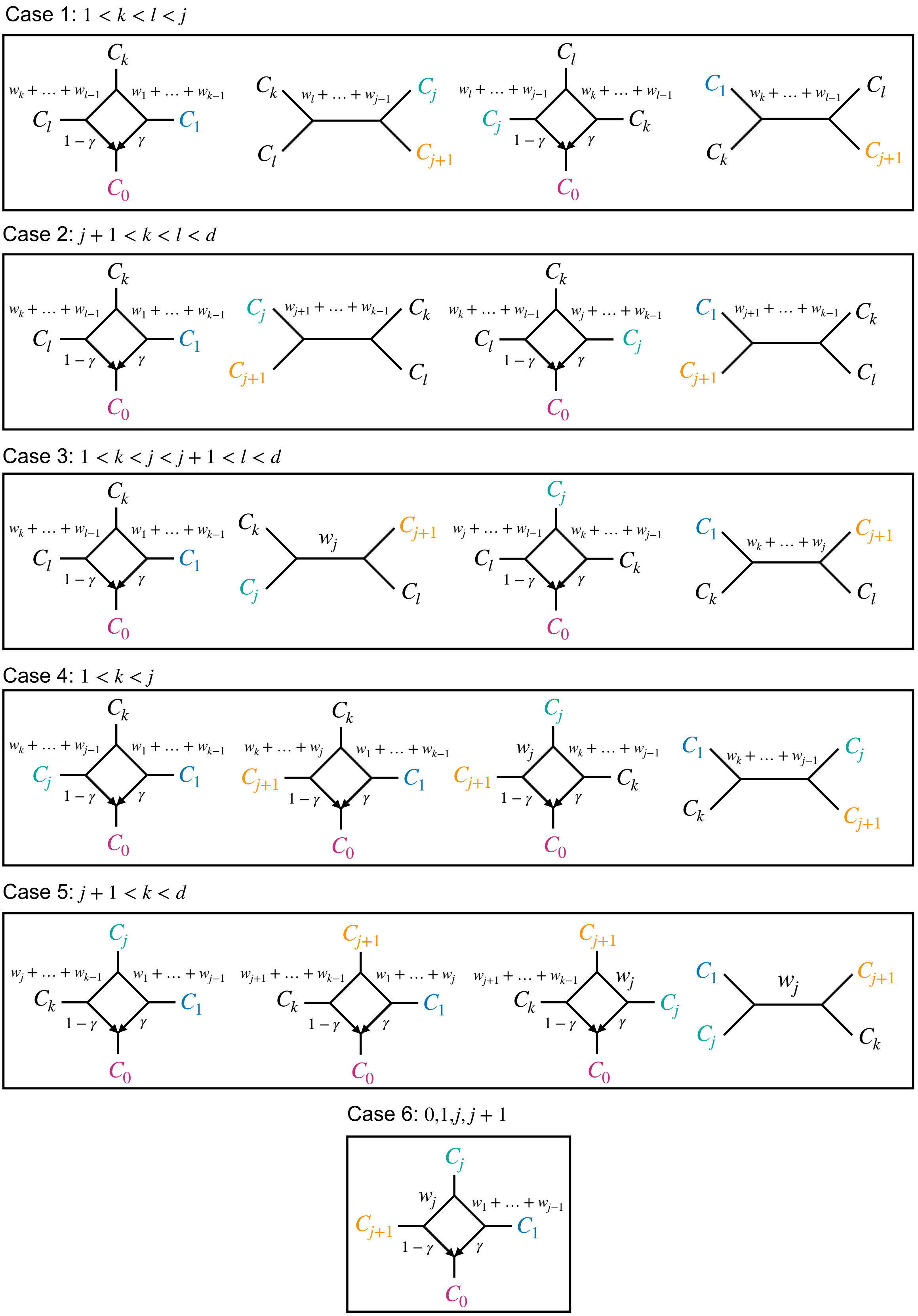


**Figure S23** All possible choices of four components containing any of the pairs $C_0$ and $C_1$, $C_j$ and $C_{j+1}$, $C_0$ and $C_j$, or $C_1$ and $C_{j+1}$. The corresponding metric quartet/quarnets are shown for each case in the true order.

- **Case 1.** Suppose $1 < k < l < j$. For any choice of $k, l$ we have

$$
\begin{aligned}
CF_{0,1|k,l} &= \gamma[1 - \frac{2}{3}e^{-(w_1+\cdots+w_{k-1})}] + (1-\gamma)[\frac{1}{3}e^{-(w_k+\cdots+w_{l-1})}] \\
CF_{j,j+1|k,l} &= 1 - \frac{2}{3}e^{-(w_l+\cdots+w_{j-1})} \\
CF_{0,j|k,l} &= \gamma[\frac{1}{3}e^{-(w_k+\cdots+w_{l-1})}] + (1-\gamma)[1 - \frac{2}{3}e^{-(w_l+\cdots+w_{j-1})}] \\
CF_{1,j+1|k,l} &= \frac{1}{3}e^{-(w_k+\cdots+w_{l-1})}.
\end{aligned}
$$

Therefore,

$$
\begin{aligned}
\Delta &= CF_{0,1|k,l} + CF_{j,j+1|k,l} - CF_{0,j|k,l} - CF_{1,j+1|k,l} \\
&= \gamma[1 - \frac{2}{3}e^{-(w_1+\cdots+w_{k-1})}] - 2\gamma[\frac{1}{3}e^{-(w_k+\cdots+w_{l-1})}] + \gamma[1 - \frac{2}{3}e^{-(w_l+\cdots+w_{j-1})}]
\end{aligned} \tag{S8}
$$

Since $1 - \frac{2}{3}e^{-(w_1+\cdots+w_{k-1})} > \frac{1}{3}$ and $1 - \frac{2}{3}e^{-(w_l+\cdots+w_{j-1})} > \frac{1}{3}$ and $\frac{1}{3}e^{-(w_k+\cdots+w_{l-1})} < \frac{1}{3}$, we have,

$$\Delta = CF_{0,1|k,l} + CF_{j,j+1|k,l} - CF_{0,j|k,l} - CF_{1,j+1|k,l} > 0.$$

- **Case 2.** Suppose $j + 1 < k < l < d$. For any choice of $k, l$ we have

$$
\begin{aligned}
CF_{0,1|k,l} &= \gamma[1 - \frac{2}{3}e^{-(w_1+\cdots+w_{k-1})}] + (1-\gamma)[\frac{1}{3}e^{-(w_k+\cdots+w_{l-1})}] \\
CF_{j,j+1|k,l} &= 1 - \frac{2}{3}e^{-(w_{j+1}+\cdots+w_{k-1})} \\
CF_{0,j|k,l} &= \gamma[1 - \frac{2}{3}e^{-(w_j+\cdots+w_{k-1})}] + (1-\gamma)[\frac{1}{3}e^{-(w_k+\cdots+w_{l-1})}] \\
CF_{1,j+1|k,l} &= 1 - \frac{2}{3}e^{-(w_{j+1}+\cdots+w_{k-1})}.
\end{aligned}
$$

Therefore,

$$\Delta = CF_{0,1|k,l} + CF_{j,j+1|k,l} - CF_{0,j|k,l} - CF_{1,j+1|k,l} = \gamma[1 - \frac{2}{3}e^{-(w_1+\cdots+w_{k-1})}] - \gamma[1 - \frac{2}{3}e^{-(w_j+\cdots+w_{k-1})}]$$

Since $w_1 + \cdots + w_{k-1} > w_j + \cdots + w_{k-1}$, therefore, $1 - \frac{2}{3}e^{-(w_1+\cdots+w_{k-1})} > 1 - \frac{2}{3}e^{-(w_j+\cdots+w_{k-1})}$ and,

$$\Delta = CF_{0,1|k,l} + CF_{j,j+1|k,l} - CF_{0,j|k,l} - CF_{1,j+1|k,l} > 0.$$

- **Case 3.** Suppose $1 < k < j$ and $j + 1 < l < d$. For any choice of $k, l$ we have

$$
\begin{aligned}
CF_{0,1|k,l} &= \gamma[1 - \frac{2}{3}e^{-(w_1+\cdots+w_{k-1})}] + (1-\gamma)[\frac{1}{3}e^{-(w_k+\cdots+w_{l-1})}] \\
CF_{j,j+1|k,l} &= \frac{1}{3}e^{-w_j} \\
CF_{0,j|k,l} &= \gamma[\frac{1}{3}e^{-(w_k+\cdots+w_{j-1})}] + (1-\gamma)[\frac{1}{3}e^{-(w_j+\cdots+w_{l-1})}] \\
CF_{1,j+1|k,l} &= \frac{1}{3}e^{-(w_k+\cdots+w_j)}.
\end{aligned}
$$

Let $u_1 = e^{-(w_1+\cdots+w_{k-1})}$, $u_2 = e^{-(w_k+\cdots+w_{j-1})}$, $u_3 = e^{-w_j}$ and $u_4 = e^{-(w_{j+1}+\cdots+w_{l-1})}$, so $u_1,\ldots,u_4 \in (0,1)$. Therefore,

$$\begin{aligned}
\Delta &= CF_{0,1|k,l} + CF_{j,j+1|k,l} - CF_{0,j|k,l} - CF_{1,j+1|k,l} \\
&= \gamma(1-\frac{2}{3}u_1) + (1-\gamma)(\frac{1}{3}u_2u_3u_4) + \frac{1}{3}u_3 - \gamma\frac{1}{3}u_2 - (1-\gamma)\frac{1}{3}u_3u_4 - \frac{1}{3}u_2u_3 \\
&= \gamma(1-\frac{2}{3}u_1 - \frac{1}{3}u_2) - (1-\gamma)\frac{1}{3}u_3u_4(1-u_2) + \frac{1}{3}u_3(1-u_2) \\
&= \gamma(1-\frac{2}{3}u_1 - \frac{1}{3}u_2) + \frac{1}{3}u_3(1-u_2)(1-(1-\gamma)u_4)
\end{aligned}$$

Since each term is easily shown to be positive, therefore,

$$\Delta = CF_{0,1|k,l} + CF_{j,j+1|k,l} - CF_{0,j|k,l} - CF_{1,j+1|k,l} > 0.$$

- **Case 4.** Suppose $1 < k < j$. For any choice of $k$ we have

$$\begin{aligned}
CF_{0,1|k,l} &= \gamma[1-\frac{2}{3}e^{-(w_1+\cdots+w_{k-1})}] + (1-\gamma)[\frac{1}{3}e^{-(w_k+\cdots+w_{j-1})}] + \gamma[1-\frac{2}{3}e^{-(w_1+\cdots+w_{k-1})}] + (1-\gamma)[\frac{1}{3}e^{-(w_k+\cdots+w_j)}] \\
CF_{j,j+1|k,l} &= \gamma[1-\frac{2}{3}e^{-(w_k+\cdots+w_{j-1})}] + (1-\gamma)\frac{1}{3}e^{-w_j} + 1 - \frac{2}{3}e^{-(w_k+\cdots+w_{j-1})} \\
CF_{0,j|k,l} &= \gamma[\frac{1}{3}e^{-(w_1+\cdots+w_{k-1})}] + (1-\gamma)[1-\frac{2}{3}e^{-(w_k+\cdots+w_{j-1})}] + \gamma[\frac{1}{3}e^{-(w_k+\cdots+w_{j-1})}] + (1-\gamma)\frac{1}{3}e^{-w_j} \\
CF_{1,j+1|k,l} &= \gamma[\frac{1}{3}e^{-(w_1+\cdots+w_{k-1})}] + (1-\gamma)[\frac{1}{3}e^{-(w_k+\cdots+w_j)}] + \frac{1}{3}e^{-(w_k+\cdots+w_{j-1})}.
\end{aligned}$$

Therefore,

$$\begin{aligned}
\Delta &= CF_{0,1|k,l} + CF_{j,j+1|k,l} - CF_{0,j|k,l} - CF_{1,j+1|k,l} \\
&= 2\gamma[1-e^{-(w_1+\cdots+w_{k-1})}] + 2\gamma[1-\frac{2}{3}e^{-(w_k+\cdots+w_{j-1})}] - 2\gamma\frac{1}{3}e^{-(w_k+\cdots+w_{j-1})} \\
&= 2\gamma[1-e^{-(w_1+\cdots+w_{k-1})}] + 2\gamma[1-e^{-(w_k+\cdots+w_{j-1})}].
\end{aligned}$$

Therefore,

$$\Delta = CF_{0,1|k,l} + CF_{j,j+1|k,l} - CF_{0,j|k,l} - CF_{1,j+1|k,l} > 0.$$

- **Case 5.** Suppose $j+1 < k < d$. For any choice of $k$ we have

$$\begin{aligned}
CF_{0,1|k,l} &= \gamma[1-\frac{2}{3}e^{-(w_1+\cdots+w_{j-1})}] + (1-\gamma)[\frac{1}{3}e^{-(w_j+\cdots+w_{k-1})}] + \gamma[1-\frac{2}{3}e^{-(w_1+\cdots+w_j)}] + (1-\gamma)[\frac{1}{3}e^{-(w_{j+1}+\cdots+w_{k-1})}] \\
CF_{j,j+1|k,l} &= \gamma\frac{1}{3}e^{-w_j} + (1-\gamma)[1-\frac{2}{3}e^{-(w_{j+1}+\cdots+w_{k-1})}] + \frac{1}{3}e^{-w_j} \\
CF_{0,j|k,l} &= \gamma[\frac{1}{3}e^{-(w_1+\cdots+w_{j-1})}] + (1-\gamma)[\frac{1}{3}e^{-(w_j+\cdots+w_{k-1})}] + \gamma[1-\frac{2}{3}e^{-w_j}] + (1-\gamma)[\frac{1}{3}e^{-(w_{j+1}+\cdots+w_{k-1})}] \\
CF_{1,j+1|k,l} &= \gamma[\frac{1}{3}e^{-(w_1+\cdots+w_j)}] + (1-\gamma)[1-\frac{2}{3}e^{-(w_{j+1}+\cdots+w_{k-1})}] + \frac{1}{3}e^{-w_j}.
\end{aligned}$$

Therefore,

$$\begin{aligned}
\Delta &= CF_{0,1|k,l} + CF_{j,j+1|k,l} - CF_{0,j|k,l} - CF_{1,j+1|k,l} \\
&= \gamma[1-e^{-(w_1+\cdots+w_{j-1})}] + \gamma[1-e^{-(w_1+\cdots+w_j)}] - \gamma[1-e^{-w_j}].
\end{aligned}$$

Since $w_1 + \cdots + w_j > w_j$, therefore $[1 - e^{-(w_1+\cdots+w_j)}] > [1 - e^{-w_j}]$ and,

$$\Delta = CF_{0,1|k,l} + CF_{j,j+1|k,l} - CF_{0,j|k,l} - CF_{1,j+1|k,l} > 0.$$

- **Case 6.** Consider the quarnet on $C_0, C_1, C_j, C_{j+1}$. Based on the quarnet concordance factors, we have:

$$CF_{0,j|1,j+1} = CF_{1,j+1|0,j} < CF_{0,1|j,j+1} = CF_{j,j+1|0,1}.$$

Therefore,

$$\Delta = CF_{0,1|k,l} + CF_{j,j+1|k,l} - CF_{0,j|k,l} - CF_{1,j+1|k,l} > 0.$$

The above cases constitute all the summands in $\Delta = F[i, i+1] + F[j, j+1] - F[i, j] - F[i+1, j+1]$, and together they prove that,

$$\sum_{i=0}^{d-1} F[\pi(i), \pi(i+1 \mod d)] < \sum_{i=0}^{d-1} F[\pi^*(i), \pi^*(i+1 \mod d)]$$

$\square$

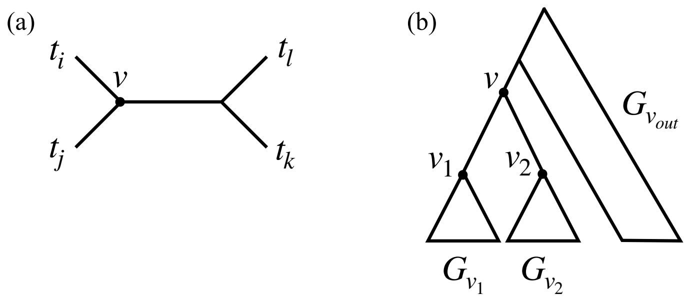


**Figure S24** (a) Each quartet $q = t_i t_j | t_k t_l$ in $G$ is counted exactly once at the node $v$ where $v$ is the anchor node that induces the tripartition $\{t_i\}, \{t_j\}, \{t_k, t_l\}$. (b) Removing each node $v \in V_G$ and its incident edges creates three subtrees, $G_{v_1}$, $G_{v_2}$, and $G_{v_{out}}$, where $v_1$ and $v_2$ are the two children of $v$.

#### B.5 Description of the algorithm used to compute $\widehat{F}$

Combining Equation (1) and Equation (2), we obtain

$$\widehat{F}[i,j] = \sum_{\substack{k<l \\ k,l \neq i,j}} \frac{1}{|\mathcal{G}|} \sum_{G \in \mathcal{G}} \sum_{t_i \in C_i} \sum_{t_j \in C_j} \sum_{t_k \in C_k} \sum_{t_l \in C_l} \frac{\mathbf{1}\Big[G|_{\{t_i,t_j,t_k,t_l\}} = t_i t_j | t_k t_l\Big]}{|C_i||C_j||C_k||C_l|} \tag{S9}$$

Assume that every gene tree in $\mathcal{G}$ contains at least one taxon from every component in $\mathcal{C}$. Under this assumption, the order of summation over gene trees and component pairs can be exchanged, giving

$$\widehat{F}[i,j] = \frac{1}{|\mathcal{G}|} \sum_{G \in \mathcal{G}} \sum_{\substack{k<l \\ k,l \neq i,j}} \sum_{t_i \in C_i} \sum_{t_j \in C_j} \sum_{t_k \in C_k} \sum_{t_l \in C_l} \frac{\mathbf{1}\Big[G|_{\{t_i,t_j,t_k,t_l\}} = t_i t_j | t_k t_l\Big]}{|C_i||C_j||C_k||C_l|}$$

The interchange of these summations is valid only under the assumption above. In Section B.5.1, we describe the heuristics used when one or more components are absent from a gene tree.

Let

$$\widehat{F}[i,j][G] = \sum_{\substack{k<l \\ k,l \neq i,j}} \sum_{t_i \in C_i} \sum_{t_j \in C_j} \sum_{t_k \in C_k} \sum_{t_l \in C_l} \frac{\mathbf{1}\Big[G|_{\{t_i,t_j,t_k,t_l\}} = t_i t_j | t_k t_l\Big]}{|C_i||C_j||C_k||C_l|}$$

denote the contribution of a single gene tree $G$ to the empirical F-score between components $C_i$ and $C_j$ so that

$$\widehat{F}[i,j] = \frac{1}{|\mathcal{G}|} \sum_{G \in \mathcal{G}} \widehat{F}[i,j][G]$$

Our efficient algorithm computes $\widehat{F}[i,j][G]$ for a single gene tree $G$ and a pair of components $C_i$ and $C_j$ in a single traversal of $G$, resulting in $O(n)$ running time for one pair of components and $O(d^2 n)$ time for all pairs.

For each internal node $v \in V_G$, we count the number of quartets of the form $t_i t_j | t_k t_l$ for which $v$ is the anchor node that induces the tripartition $\{t_i\}, \{t_j\}, \{t_k, t_l\}$ on the quartet (Figure S24a). Assigning every quartet to its anchor node ensures that each quartet is counted exactly once.

Let $G_{v_1}$, $G_{v_2}$, and $G_{v_{\text{out}}}$ denote the three subtrees obtained by removing $v$ and its incident edges from $G$ (Figure S24b), and let $L_{v_1}$, $L_{v_2}$, and $L_{v_{\text{out}}}$ denote their corresponding leaf sets. Similar to the dynamic programming approach used in ASTRAL Mirarab et al. (2014), we use

$$\widehat{F}[i,j][G] = \sum_{v\in V_G} \sum_{\substack{L_X,L_Y,L_Z\\ \in\{L_{v_1},L_{v_2},L_{v_{\text{out}}}\}}} \overbrace{\frac{|L_X\cap C_i|}{|C_i|}}^{I_{X,i}} \overbrace{\frac{|L_Y\cap C_j|}{|C_j|}}^{I_{Y,j}} \overbrace{\sum_{\substack{k<l\\ k,l\neq i,j}} \frac{|L_Z\cap C_k|}{|C_k|}\frac{|L_Z\cap C_l|}{|C_l|}}^{I_{Z,\bar{i},\bar{j}}}. \tag{S10}$$

The equation above can be evaluated in $O(n)$ time for one gene tree. This is possible due to a preprocessing step (Algorithm S3) that takes $O(dn)$ time and precomputes a collection of quantities for every node $v$ and every component $C_i$, including $I_{v,i}$, which can be computed with a trivial sum as we move up the tree (see Algorithm S3). For the $I_{Z,\bar{i},\bar{j}}$ term, we can calculate it in constant time for each pair $(C_i, C_j)$ using

$$I_{Z,\bar{i},\bar{j}} = \sum\nolimits_{\substack{k<l\\ k,l\neq i,j}} \tfrac{|L_Z\cap C_k|}{|C_k|}\tfrac{|L_Z\cap C_l|}{|C_l|} = \sum\nolimits_{\substack{k<l\\ k,l\neq i,j}} I_{Z,k} I_{Z,l} = \tag{S11}$$

$$\overbrace{\sum_m \sum_{n<m} I_{Z,m} I_{Z,n}}^{T_Z} - \overbrace{I_{Z,i} \sum_{m\neq i} I_{Z,m}}^{D_{Z,i}} - \overbrace{I_{Z,j} \sum_{m\neq j} I_{Z,m}}^{D_{Z,j}} + I_{Z,i} I_{Z,j} = \tag{S12}$$

$$\overbrace{\sum_m \frac{1}{2} I_{Z,m} \sum_{n\neq m} I_{Z,n}}^{T_Z} \overbrace{- I_{Z,i}(-I_{Z,i} + \overbrace{\sum_m I_{Z,m}}^{N_Z})}^{D_{Z,i}} - D_{Z,j} + I_{Z,i} I_{Z,j} = \tag{S13}$$

$$\overbrace{\sum_m \frac{1}{2} D_{Z,m}}^{T_Z} - \overbrace{I_{Z,i}(N_Z - I_{Z,i})}^{D_{Z,i}} - D_{Z,j} + I_{Z,i} I_{Z,j}. \tag{S14}$$

This computation requires only two additional quantities for each node $v$: the total sum over all component pairs ($T_v$) and the sum involving a fixed component ($D_{v,i}$). Both quantities are computed during preprocessing in $O(dn)$ time, as shown in Algorithm S3, using simple recursions shown in Equations (S13) and (S14). Care must be taken for the $v_{out}$ group, as shown in the Algorithm.

Once preprocessing is completed for a gene tree $G$, all pairwise F-scores can be computed in $O(n)$ time each using Algorithm S4.

**Algorithm S3** PREPROCESSGENETREE

**function** PREPROCESSGENETREE$(G, \phi)$
$root \leftarrow$ root of $G$
Initialize all entries of $N$, $I$, $D$, $T$, $D'$, and $T'$ to $0$
**for all** nodes $v$ in postorder traversal of $V_G$ **do**
**if** $v$ is a leaf **then**
$i \leftarrow \phi(v)$ ▷ $\phi(v)$ is the mapping of each leaf in $L_G$ to its corresponding component $C_i$
$I[v,i] \leftarrow \frac{1}{|C_i|}$ ▷ Defined in Equation (S10)
$N[v] \leftarrow I[v,i]$ ▷ Defined in Equation (S13)
**else**
**for all** children $u$ of $v$ **do**
$N[v] \leftarrow N[v] + N[u]$ ▷ See Equation (S13)
**for** $i = 1$ to $d$ **do**
$I[v,i] \leftarrow I[v,i] + I[u,i]$ ▷ See Equation (S10)
**end for**
**end for**
**end if**
**end for**
**for all** nodes $v$ in postorder **do**
**for** $i = 1$ to $d$ **do**
$D[v,i] \leftarrow I[v,i]\big(N[v] - I[v,i]\big)$ ▷ See Equation (S14)
$T[v] \leftarrow T[v] + \frac{1}{2}\, D[v,i]$ ▷ See Equation (S14)
$D'[v,i] \leftarrow \big(I[root,i] - I[v,i]\big)\Big(N[root] - I[root,i] - \big(N[v] - I[v,i]\big)\Big)$ ▷ Outgroup partition slightly different.
$T'[v] \leftarrow T'[v] + \frac{1}{2}\, D'[v,i]$ ▷ Similar to $T$ but for the outgroup clade
**end for**
**end for**
**return** $N, I, D, T, D', T'$
**end function**

**Algorithm S4** COMPUTEGENETREEFSCORE

**function** COMPUTEGENETREEFSCORE$(G, C_i, C_j)$
Retrieve the preprocessing tables from Algorithm S3
$root \leftarrow$ root of $G$
$F \leftarrow 0$
**for all** internal nodes $v$ **do**
$O_i \leftarrow I[root,i] - I[v,i]$
$O_j \leftarrow I[root,j] - I[v,j]$
$O_{\neq ij} \leftarrow T'[v] - D'[v,i] - D'[v,j] + O_i\, O_j$
**for all** pairs of distinct children $(v_1, v_2)$ of $v$ **do**
$F \leftarrow F + I[v_1,i] \cdot I[v_2,j] \cdot O_{\neq ij}$
$F \leftarrow F + I[v_1,i] \cdot \Big(T[v_2] - D[v_2,i] - D[v_2,j] + I[v_2,i] \cdot I[v_2,j]\Big) \cdot O_j$
$F \leftarrow F + I[v_1,j] \cdot \Big(T[v_2] - D[v_2,i] - D[v_2,j] + I[v_2,i] \cdot I[v_2,j]\Big) \cdot O_i$
**end for**
**end for**
**return** $F$
**end function**

##### B.5.1 Handling missing taxa and missing components

In the formulation given above, missing taxa from a component can be organically handled as long as the entire component is not missing. To handle these cases, for every gene tree $G$, we simply replace $|C_i|$ by $|L_G \cap C_i|$ for every component in Equation (S10). Thus, each gene tree with at least one taxon from each component contributes equally to the final $F$ score.

When entire components are missing from gene trees, Equation (S9) should ideally be rewritten as

$$\widehat{F}[i,j] = \sum_{\substack{k<l \\ k,l \neq i,j}} \frac{1}{|\mathcal{G}_{C_i,C_j,C_k,C_l}|} \sum_{G \in \mathcal{G}_{C_i,C_j,C_k,C_l}} \sum_{t_i \in C_i} \sum_{t_j \in C_j} \sum_{t_k \in C_k} \sum_{t_l \in C_l} \frac{\mathbf{1}\left[G|_{\{t_i,t_j,t_k,t_l\}} = t_i t_j | t_k t_l\right]}{\prod_{x \in \{i,j,k,l\}} |L_G \cap C_x|}, \tag{S15}$$

where $\mathcal{G}_{C_i,C_j,C_k,C_l}$ denotes the set of gene trees that contain at least one taxon from each of the four components $C_i$, $C_j$, $C_k$, and $C_l$.

A direct implementation of Equation (S15) requires storing the number of gene trees for every combination of four components, requiring $O(d^4)$ memory and increasing the total running time of our algorithm from $O(d^2nm)$ to $O(d^4nm)$ for $m$ gene trees. To maintain the scalability of BROOQS, we instead use the following heuristic.

Assume a random subsampling that removes taxa from complete gene trees. We make one assumption about this process: Whether a component $C_i$ appears at least once in any gene tree is independent of whether another component $C_j$ appears in it. We need not assume independence of missing taxa across loci, nor uniformly random subsampling, as long as this single assumption holds. Many complex sampling processes could have this property, including the very simple model where each taxon is removed independently from others with a fixed probability from each gene tree.

Let $\mathcal{G}_{C_i}$ denote the set of gene trees containing at least one taxon from component $C_i$ and note

$$\mathbb{E}[|\mathcal{G}_{C_i}|] = |\mathcal{G}| \Pr(C_i \cap L_G \neq \emptyset)$$

Under the assumption we made,

$$\begin{aligned} \mathbb{E}[|\mathcal{G}_{C_i,C_j,C_k,C_l}|] =& |\mathcal{G}| \Pr(C_i \cap L_G \neq \emptyset \wedge C_j \cap L_G \neq \emptyset \wedge C_l \cap L_G \neq \emptyset \wedge C_k \cap L_G \neq \emptyset) = \\ & |\mathcal{G}| \prod_{x \in \{i,j,k,l\}} \Pr(C_x \cap L_G \neq \emptyset) = \\ & |\mathcal{G}| \frac{\mathbb{E}[|\mathcal{G}_{C_i}|]}{|\mathcal{G}|} \frac{\mathbb{E}[|\mathcal{G}_{C_j}|]}{|\mathcal{G}|} \frac{\mathbb{E}[|\mathcal{G}_{C_k}|]}{|\mathcal{G}|} \frac{\mathbb{E}[|\mathcal{G}_{C_l}|]}{|\mathcal{G}|} = \mathbb{E}\left[|\mathcal{G}| \prod_{x \in \{i,j,k,l\}} \frac{|\mathcal{G}_{C_x}|}{|\mathcal{G}|}\right] \end{aligned}$$

We use this calculation of expected value to devise an approximation. In Equation (S15), we replace $|\mathcal{G}_{C_i,C_j,C_k,C_l}|$, which is calculable from the data but slow to compute, with

$$|\mathcal{G}_{C_i,C_j,C_k,C_l}| \approx |\mathcal{G}| \prod_{x \in \{i,j,k,l\}} \frac{|\mathcal{G}_{C_x}|}{|\mathcal{G}|}$$

noting that the two terms are equal in expectation under our assumptions. Then,

$$\widehat{F}[i,j] \approx \sum_{\substack{k<l \\ k,l \neq i,j}} \frac{1}{|\mathcal{G}|} \sum_{G \in \mathcal{G}} \sum_{t_i \in C_i} \sum_{t_j \in C_j} \sum_{t_k \in C_k} \sum_{t_l \in C_l} \frac{\mathbf{1}\left[G|_{\{t_i,t_j,t_k,t_l\}} = t_i t_j | t_k t_l\right]}{\prod_{x \in \{i,j,k,l\}} \frac{|L_G \cap C_x| \cdot |\mathcal{G}_{C_x}|}{|\mathcal{G}|}}$$

These modified weights can be incorporated directly into Algorithm S3 by replacing the weight $\frac{1}{|C_i|}$ (line 7) with

$$\frac{|\mathcal{G}|}{|L_G \cap C_i| \cdot |\mathcal{G}_{C_i}|}.$$

Computing $|\mathcal{G}_{C_i}|$ for all components requires only a single pass over the gene trees, taking $O(md)$ time. Therefore, the overall asymptotic running time of our algorithm remains unchanged. Our simulation analyses show that this heuristic is robust to increasing levels of missing data (Fig. S4).

### C Supplementary Text

#### C.1 Evaluation Metrics

**Kendal–Tau error**

The definition of Kendal–Tau error on linear orderings can be generalized to cyclic orderings by considering triplets (choices of three components) instead of pairs. Given a true cyclic ordering $\pi^*(\mathcal{C})$ and an estimated ordering $\pi(\mathcal{C})$ on a $d-$cycle, total number of triplets are $\binom{d}{3}$. Let $t$ be the number of disagreeing triplets between the true and estimated orderings. Since $\mathrm{Rev}(\pi(\mathcal{C}))$ is equivalent to $\pi(\mathcal{C})$, the Kendal–Tau error between $\pi^*(\mathcal{C})$ and $\pi(\mathcal{C})$ is computed as:

$$\mathrm{KT}(\pi^*(\mathcal{C}), \pi(\mathcal{C})) = \frac{2\min(t, \binom{d}{3} - t)}{\binom{d}{3}}.$$

Note that the maximum possible value of $\min(t, \binom{d}{3} - t)$ is $\frac{\binom{d}{3}}{2}$, hence the multiplier of 2 in the numerator ensures that $\mathrm{KT}(\pi^*(\mathcal{C}), \pi(\mathcal{C})) \in [0, 1]$ for any two orderings on the same set of components.

**Minimum normalized Robinson–Foulds distance (min nRF)**

Let $\mathcal{D}_{\mathrm{true}}$ be the set of displayed trees of the true network $\mathcal{N}_{\mathrm{true}}$. For each $T_i \in \mathcal{D}_{\mathrm{true}}$, let $w_i$ be the product of the inheritance probabilities of all hybrid edges in $T_i$. For an estimated network $\mathcal{N}_{\mathrm{est}}$ with displayed tree set $\mathcal{D}_{\mathrm{est}}$, we define the minimum normalized Robinson–Foulds distance (min nRF) as

$$\min \mathrm{nRF}(\mathcal{N}_{\mathrm{true}}, \mathcal{N}_{\mathrm{est}}) = \sum_{T_i \in \mathcal{D}_{\mathrm{true}}} w_i \min_{T' \in \mathcal{D}_{\mathrm{est}}} RF(T_i, T'),$$

where $RF(T_1, T_2)$ is the normalized Robinson–Foulds distance between the two trees. Note that $\sum_i w_i = 1$, and therefore, $\min \mathrm{nRF}(\mathcal{N}_{\mathrm{true}}, \mathcal{N}_{\mathrm{est}}) \in [0, 1]$ for any two networks on the same set of taxa.

#### C.2 Empirical examination

Appendix Section B.1 proved (Theorem 1) that for any cycle $\Omega$ of size $4 \leq d \leq 15$, the F-matrix $F$ constructed on $\Omega$ is a strict circular Robinson similarity matrix. Together with Lemma 1, we prove that the PQ-tree algorithm identifies the true cyclic ordering for all cycles of size $4 \leq d \leq 15$. For larger cycles, Theorem 1 does not hold for all choices of parameters (as shown in the counterexample in Fig. S2). We explore this violation and its impact on recovering the true cyclic ordering in extensive simulations with extreme choices of parameters. Moreover, Conjecture 1 states that the spectral algorithm of Coifman et al. (2008) recovers the true ordering of any cycle $\Omega$ of size $4 \leq d \leq 19$. This conjecture is backed by the extensive simulations summarized in Table S3.

Table S3 shows four model conditions based on cycle size:

- $d$ ranges from $[4, 15]$, the regime where both Theorem 1 and Conjecture 1 hold.
- $[16, 19]$, the regime where Theorem 1 fails and Conjecture 1 holds.
- $[20, 25]$, a regime where both Theorem 1 and Conjecture 1 fail.

We simulate $10^6$ replicates for each model condition and each range of $d$, and report the number of replicates where Theorem 1 holds, and the PQ-tree and the spectral method recover the true ordering. We observe no violations of Theorem 1 for $d \in [4, 15]$ and Conjecture 1 for $d \in [4, 19]$.

Each model condition captures an extreme choice of parameters to evaluate whether each condition holds at the borders of the parameter space. The four model conditions are: extreme $\gamma$ where $\gamma$ is drawn uniformly from $(0.001, 0.05) \cup (0.95, 0.999)$, bimodal branch lengths where each branch length $w_i$ is drawn from a log-uniform distribution of $[10^{-6}, 10^{-3}]$ or $[3, 15]$ with probability $0.5$, extreme $\gamma$ and bimodal branch lengths, and a more extreme scenario, referred to as the lopsided blocks, where the non-hybrid components of the cycle are divided into a large block adjacent to the hybrid component (with size randomly drawn from $[\lceil (d-1)/2 \rceil, d-3]$), and a set of smaller blocks (with sizes randomly drawn from $[1, 4]$). Within each block, the branch lengths are drawn from a log-uniform distribution of $[10^{-6}, 10^{-3}]$ and the branch lengths separating blocks are drawn from a log-uniform distribution of $[3, 15]$. The inheritance probability of the hybrid edge incident to the large block of components is set to be much larger than the other hybrid edge, which is drawn from a log-uniform distribution of $[10^{-7}, 10^{-2}]$. We chose the parameters for this model condition carefully to replicate the known scenarios where the spectral algorithm does not recover the true ordering for $d \geq 20$, while preventing failures due to numerical underflow.

**Table S3** Validation of the strict circular Robinson property of $F$ and recovery of the true cyclic ordering using PQ-tree algorithm and the spectral method.

| Condition | $d$ | Branch lengths $w_i$ | Inheritance prob. $\gamma$ | $F$ strictly circ. Robinson | Spectral recovers | PQ-tree recovers |
|---|---|---|---|---|---|---|
| Extreme $\gamma$ | 4–15 | $w_i \sim U(0.3, 3)$ | $\gamma \sim U(0.001, 0.05)$ $\cup U(0.95, 0.999)$ | $10^6/10^6$ | $10^6/10^6$ | $10^6/10^6$ |
| | 16–19 | | | $10^6/10^6$ | $10^6/10^6$ | $10^6/10^6$ |
| | 20–25 | | | $10^6/10^6$ | $10^6/10^6$ | $10^6/10^6$ |
| Bimodal $w$ | 4–15 | $w_i \sim \text{LogU}(10^{-6}, 10^{-3})$ $\vee \text{LogU}(3, 15)$ | $\gamma \sim U(0.001, 0.999)$ | $10^6/10^6$ | $10^6/10^6$ | $10^6/10^6$ |
| | 16–19 | | | $996,246/10^6$ | $10^6/10^6$ | $10^6/10^6$ |
| | 20–25 | | | $972,792/10^6$ | $10^6/10^6$ | $10^6/10^6$ |
| Extreme $\gamma$ + Bimodal $w$ | 4–15 | $w_i \sim \text{LogU}(10^{-6}, 10^{-3})$ $\vee \text{LogU}(3, 15)$ | $\gamma \sim U(0.001, 0.05)$ $\cup U(0.95, 0.999)$ | $10^6/10^6$ | $10^6/10^6$ | $10^6/10^6$ |
| | 16–19 | | | $10^6/10^6$ | $10^6/10^6$ | $10^6/10^6$ |
| | 20–25 | | | $10^6/10^6$ | $10^6/10^6$ | $10^6/10^6$ |
| Lopsided blocks* | 4–15 | bimodal* $w_i$ | $1-\gamma$ or $\gamma \sim \text{LogU}(10^{-7}, 10^{-2})$ | $10^6/10^6$ | $10^6/10^6$ | $10^6/10^6$ |
| | 16–19 | | | $10^6/10^6$ | $10^6/10^6$ | $10^6/10^6$ |
| | 20–25 | | | $10^6/10^6$ | 973,496/$10^6$ † | $10^6/10^6$ |

* One block of size $U\{\lceil (d-1)/2 \rceil, \ldots, d-3\}$ plus blocks of size 1–4; within each block edges are drawn from $\text{LogU}(10^{-6}, 10^{-3})$, between-block edges are drawn from $\text{LogU}(3, 15)$. † All 26,504 failures have strictly circular Robinson $F$. The number of failed cases by $d$ are $961$ for $d = 20$, $647$ for $d = 21$, $2,613$ for $d = 22$, $4,208$ for $d = 23$, $7,650$ for $d = 24$, and $10,425$ for $d = 25$.
